# Duplication and neofunctionalization of a horizontally-transferred xyloglucanase as a facet of the red queen co-evolutionary dynamic

**DOI:** 10.1101/2023.10.09.561229

**Authors:** Victoria Attah, David S Milner, Yufeng Fang, Xia Yan, Guy Leonard, Joseph Heitman, Nicholas J. Talbot, Thomas A Richards

**Affiliations:** Department of Biology, University of Oxford, Oxford, United Kingdom; Department of Molecular Genetics and Microbiology, Duke University Medical Center, Durham, North Carolina, United States of America; The Sainsbury Laboratory, University of East Anglia, Norwich Research Park, United Kingdom; GreenLight Biosciences Inc., Research Triangle Park, North Carolina, United States of America

**Keywords:** Horizontal Gene Transfer, Gene Duplication, Host Pathogen Interaction, Plant Cell Wall

## Abstract

Oomycetes are heterotrophic protists that share phenotypic similarities with fungi, including the ability to cause plant diseases, but branch in a separate and distant region of the eukaryotic tree of life. It has been suggested that multiple horizontal gene transfers (HGTs) from fungi-to-oomycetes contributed to the evolution of plant-pathogenic traits. These HGTs are predicted to include secreted proteins that degrade plant cell walls. This is a key trait in the pathology of many oomycetes, as the plant cell wall represents a primary barrier to pathogen invasion and a rich source of carbohydrates. Many of the HGT gene families identified have undergone multiple rounds of duplication. Using a combination of phylogenomic analysis and functional assays, we investigate the diversification of a horizontally-transferred xyloglucanase gene family in the model oomycete species *Phytophthora sojae*. Our analyses detect 11 genes retained in *P. sojae* among a complex pattern of gene duplications and losses. Using a phenotype assay, based on heterologous expression in yeast, we show that eight of these paralogs have xyloglucanase function, including variants with distinct protein characteristics, such as a long-disordered C-terminal extension that can increase xyloglucanase activity. The functional xyloglucanase variants analysed subtend an ancestral node close to the fungi-oomycetes gene transfer, suggesting the horizontally-transferred gene was a *bona fide* xyloglucanase. Expression of xyloglucanase paralogs in *Nicotiana benthamiana* triggers distinct patterns of reactive oxygen species (ROS) generation, demonstrating that enzyme variants differentially stimulate pattern-triggered immunity in plants. Mass spectrometry of detectable enzymatic products demonstrates that some paralogs catalyze production of variant breakdown profiles, suggesting that secretion of multiple xyloglucanase variants increases efficiency of xyloglucan breakdown, as well as potentially diversifying the range of Damage-Associated Molecular Patterns (DAMPs) released during pathogen attack. We suggest that such patterns of protein neofunctionalization, and variant host responses, represent an aspect of the Red Queen host-pathogen co-evolutionary dynamic.

**Significance Statement:** The oomycetes are a diverse group of eukaryotic microbes that include some of the most devastating pathogens of plants. Oomycetes perceive, invade, and colonize plants in similar ways to fungi, in part because they acquired the genes to attack and feed on plants from fungi. These genes are predicted to be useful to oomycete plant pathogens because they have undergone multiple rounds of gene duplication. One key enzyme for attacking plant cell wall structures is called xyloglucanase. Xyloglucanase in the oomycetes has undergone multiple rounds of gene duplication, leading to variants including an enzyme with a C-terminal extension that increases activity. Some xyloglucanase variants trigger unique patterns of reactive oxygen species (ROS) *in planta*, and generate different profiles of cell wall breakdown products - such outcomes could act to mystify and increase the workload of the plant immune system, allowing successful pathogens to proliferate.

## Introduction

Oomycetes are heterotrophic protists that form part of the stramenopile lineage (also sometimes called heterokonts) (1–5). They resemble fungi in their filamentous growth and osmotrophic feeding (6), which initially led to their taxonomic placement within the kingdom Fungi (7). It is now known that oomycetes and fungi branch on distant parts of the eukaryotic tree of life (8–13), suggesting that biological similarities of the two groups were the product of convergent evolution (14, 15). The oomycetes are a highly diverse group of protists, but include ecologically- and agriculturally-destructive pathogens of plants (16). Many plant pathogens penetrate or break down the cell walls of their hosts, a heterogenous refractory structure rich in diverse polysaccharides. Plant cell walls are composed of cellulose cross-linked by glycans, and embedded within a matrix of pectin, hemicelluloses, lignin, other aromatic polymers, and proteins, which together provide structural integrity (17, 18). One of the most abundant hemicellulosic polysaccharides is xyloglucan, consisting of a backbone of D-glucose linked by β-1-4 glycosidic bonds (similar to cellulose), however, most are substituted with α-1,6-linked xylose (often in turn capped with galactose, arabinose, or fucose). Owing to the different possible variations of the sugar side chains on xylosyl residues, the xyloglucan structure varies significantly between plant species (19–23). A nomenclature-based system has been used to define side chain variants and describe the oligosaccharides released by enzymatic degradation of xyloglucan (24). Interestingly, mixed xyloglucan oligosaccharides (obtained through enzymatic extraction from apple pomace) have also been demonstrated to act as Damage-Associated Molecular Patterns (DAMPs) in *Vitis vinifera* (grapevine) and *Arabidopsis thaliana*, leading to direct activation of a range of plant immune responses across phylogenetically-distant plant species (25).

The genomes of many plant pathogens, especially hemibiotrophic species, are replete in genes encoding secreted glycoside hydrolases, predicted to break down external plant cell wall layers (26, 27). These gene families include previously identified horizontal gene transfers (HGTs) from fungi-to-oomycetes (28–34). HGT, or lateral gene transfer (LGT), describes the movement of genes across species boundaries, a process which can potentially add new functionality to the recipient lineage (e.g. (35, 36)). New functionality can also be acquired through gene duplication events– specifically when the gene duplication is coupled with neofunctionalization of sister paralogs. In oomycetes there is evidence of widespread gene duplication (post-acquisition) of fungal-derived HGT genes (31, 37), resulting in multiple paralogs, many of which are predicted to encode secreted products which interact with a host. This pattern of evolution seems to be prevalent among economically-important *Phytophthora* species, including *Phytophthora sojae* – a hemibiotrophic pathogen of soybean (31, 38, 39).

We know very little regarding the functional significance of HGT events and subsequent gene family duplications in *Phytophthora*. Gene duplication events can be linked to subsequent adaptation regarding increased gene dosage and/or functional divergence known as neofunctionalization (40) or subfunctionalisation (41, 42). Previous work has demonstrated that one paralog of the *P. sojae* Glycoside Hydrolase 12 (GH12) gene family, PsXLP1, has undergone a significant pattern of deletion within the encoded open reading frame (ORF), rendering a catalytically-inactivated protein that acts as a ‘decoy’ of plant immune response, helping to facilitate infection (43). The PsXLP1 gene is located in a conserved head-to-head orientation with the closely-related paralogous PsXEG1 gene, and has been shown to inhibit the host immune protein GmGIP1, whereas none of the other GH12 members tested were shown to inhibit the same host protein (43). Additional work has also focused on detailed examination of the functional PsXEG1 paralog and its contribution to *P. sojae* virulence (44). These results demonstrated that neofunctionalization through peptide sequence evolution has been adaptive with regard to *P. sojae* virulence and immunity evasion.

Here, we characterize all 11 HGT derived paralogs [xenologues] that have resulted from gene duplication in the same HGT-derived *P. sojae* GH12 gene family, using bioinformatics, heterologous protein expression in *Saccharomyces cerevisiae* and transient expression in *N. benthamiana*. Our analysis suggests a pattern of functional divergence linked to unique protein characteristics important for xyloglucanase function. We also show that some paralogous functional forms render different xyloglucan breakdown product profiles, and suggest that secretion of multiple variants could therefore increase the efficacy of substrate breakdown. Importantly, we demonstrate that the variant paralogs trigger unique patterns of reactive oxygen species (ROS) accumulation *in planta*, suggesting that the host must be on high alert to multiple related protein signatures during pathogen interaction. The concordant action of multiple paralogs transcribed during infection may therefore act to perturb plant immune perception both within individual infections (e.g. via differential transcription), and over evolutionary divergence (e.g. by replacement of variant paralog functions). Such a scenario would suggest that the pattern of gene loss, duplication and neofunctionalization observed here is an evolutionary consequence of the Red Queen co-evolutionary dynamic (45, 46), which predicts that host-pathogen co-evolution acts to increase genetic variation by generating diversity on molecular signatures involved in the interaction (47).

## Results

The aim of this work was to investigate the functional consequences of a series of gene duplications of a horizontally-transferred, putatively secreted, enzyme using a bioinformatics approach, combined with heterologous expression in *S. cerevisiae* to allow for functional assessment of enzymatic function. The enzyme is a putative xyloglucanase, and this gene family was previously identified as a candidate HGT from fungi to the oomycetes primarily based on taxon distribution data (29, 31, 33). In this study we investigated the hypothesis that gene duplication events which occurred after the HGT event, drove diversification and neofunctionalization of this enzyme function in *P. sojae.* To better understand the dynamics of GH12 gene turnover in oomycetes, we explored patterns of lineage-specific gene gains (e.g. via duplication and fixation) or losses (e.g. via inactivation and deletion) across representative *Phytophthora* spp., and *Hyaloperonospora arabidopsidis* (an obligate biotrophic pathogen of *Arabidopsis thaliana*) taxa (Figure 1A). A total of 11 paralogs of GH12 in *P. sojae* were confirmed by HMM searches against Ensembl genomes (48–50). The *P. sojae* GH12 paralogs were aligned to a range of candidate GH12 genes identified in additional oomycete genomes sampled using the same HMM search method described above but excluding some sequences (see Figure 1A). We then constructed a phylogenetic tree to explore the specific evolutionary relationships between *P. sojae* GH12 paralogs, using focused sampling of orthologs in closely related oomycetes, demonstrating the presence of 11 *P. sojae* genes across multiple paralog clades (Figure 1). We note that this analysis shows weak phylogenetic resolution in several sections of the tree (i.e. bootstrap support values) and clear evidence of multiple cases of gene loss both within the 9 clades identified (Figure 1A), and unspecified gene losses among the intermediate phylogenetic groups which lack resolution. Even with many nodes lacking strong support, these data demonstrate that the gene family has been subject to numerous cases of gene duplication and loss typical of strong diversifying selection operating on the GH12 gene repertoire and putatively contributing to functional variation in this lineage important for pathogenesis.

**Figure 1.**
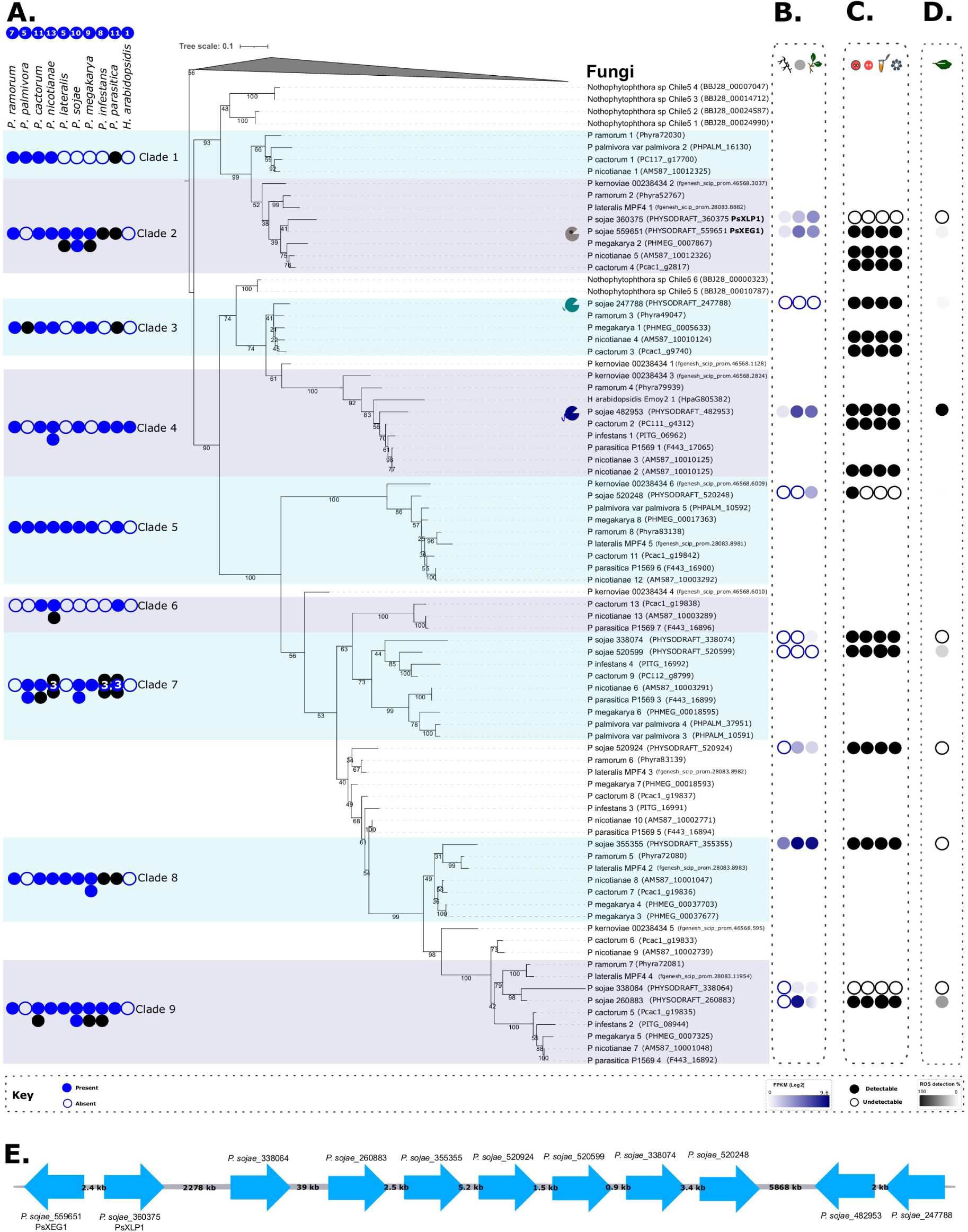
Evolutionary history of oomycete GH12. **A.** Maximum likelihood tree constructed with IQ-Tree v2.0.3 (WAG+R5 model of evolution (87) selected as the best-fit model by ModelFinder (74)). The tree was constructed from an alignment of 161 sequences comprising 211 amino acids to confirm the evolutionary relationships between oomycete GH12. The final tree was visualized with iTOL (76), and nodes indicate results (%) of non-parametric bootstrap (200 pseudoreplicates) (75). The tree is rooted with a fungal outgroup because fungi were previously identified as the putative donor group of the GH12 HGT into the oomycetes (31, 33). The eleven *P. sojae* GH12 paralogs are visually identifiable as they have adjacent transcriptome profiles (column B). Symbols indicate the *P. sojae* paralog with a structurally-inferred additional binding site (grey), and paralogs with C-terminal tails 1 and 2 (blue and green, respectively). The numbers of GH12 orthologs across representative *Phytophthora* spp., and *Hyaloperonospora arabidopsidis* (an obligate pathogen of *Arabidopsis thaliana*) genomes are shown to the left of the phylogenetic tree, indicating lineage-specific putative local duplications or losses. Some are coloured black, indicating they are absent in the tree due to exclusion because of gappy gene models, or they formed uninformative branches (sometimes long-branches sometimes very short), potentially affecting bootstrap resolution. All *P. sojae* paralogs were retained. **B.** *P. sojae* life-cycle transcriptome data (FungiDB: (51, 52)) were used to identify how the eleven *P. sojae* GH12 paralogs were expressed across (column 1) mycelial, (column 2) cyst and (column 3) 3 days post-infection (soybean hypocotyls infected with *P. sojae* strain P6497) – expressed in FPKM(Log2). **C.** Xyloglucanase functional data generated during this study from (column 1) cell culture agar plate-based enzyme assay, (column 2) concentrated supernatant agar-plate based enzyme assay, (column 3) concentrated supernatant DNS reducing sugar assay, and (column 4) concentrated supernatant mass spectrometry of xyloglucan breakdown products (black fill = function detected, white fill = no function detected). **D.** ROS data in response to *P. sojae* xyloglucanase variants, presented as percentage ROS generation (%) of the variant that triggered the highest ROS accumulation in *N. benthamiana* (i.e. *P. sojae*_482953, which is presented as 100%). **E.** FungiDB (51, 52) was used to locate approximate genomic coordinates of each *P. sojae* GH12 gene - the relative approximate distances between the paralogs are shown in kilobases (kb), providing additional support for the sister paralog relationships in the phylogenetic tree for: *P. sojae*_559651 PsXEG1 and *P. sojae*_360375 PsXLP1 (PHYSO_scaffold_4); *P. sojae_*247788 and *P. sojae_*482953 (PHYSO_scaffold_2).

FungiDB (51, 52) was used to locate approximate genomic coordinates of each *P. sojae* GH12 gene - the relative approximate distances between the paralogs are shown in kilobases (kb) (Figure 1E). These results demonstrate two phylogenetically-defined sister paralog pairs, which also demonstrate close clustering on chromosomal contigs (*P. sojae*_559651 (PsXEG1) and *P. sojae*_360375 (PsXLP1); *P. sojae_*247788 and *P. sojae_*482953). Close chromosomal positions can be used as additional evidence to support sister-paralog relationships identified in phylogenetic trees.

We searched *P. sojae* life-cycle transcriptome datasets (FungiDB (51, 52)) to identify and compare how the eleven *P. sojae* GH12 paralogs were expressed across mycelial, cyst and 3 days post-infection (soybean hypocotyls infected with *P. sojae* strain P6497). Analysis of these transcriptomes demonstrated that the duplicated sequences exhibit stage-specific expression, as well as differences in levels of expression (Fragments Per Kilobase of exon model per Million mapped reads (FPKM) were used to calculate Log2 values (Figure 1B)). Four paralogs are expressed in *P. sojae* mycelia, seven paralogs are expressed in cyst, and nine paralogs are expressed during plant infection. Four of the eleven xyloglucanases are expressed to some level during all three stages: *P. sojae*_360375 (PsXLP1), *P. sojae*_559651 (PsXEG1), *P. sojae*_482953, and *P. sojae*_355355 (Figure 1B). Gene expression of two of the eleven xyloglucanases was undetectable in all stages: *P. sojae*_247788 and *P. sojae*_520599 (Figure 1B). We do not include it in Figure 1B, but we note that previous analyses describe fine-scale correlated expression patterns for *P. sojae*_360375 (PsXLP1) and *P. sojae*_559651 (PsXEG1), which are consistent with their expression during all stages as described above (43, 44). The authors demonstrate that both variants are expressed at low levels in mycelia, cyst, germinating cyst and zoospore stages, with an increase in expression at 10 minutes post-infection of soybean hypocotyls, up to the highest peak in expression at 1-hour post-infection, before declining (43).

Alignment analysis demonstrated the predicted amino acid sequences of *P. sojae*_482953 and *P. sojae*_247788 had an extended C-terminus compared to the other nine paralogs, while *P. sojae*_559651 (PsXEG1) has two single amino acid insertions not found in other *P. sojae* GH12 paralogs (Supplemental Figure 1). This analysis also confirmed that *P. sojae*_360375 (PsXLP1) has a curtailed protein sequence in comparison to the other eleven *P. sojae* paralogs, as previously documented (43).

### Heterologous expression of *P. sojae* GH12 paralogs in *S. cerevisiae* demonstrates that eight enzymes exhibit activity against xyloglucan

All *P. sojae* GH12 gene models were manually checked using publically-available RNAseq data for *P. sojae* (51, 52). Confirmed candidate nucleotide sequences for all eleven *P. sojae* GH12 paralogs were synthesized, codon-optimized for *S. cerevisiae*, and cloned into plasmid p426-GPD for expression in *S. cerevisiae*. To maximize heterologous protein secretion in yeast, *P. sojae* N-terminal signal peptide sequences were replaced with the native *S. cerevisiae* mating factor α (MFA) signal peptide sequences (53). Concentrated supernatants of yeast strains, each expressing a *P. sojae* GH12 paralog, were used as crude protein extracts for enzyme activity assays using 3,5-Dinitrosalicylic acid (DNS) reagent compared to a yeast strain carrying an empty vector. DNS is an aromatic compound that is reduced in the presence of reducing sugars (released during the breakdown of polysaccharides), to 3-amino-5-nitrosalicylic acid, which absorbs light at ∼540 nm (54). Therefore, an increase in absorbance at 544 nm over time was used to infer enzymatic activity.

After 6 hours incubation with 1% (w/v) xyloglucan (pH 7, 30°C), significant levels of reducing sugars released by enzyme activity were detected for the *P. sojae*_482953, 559651 (PsXEG1), 520924, and 338074 xyloglucanase variants (Dunnett’s test; Figure 2A). After prolonged incubation (72 h), significant levels of reducing sugars were also detected for the

**Figure 2.**
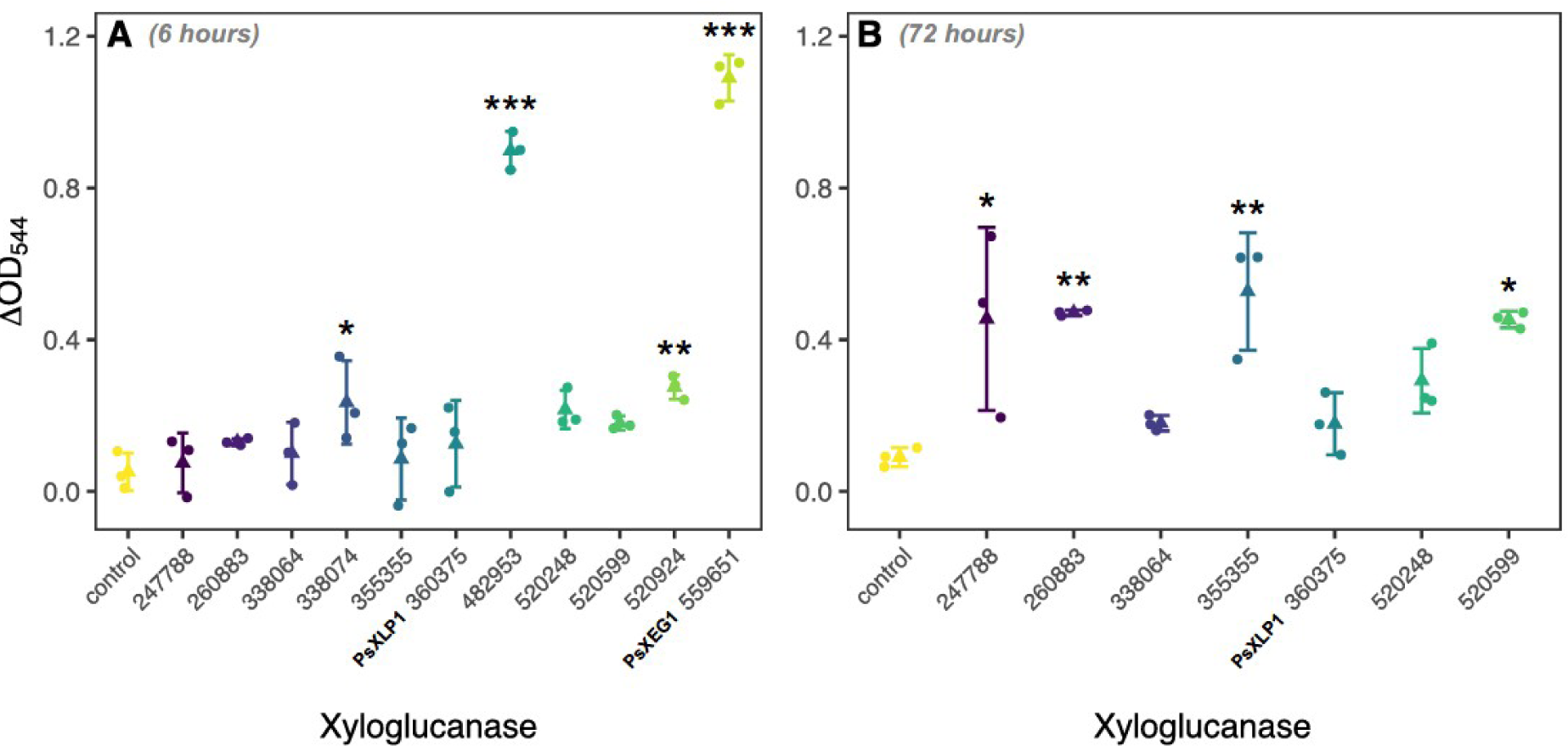
Comparative assay of xyloglucanase function across the 11 *P. sojae* GH12 paralogs. *P. sojae* GH12 paralogs secreted into *S. cerevisiae* culture supernatants were incubated with 1% (w/v) xyloglucan at 30°C, pH7; an increase in absorbance (OD_544_) of DNS reagent added to the samples is suggestive of an increase in the reducing sugars released (i.e. from the breakdown of the substrate). **A.** Significant enzymatic activity towards xyloglucan was detected up to 6 hours of incubation for *P. sojae*_338074, 482953, 520924 and 559651 xyloglucanase variants (Dunnett’s test). **B.** After 72 hours of incubation, significant levels of reducing sugars were also detected for the *P. sojae*_247788, 260883, 355355 and 520599 enzymes (Dunnett’s test), indicating that eight out of the eleven paralogs display enzymatic activity towards xyloglucan. *P. sojae*_482953 and *P. sojae*_559651 appeared to show more rapid degradation than the other active paralogs under these conditions. No significant reducing sugars were detected in the vector-only sample (N=3, +/- SD).

*P. sojae*_260883, 355355, 520599, and 247788 enzymes (Dunnett’s test; Figure 2B), indicating that eight out of eleven paralogs display enzymatic activity towards xyloglucan compared to a control yeast carrying an empty vector. However, these data demonstrate that the paralogs have very different rates of catalysis, consistent with a pattern of diversified function. Interestingly, *P. sojae*_482953 and *P. sojae*_559651 (PsXEG1) appeared to show more rapid degradation than the other active paralogs under these conditions (Figure 2A). As both paralogs subtend the ancestral node, at the point when horizontal gene transfer from the fungi to the oomycetes was predicted to occur, they therefore putatively share the conserved function of the ancestral fungal-to-oomycete HGT, i.e. the HGT was likely a *bona fide* xyloglucanase. The shared derived function of this gene family is further suggested by previous studies which report homologous fungal proteins that function as xyloglucanases (55). These results confirm that detectable xyloglucanase function was, however, lost during expansion of this gene family in the branch leading to *P. sojae*_360375 (PsXLP1). Loss of enzymatic function in *P. sojae*_360375 (PsXLP1) has previously been demonstrated to be linked to a deletion event within the peptide-encoding sequence and shown to result in improved host immune-evasion, demonstrating a link between evolution of peptide characteristics and an evolutionary response to host function (43). We also find loss of enzymatic function in the branch leading to *P. sojae*_338064, but note that this paralog expressed in *S. cerevisiae* was subject to manual sequence correction, due to identification of three miscalled introns that were removed from the predicted gene sequence (Supplemental Figure 7). It is therefore possible that this paralog does not accurately reflect the mature transcript expressed by *P. sojae in vivo*, or indeed that this variant is a possible pseudogene. We note that *P. sojae*_520248 displayed enzymatic activity to xyloglucan in a cell culture agar plate-based assay (Figure 1C; Supplemental Figure 5B), but it was not possible to detect function of this paralog from concentrated supernatants, potentially falling below our limit of detection for enzymatic activity.

Both *P. sojae*_482953 and *P. sojae*_559651 (PsXEG1), although representing functional forms that subtend the ancestral HGT phylogenetic node, showed radically different peptide modifications (i.e. in the case of *P. sojae*_559651 (PsXEG1), two separate single amino acid insertions and, in the case of *P. sojae*_482953, a C-terminal extension (Supplemental Figure 1)). These results make it difficult to identify how peptide characteristics relate to xyloglucanase function or the ancestral function of the horizontally-transferred encoded gene, but confirm that this gene family has been subject to additional peptide level variation beyond the deletion identified by Ma et al. (2017) for *P. sojae*_360375 (PsXLP1) (43). To explore this puzzle further, we conducted a series of additional assays to investigate how variation in these peptide characteristics relates to function.

### *P. sojae*_482953 and the orthologs in *Phytophthora cactorum* and *Phytophthora nicotianae* have a highly disordered C-terminal extension which influences catalysis

*P. sojae*_482953 and the orthologs identified from *P. cactorum* and *P. nicotianae* encode disordered C-terminal extensions (Supplemental Figure 1). As such, the protein structures were unable to be accurately modelled using Phyre2 (56) or AlphaFold (57, 58). To investigate the functional significance of the extensions at the C-terminus of *P. sojae*_482953 and orthologs from *P. cactorum* and *P. nicotianae*, truncated versions of the proteins (removing the C-terminal regions) were engineered and expressed in *S. cerevisiae*. Following incubation with 1% (w/v) xyloglucan (pH 7, 30°C) for 6 hours, we find that removal of the extension significantly impairs xyloglucanase function for *P. sojae*_482953 (T-test; p-value = 0.04) and its ortholog in *P. cactorum* (T-test; p-value = 0.0001), but this reduction in enzymatic activity was not found to be significant for the *P. nicotianae* ortholog (Figure 3A).

**Figure 3.**
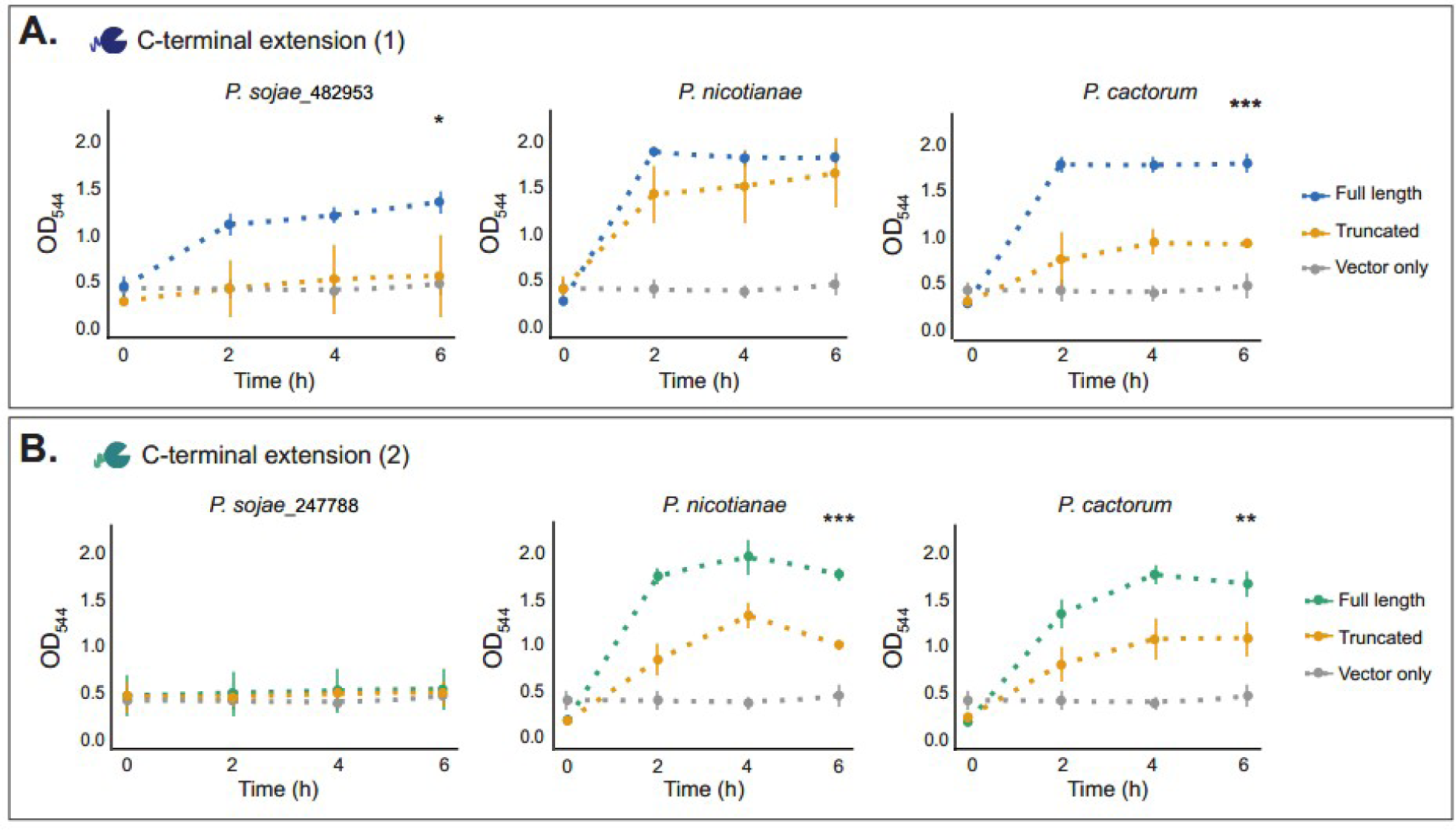
Exploration of function among xyloglucanase paralogs with C-terminal extensions. **A.** *P. sojae*_482953 (full-length) and *P. sojae*_482953 (truncated), and orthologs in *P. cactorum* and *P. nicotianae* secreted into *S. cerevisiae* culture supernatants were incubated with 1% (w/v) xyloglucan at 30°C, pH 7; an increase in absorbance (OD_544_) of DNS reagent added to the samples is suggestive of an increase in the reducing sugars released (i.e. from the breakdown of the substrate). Following incubation with xyloglucan for 6 hours, we find that removal of the C-terminal extension significantly impairs xyloglucanase function for *P. sojae*_482953 (T-test; p-value = 0.04) and its ortholog in *P. cactorum* (T-test; p-value = 0.0001), but this reduction was not found to be significant for the *P. nicotianae* ortholog at any timepoint. No significant reducing sugars were detected in the vector-only sample (N=3, +/- SD). **B.** *P. sojae*_247788 (truncated) and *P. sojae*_247788 (full-length), and orthologs in *P. cactorum* and *P. nicotianae* secreted into *S. cerevisiae* culture supernatants were incubated with 1% (w/v) xyloglucan at 30°C, pH 7; an increase in absorbance (OD_544_) of DNS reagent added to the samples is suggestive of an increase in the reducing sugars released (i.e. from the breakdown of the substrate). *P. sojae*_247788 (full-length) gave weak enzymatic activity towards xyloglucan by this method, but the orthologous proteins of *P. cactorum* and *P. nicotianae* demonstrated significantly higher xyloglucanase function. The truncated orthologs were found to be enzymatically active towards xyloglucan, but with significantly reduced catalysis compared to the full-length proteins after 6 hours incubation with the substrate (T-test; p-value = 0.0002 and 0.01 for *P. nicotianae* and *P. cactorum*, respectively). No reducing sugars were detected in the vector-only sample (N=3, +/- SD).

### *P. sojae*_247788 and the orthologs in *P. cactorum* and *P. nicotianae* also have a highly disordered C-terminal extension

*P. sojae*_247788 is the most closely related GH12 paralog to *P. sojae*_482953, forming a sister relationship with 74% bootstrap support (Figure 1A), and lying in close proximity to this gene on the chromosomal contig (Figure 1E). Interestingly, like *P. sojae*_482953, *P. sojae*_247788 (and orthologs in *P. cactorum* and *P. nicotianae*) also encode a long, disordered C-terminal extension. Similarly, these amino acid extensions were unable to be accurately modelled to a protein structure using Phyre2 (56) or AlphaFold (57, 58). Furthermore, it was not possible to identify any regions of consistently alignable sequence homology between the *P. sojae*_482953 and *P. sojae*_247788 C-terminal tail extensions. Interestingly, we detected low enzymatic activity of *P. sojae*_247788 towards xyloglucan, suggesting reduced function in this branch (Figure 2). Prolonged incubation of *P. sojae*_247788 (full-length and truncated) (72 h), demonstrated no significant differences between these two variant protein forms (Supplemental Figure 6). To further explore this, we repeated our experimental assay for orthologous proteins of *P. sojae*_247788 sampled from *P. cactorum* and *P. nicotianae.* We note that the C-terminal tail extensions were also present in the *P. cactorum* and *P. nicotianae* orthologous proteins, but they showed considerable amino acid alignment variation from *P. sojae*_247788 (Supplemental Figure 1). In contrast to *P. sojae*_247788, the orthologous proteins of *P. cactorum* and *P. nicotianae* demonstrated significantly higher xyloglucanase function (Figure 3B). To investigate if this function was specific to the presence or absence of the C-terminal extensions, truncated versions of the proteins (removing the C-terminal regions) were engineered and expressed in *S. cerevisiae*. The truncated orthologs in *P. nicotianae* and *P. cactorum* were found to be enzymatically-active towards xyloglucan, but with significantly reduced catalysis compared to the full-length proteins after 6 hours incubation with the substrate (T-test; p-value = 0.0002 and 0.01 for *P. nicotianae* and *P. cactorum*, respectively) (Figure 3B). These data suggest that high-level xyloglucanase catalytic function within the 482953 - 247788 clade was, in part, a result of the acquisition of a highly disordered C-terminal extension, but that this high-level function has not been retained specifically in the *P. sojae*_247788 paralog, yet is present in the *P. cactorum* and *P. nicotianae* orthologous proteins.

### The functional *P. sojae*_559651 (PsXEG1) xyloglucanase and its orthologs are predicted to encode an additional substrate-binding site

As discussed above, *P. sojae*_559651 (PsXEG1) and its associated orthologs were shown to have two separate single amino acid insertions (Supplemental Figure 1). Predictions of substrate-binding sites (active site residues) for GH12 paralogs were generated by 3DLigandSite (59). *P. sojae*_559651 (PsXEG1) was the only GH12 paralog in which an additional binding site was predicted; the amino acid residues for this second putative site were Gly55, Ala56, Ala57, Thr58, Val97, Phe205, Val206 (residue position numbers given for the protein sequence in the absence of its N-terminal signal peptide); Thr58 was the common residue predicted in both ligand sites for this paralog (Supplemental Figure 4A). Orthologous proteins in *P. cactorum* and *P. nicotianae* were also predicted to encode this structurally-inferred binding site. Whilst all of the additional binding site residues were found to be conserved amongst the three orthologous protein sequences, for *P. cactorum*, Ala57 was not predicted to form part of its putative binding site. Interestingly, the two putative insertions coding for alanine (Ala, A) and Serine (Ser, S) that are conserved among the three orthologs, were important for defining the additional binding site prediction (Supplemental Figure 1). *In silico* removal of both amino acids from the sequence prior to 3DLigandSite (59) analyses abolished prediction of the second binding site, whilst the ‘primary’ binding site common amongst all GH12 paralogs remained intact. Of all the *P. sojae* GH12 paralog sequences, the two putative indels are present only in *P. sojae*_559651 (PsXEG1) and *P. sojae*_360375 (PsXLP1) (Supplemental Figure 1), however, as the latter paralog is significantly truncated at the C-terminus, a second putative binding site was not predicted for this protein. This paralog was not studied further, as it was investigated previously by Ma et al. (2017), with interesting insights into host-pathogen interactions (43).

To investigate the functional significance of the structurally-inferred binding site, *P. sojae*_559651 (PsXEG1) and its ortholog identified from *P. cactorum* were engineered to delete both the A and S residues. The amended genes were heterologously expressed in *S. cerevisiae*, but we found no difference in xyloglucanase activity as monitored by the release of reducing sugars over 6 hours incubation (pH 7, 30°C) (Supplemental Figure 4B). Taken together, the data suggest an as yet unknown role, if any, for the structurally-inferred additional binding site in *P. sojae*_559651 (PsXEG1).

### *P. sojae*_482953, 260883 and 520599 trigger rapid generation of reactive oxygen species (ROS) in *Nicotiana benthamiana*

The generation of Reactive Oxygen Species (ROS) in plants is a marker for stress and early immune responses (60). To investigate how plant immunity is modulated in response to each of the xyloglucanase variants, the eleven paralogs (and a vector-only negative control) were separately infiltrated into *N. bethamiana* leaves by agroinfiltration. Leaf discs were collected at 48 h post-infiltration and incubated in water overnight. The generation of ROS was then measured from 0 to 60 minutes. ROS accumulation was detected within minutes of adding ROS assay reagent for leave discs expressing *P. sojae*_482953, 260883 and 520599 paralogs, and remained high for the entire measurement period (Figure 4A). Total photon counts at 60 minutes suggests that the ROS response to *P. sojae*_482953 was significantly higher than that of *P. sojae*_260883 and 520599 (Figure 4B). *P. sojae*_482953 represents one of the most active enzymes in this gene family under the conditions we tested (Figure 2A); to further probe if the ROS response triggered by this variant protein could be attributed to the presence of the C-terminal extension (which we demonstrated increases its xyloglucanase activity (Figure 3A)), the *P. sojae*_482953 C-terminal tail only was infiltrated into *N. benthamiana* leaves. Interestingly, we find that the C-terminal extension alone triggers ROS generation, but significantly reduced compared to the full-length protein, and not statistically significant from the vector-only control (Supplementary Figure 2).

**Figure 4.**
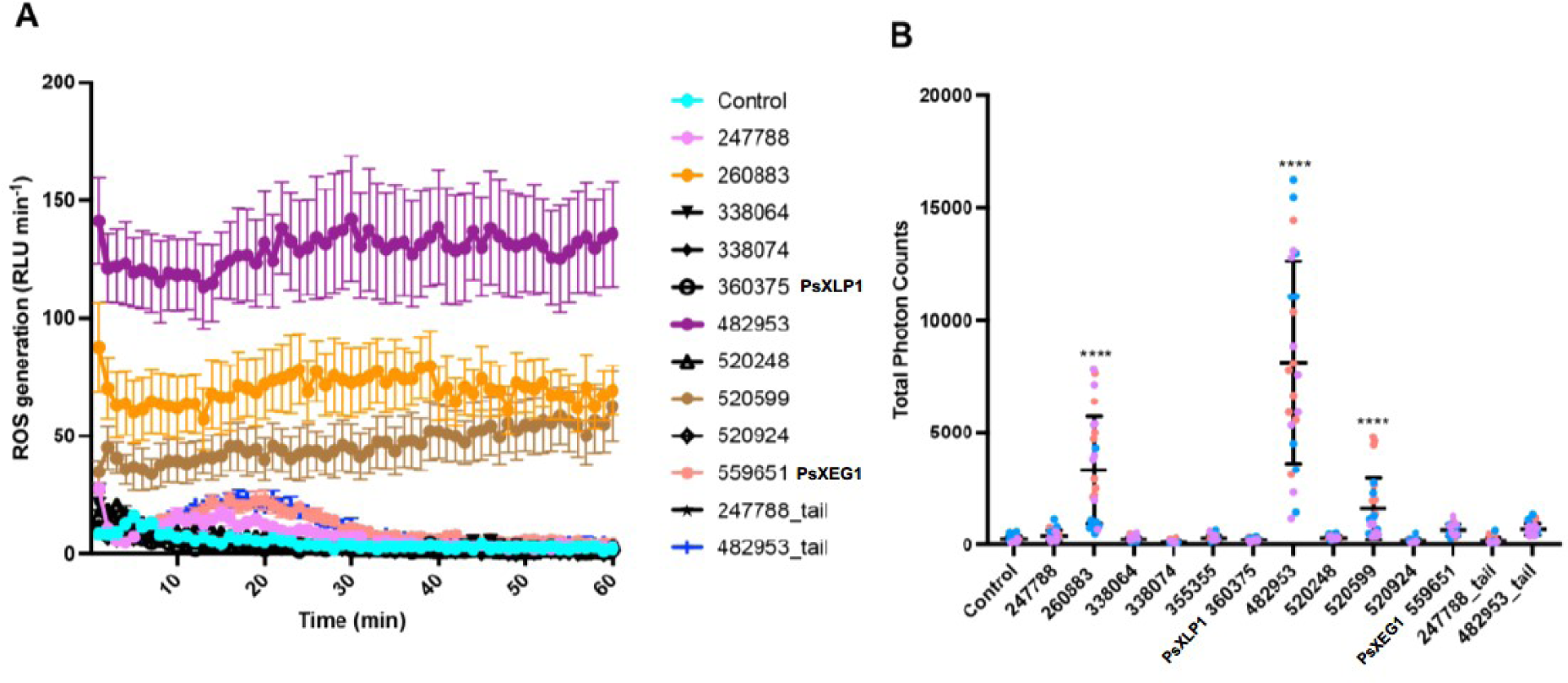
*P. sojae* xyloglucanase paralogs induction of ROS responses in a model plant system. **A.** *P. sojae* xyloglucanase paralogs were codon-optimized for expression in plants and infiltrated into the leaves of *N. benthamiana*. The dynamics of ROS production in *N. benthamiana* plants was measured from 0-60 minutes. ROS production corresponds to the mean of 8 samples (+/- SE). **B.** Total ROS production over 60 minutes. Horizontal bars represent the mean and SD. Asterisks showing p-value < 0.0001 using T-test. The experiments were performed 3 times (orange: replicate 1, blue: replicate 2, purple: replicate 3).

Intriguingly, ROS accumulation was also significantly enhanced by *P. sojae*_260883 and *P. sojae*_520599 proteins – these paralogs showed significantly reduced xyloglucanase activity compared with *P. sojae*_482953 (Figure 2). Furthermore, gene expression of *P. sojae*_520599 is undetectable in all life stages, whilst *P. sojae*_260883 shows limited expression during plant infection in comparison with *P. sojae*_482953 (Figure 1B).

Agroinfiltration of *P. sojae*_247788 and 559651 (PsXEG1) also triggered ROS generation in *N. benthamiana*, but significantly reduced compared to *P. sojae*_482953, 260883 and 520599 proteins, and total photon counts at 60 minutes were not statistically significant from the vector-only control (Supplementary Figure 2). Interestingly, ROS accumulation in response to these xyloglucanase variants peaked at around 20 minutes post-infiltration and declined by 60 minutes of observation (Figure 4A). This pattern was not observed in any of the variants that triggered the highest ROS accumulation (*P. sojae*_482953, 260883 and 520599), suggesting that these paralogs have diverse roles in triggering plant immunity functions. We did not find evidence of ROS accumulation as a result of infiltration with *P. sojae*_338064, 338074, 355355, 360375 (PsXLP1), 520248, 520924, and the *P. sojae*_247788 C-terminal extension only (Supplementary Figure 2).

Bacterial-associated flg22 and fungal-associated chitin represent well-known pathogen-associated molecular patterns (PAMPs), both triggering ROS generation *in planta* (61–63). To explore whether the xyloglucanase variants could enhance or inhibit flg22- and chitin-triggered ROS generation, we infiltrated *P. sojae* proteins into *N. benthamiana* leaves and applied flg22 or chitin treatment. Interestingly, we find that *P. sojae*_247788 and 260883 can significantly suppress flg22-triggered ROS generation in *N. benthamiana* (Supplemental Figure 3A). However, no significant effects of the xyloglucanase variants on chitin-triggered ROS generation were observed (Supplemental Figure 3B).

### *P. sojae* GH12 paralogs cleave the same glycosidic linkages of the xyloglucan backbone, but show differences in their oligosaccharide products

Xyloglucan oligosaccharides released by *P. sojae*, *P. nicotianae*, and *P. cactorum* xyloglucanases (or a vector-only control) upon digestion of 1% (w/v) xyloglucan (72 h, pH 7, 30°C), were analysed by mass spectrometry (MALDI) (analysis carried out by Wyatt Analytical Ltd). Four peaks of interest in the mass spectra were observed across the samples containing functional enzyme variants; ions with *m/z* of ∼1085, 1247, 1409, and 1571 – putatively corresponding to the xyloglucan oligosaccharides XXXG, XXLG (or XLXG), XLLG, and XLFG, respectively (24). These species were not observed in the vector-only controls – consistent with the observed xyloglucanase activity of the functional enzyme variants (Figure 2). The relative intensities of species identified at m/z ∼1085, 1247, 1409, and 1571 were compared by calculating the ratios between the areas under the peaks.

Figure 5 provides strong evidence that the composition of oligosaccharides varies significantly between active *P. sojae* xyloglucanase paralogs (Dirichlet likelihood ratio test (randomization approach); p-value < 0.001). Goodness-of-fit testing indicates that the Dirichlet distribution provides an adequate model for these data (p-value = 0.48). We suggest that putative differences in enzymatic output across the wider GH12 family could act to increase the efficiency of substrate breakdown during *P. sojae* growth and infection (Figure 5). We see virtually undetectable levels of XLFG for *P. sojae*_482953, *P. sojae*_559651 (PsXEG1), and *P. sojae*_338074, representing the two most active xyloglucanase variants under the conditions we tested (*P. sojae*_482953 and *P. sojae*_559651 (PsXEG1)), with *P. sojae*_338074 also showing significant enzymatic activity at 6 h (Figure 2A)) - suggesting that the individual enzymatic activities of these variants do not especially act to generate this particular oligosaccharide product. Intriguingly, *P. sojae*_482953 is the variant we demonstrate has a C-terminal extension that increases activity (Figure 3A), and this protein triggers the highest generation of ROS in *N. benthamiana* (Figure 4). We also observed an increased output of XLFG for *P. sojae*_355355, and this paralog is expressed during all three *P. sojae* lifecycle stages (it is notably amongst the most highly expressed of the GH12 paralogs in the cyst and infection stages; Figure 1B). Our results also demonstrate increased output of XXXG for *P. sojae*_260883, and this paralog is among the most highly expressed of the GH12 paralogs in the cyst stage (Figure 1B), as well as triggering significant ROS accumulation in *N. benthamiana* (Figure 4). Differences in enzymatic output between the enzyme variants across different stages of the *P. sojae* lifecycle could not only provide more effective breakdown of xyloglucan, but could also act to diversify the types of xyloglucan oligosaccharides that are released and therefore perceived as DAMPs by the plant host. Further work is necessary to probe the effects, if any, of the role of individual xyloglucan oligosaccharides in plant immune responses.

**Figure 5.**
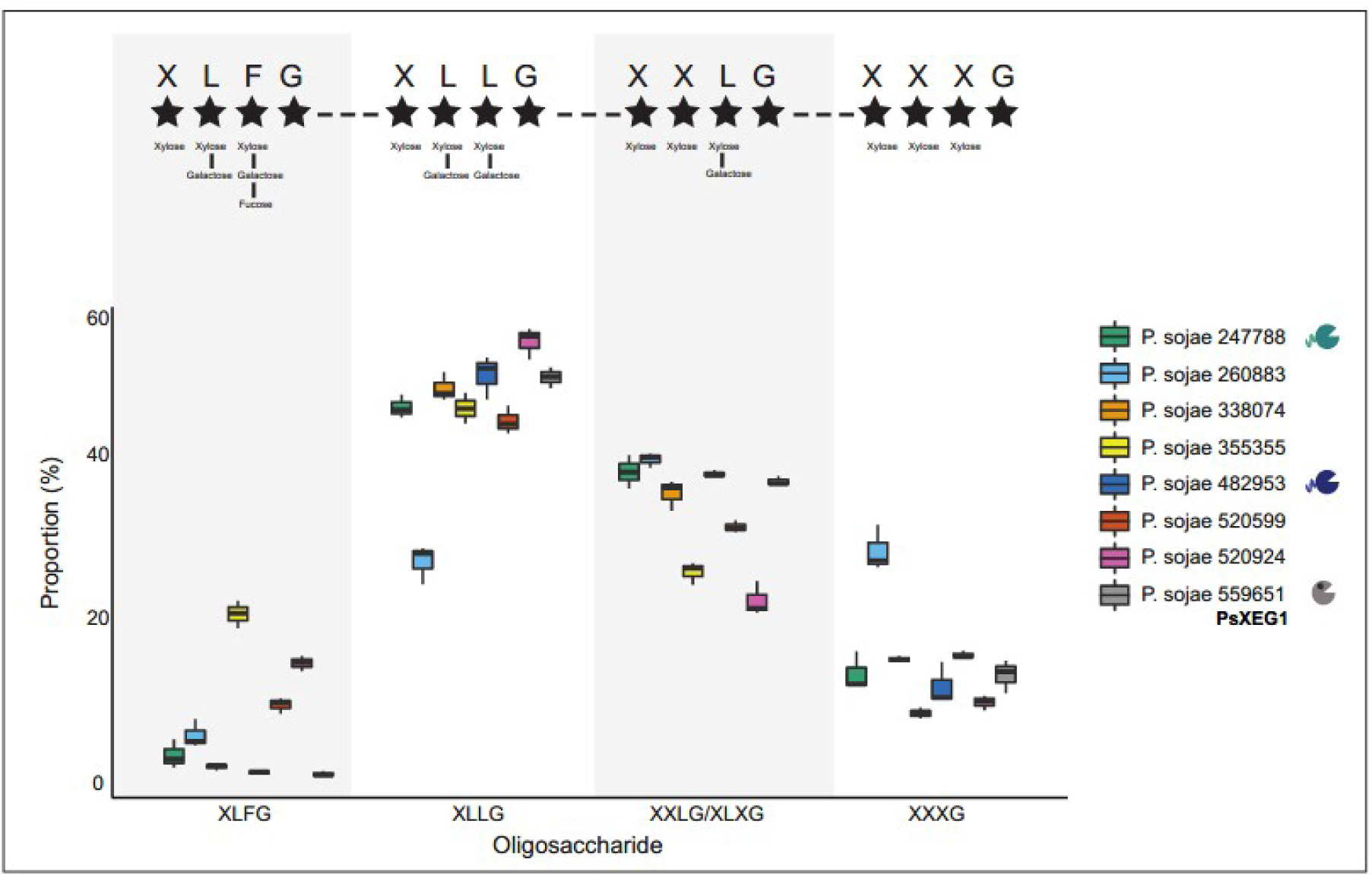
*P. sojae* xyloglucanase paralogs produce variant oligosaccharide break-down products. *P. sojae* xyloglucanase paralogs secreted into *S. cerevisiae* culture supernatants were incubated with 1% (w/v) xyloglucan at 30°C, pH 7, for 72 h. MALDI-MS spectra confirmed the release of xyloglucan oligosaccharides by eight of the eleven variants (corresponding to the functional paralogs identified in Figure 2). Four peaks of interest were observed; ions with *m/z* of ∼1085, 1247, 1409 and 1571 – putatively corresponding to the oligosaccharides XXXG, XXLG (or XLXG), XLLG, and XLFG respectively (24). The top panel indicates the side chain residues present in each of the oligosaccharide variant types, as described by Fry et al., 1993 (24). The relative intensity of species identified at these peaks were compared by calculating the ratios between the areas under the peaks, to probe putative differences in preferential binding of the xyloglucan backbone. We find strong evidence that the composition of oligosaccharides varies significantly between the *P. sojae* xyloglucanase paralogs (Dirichlet likelihood ratio test (randomization approach); p-value < 0.001). For example, we see almost absence of the XLFG product for three enzymes. Goodness-of-fit testing indicates that the Dirichlet distribution provides an adequate model for these data (p-value = 0.48).

We also tested whether the composition of oligosaccharides varied between the full-length and truncated xyloglucanase variants for each ortholog. For C-terminal extension 1 (*P. sojae*_482953; Figure 6A), the analysis indicated a significant difference in the composition of oligosaccharides produced by the full-length and truncated *P. nicotianae* variants (Dirichlet likelihood ratio test (randomization approach); p-value < 0.001), but no significant difference between the *P. sojae* (p-value = 0.098) or *P. cactorum* (p-value = 0.104) variants. For C-terminal extension 2 (*P. sojae*_247788; Figure 6B), the analysis indicated a significant difference in the composition of oligosaccharides produced by full-length and truncated *P. nicotianae* variants (Dirichlet likelihood ratio test (randomization approach); p-value <0.001), but no significant difference between the *P. cactorum* variants (p-value = 0.392) variants. We find no evidence of significantly altered oligosaccharide profiles when comparing *P. sojae*_559651 (PsXEG1) with and without A and S residues (Supplemental Figure 4C).

**Figure 6.**
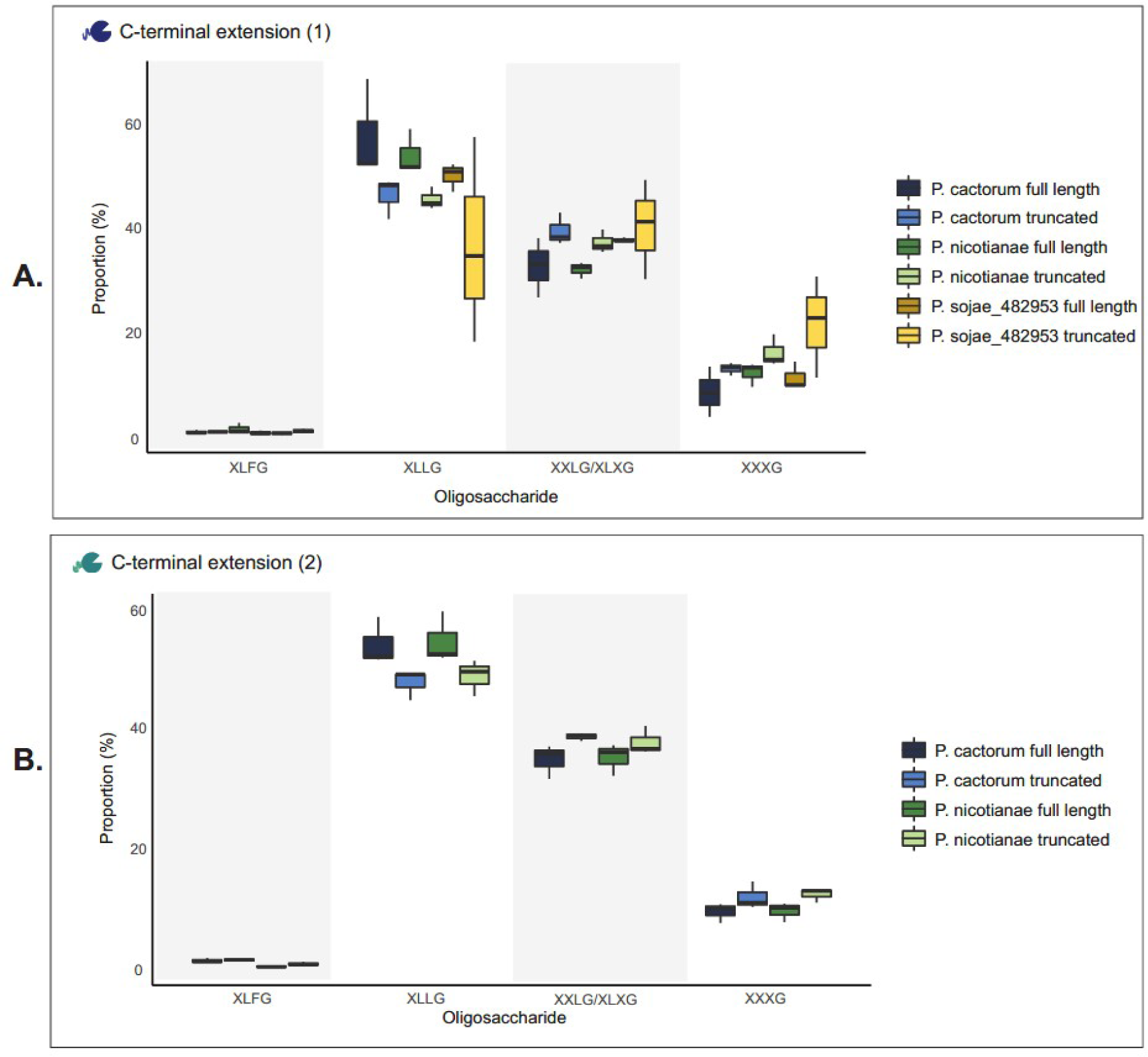
*P. sojae* xyloglucanase paralogs with C-terminal extensions do not produce variant oligosaccharide break-down products dependent on C-terminal tail extension. We tested whether the composition of oligosaccharides varied between the full-length and truncated xyloglucanase variants for each ortholog. Xyloglucanase variants secreted into *S. cerevisiae* culture supernatants were incubated with 1% (w/v) xyloglucan at 30°C, pH 7, for 72 h. MALDI-MS spectra confirmed the release of xyloglucan oligosaccharides. Four peaks of interest were observed; ions with *m/z* of ∼1085, 1247, 1409 and 1571 – putatively corresponding to the oligosaccharides XXXG, XXLG (or XLXG), XLLG, and XLFG respectively (24). The relative intensity of species identified at these peaks were compared by calculating the ratios between the areas under the peaks, to probe putative differences in preferential binding of the xyloglucan backbone. **A.** For C-terminal extension 1 (*P. sojae*_482953), we saw a significant difference in the composition of oligosaccharides produced by the full-length and truncated *P. nicotianae* variants (Dirichlet likelihood ratio test (randomization approach); p-value < 0.001), but there was no significant difference between each of the *P. sojae* (p-value = 0.098) or *P. cactorum* (p-value = 0.104) variants. **B.** For C-terminal extension 2 (*P. sojae*_247788), we saw a significant difference in the composition of oligosaccharides produced by the full-length and truncated *P. nicotianae* variants (Dirichlet likelihood ratio test (randomization approach); p-value <0.001), but there was no significant difference between the *P. cactorum* variants (p-value = 0.392). The MS spectra for *P. sojae_247788* (truncated) were noisy, so this paralog was omitted from (B) and our analysis.

### Targeted deletion of the *P. sojae*_482953 gene does not affect xyloglucan utilisation *in vivo*

To investigate the impact of single HGT paralogs on *P. sojae* xyloglucan utilization *in vivo*, the gene encoding *P. sojae*_482953 was selected as a candidate for deletion, using CRISPR/Cas9 methods developed previously (64). *P. sojae*_482953 showed rapid degradation of xyloglucan compared to most of the other enzyme variants (Figure 2A), as well as high gene expression during the *P. sojae* lifecycle (Figure 1B). Therefore, we hypothesised that this paralog could play a significant role in xyloglucan digestion *in vivo*. Employing the CRISPR/Cas9 system, we mutated *P. sojae*_482953 in the reference strain P6497 (Supplemental Figure 8); however, we found no significant effect to *P. sojae* growth on agar plates supplemented with 1% (w/v) xyloglucan as the sole carbon source, between the wild-type strain and the homokaryotic mutant (Supplemental Figure 9A). This suggests that the remaining xyloglucanase paralogs (or other encoded glycoside hydrolases in *P. sojae*) are able to compensate for deletion of *P. sojae*_482953 and conceal the loss of its function – indicating that gene duplication events leading to functional redundancy are important for maintaining stable function. This experiment suggests, in this one case, that enzymatic function is not a driving force for maintaining a diverse suite of xyloglucanase paralogs, but rather that different selective forces are driving the maintenance of numerous xyloglucanase variants possibly, for example, to modify the host immune response.

We also tested whether the composition of oligosaccharides released during xyloglucan digestion varied between the wild-type strain and mutant grown in liquid culture, but we found no evidence of altered enzymatic products (Supplemental Figure 9B). Further work would be useful to probe the impact of HGT paralog genes on *P. sojae* xyloglucan utilization and phytopathogenicity.

## Discussion

The genomes of necrotrophic and hemibiotrophic plant pathogens are replete in genes predicted to encode secreted degradative enzymes (26, 27). Many of these enzymes are thought to function in the breakdown of the plant cell wall and additional plant tissues, which are both defensive structures and rich in valuable polysaccharides (17, 18). Horizontal gene transfer has been shown to be a key factor in the evolution of gene families predicted to function in breakdown of plant tissues (28–33, 65). Many of these gene families demonstrate evidence of multiple rounds of gene duplication post-transfer, pointing towards a process of repeat selection for retention of duplicated genes, which may subsequently drive transcriptional amplification (gene dosage); neofunctionalization of paralogs; and/or co-option of paralogs for transcriptional segregation associated with specific steps in development or host interaction. To explore this phenomenon further we investigated the functional evolution of a HGT gene family (GH12) in *P. sojae.* We were able to demonstrate that this gene family encodes eight paralogs with detectable xyloglucanase function, each with variant transcriptional profiles. We show that some functional paralogs have diversified additional structural features important for xyloglucanase function, as well as unique ROS accumulation profiles in *N. benthamiana*. We also show that xyloglucanase function amongst the wider *P. sojae* GH12 family results in diversified enzyme digestion products. Taken together, these results suggest that gene duplication and paralog evolution has played an important role not only in the modulation of xyloglucanase function, but in the alteration of host perception during pathogen attack.

*P. sojae*_482953 and *P. sojae*_247788 encode significantly disordered C-terminal extensions; truncation of *P. sojae*_482953, and orthologous proteins in *P. nicotianae* and *P. cactorum* reduces enzymatic activity upon xyloglucan– consistent with a hypothesis that the C-terminal extensions play an important role in maintaining interactions with the enzyme substrate (therefore optimizing the encoded enzyme activity at the conserved catalytic sites). Interestingly, Li et al. (2014) describe a C-terminal proline-rich sequence of xylanase XynA, and its removal also negatively affects the protein function (66). In contrast, Wen et al. (2005) describe a truncated glucanase that displays improved enzymatic activity compared to its full-length counterpart (67), suggesting that evolution of a C-terminal extension can both positively and negatively affect enzymatic function, depending on their interactions with putative substrates. We find that *P. sojae*_247788 (full-length) has minimal xyloglucanase function, although interestingly, this paralog has significantly reduced expression at any of the life-cycle stages tested previously (FungiDB; (51, 52)) (Figure 1B). Conversely, full-length orthologs in *P. cactorum* and *P. nicotianae* display increased xyloglucanase activity, and truncation of the C-terminal extensions reduces the rate at which both proteins can digest xyloglucan. These data suggest that addition of these C-terminal extensions has further amended the function of the HGT-derived xyloglucanase post-gene duplication events.

*P. sojae*_559651 (PsXEG1) and orthologs in *P. cactorum* and *P. nicotianae* were predicted to encode an additional substrate binding site, with two indels coding for alanine and serine found to be important for the bioinformatic prediction of this second binding site. We hypothesised that an additional binding pocket could allow the GH12 paralog to interact with xyloglucan by a novel mechanism– potentially able to bind more of the carbohydrate backbone or maintain closer interactions with the substrate, enabling more efficient digestion. Heterologous expression of *P. sojae*_559651 (PsXEG1) in *S. cerevisiae* without the indels does not support this hypothesis; instead, we find there is no difference in catalytic activity following removal of the indels that give rise to the structurally-inferred binding site. Further work is therefore necessary to elucidate the function, if any, of this additional putative binding site. The two indels are also conserved in *P. sojae*_360375 (PsXLP1), however, as this paralog is significantly truncated at the C-terminus (missing a glutamic acid residue required for enzymatic activity (see Supplemental Figure 1)), a second putative binding site was not originally predicted. Previously, *in vivo* work by Ma et al. (2017) demonstrated that this *P. sojae* protein transiently expressed in *N. bethamiana* leaves does not result in hydrolytic activity– consistent with both glutamic acid residues being required for catalytic activity. Interestingly, despite loss of enzymatic activity, the researchers demonstrated that this protein has a strong binding affinity to a host immune protein, GmGIP1 (43) – demonstrating a role for non-active isozymes in *P. sojae* virulence as immune system ‘decoys’. Therefore, strong selection pressures resulting from intimate pathogen-plant interactions suggest that evolution of multiple enzyme paralogs is also important for subverting or exploiting host defenses. Additionally, Xia et al. (2020) show that N-glycosylation of *P. sojae*_559651 (PsXEG1) protects it against degradation by the host protease GmAP5, and also attenuates binding by GmGIP1 (68). Recently, Sun et al. (2022) solved the crystal structure of the same enzyme variant bound to a leucine-rich repeat (LRR) receptor-like protein (RLP) from *N. benthamiana*, and demonstrated key residues involved in inactivation of the enzyme upon binding (pdb: 7DRB) (69). It is therefore possible that the function of the structurally-inferred binding site in *P. sojae*_559651 (PsXEG1) could be related to interactions with the plant immune response system.

To probe putative differences in host perception of the eleven xyloglucanase variants, each one was infiltrated into the leaves of *N. benthamiana*, and ROS accumulation monitored from 0 to 60 minutes. *P. sojae*_482953, 260883 and 520599 resulted in significant accumulation of ROS, and ROS generation in response to *P. sojae*_482953 is largely independent of the proteins C-terminal extension. Negligible accumulation of ROS was observed in response to *P. sojae*_247788 and *P. sojae*_559651 (PsXEG1), and interestingly, both xyloglucanase paralogs triggered a peak ROS response around 20 minutes post-infiltration before declining– a pattern we found unique to both of these variants. In previous work by Ma et al. (2015), the authors demonstrate that *P. sojae*_559651 (PsXEG1), 482953, 260883 and 247788 induce cell death in *N. benthamiana* (44) – consistently, we detected ROS accumulation as a result of infiltration of these proteins, albeit at remarkedly different levels between paralogs. Interestingly, *P. sojae*_520599 was not demonstrated to induce cell death in previous work (44), but our results suggest infiltration of this protein into *N. benthamiana* leaves results in the third highest accumulation of ROS amongst the xyloglucanase variants in this gene family. According to publically-available RNAseq data for *P. sojae*, there is no evidence that *P. sojae*_520599 is transcriptionally-active during plant infection – so it is interesting to consider whether the reduced expression of this gene is a response to the host-pathogen arms race to limit host immune perception. Further focused work on this xyloglucanase paralog would be useful to uncover how the evolution of this protein has been shaped by host immune responses over evolutionary time.

Investigation of xyloglucan oligosaccharides released following xyloglucan digestion suggests that enzymatic activity of *P. sojae* GH12 paralogs leads to diversified enzymatic output. We hypothesize that this example of neofunctionalization of paralogs can offer a greater efficacy of *in vivo* xyloglucan digestion, achieved through the secretion of multiple catalytically-active GH12 paralogs with differences in binding of the xyloglucan chain. We also suggest that the variation in oligosaccharides released as a product of the function of active GH12 paralogs may contribute to the diversification of the DAMPs released during pathogen invasion. Previous work by Claverie et al. (2018) confirms that xyloglucan oligosaccharides are recognized as DAMPs and trigger a broad range of immune responses (25) (although we do not know if different xyloglucan oligosaccharides promote the same response). It is therefore possible that duplication and neofunctionalization of xyloglucanase output would require the host immune system to maintain a high sensitivity of detection for one or more specific DAMP signatures among a diversity of alternative carbohydrate breakdown products present in dynamic and fluctuating relative concentrations. Further work to disentangle host responses to individual xyloglucan oligosaccharides would help to fully understand the relevance, if any, of the diversified enzymatic output we observed in our study.

The Red Queen hypothesis (inspired by Lewis Carroll’s 1871 novel, *Through the Looking-Glass*) is useful for understanding host-pathogen co-evolutionary dynamics (45, 46), and predicts that evolution will act to generate patterns of diversification on any molecular signature that activates a host immune response (e.g. (47)). Consequently, the pattern of gene duplication, loss (Figure 1A), deactivation and neofunctionalization seen in the *P. sojae* GH12 xyloglucanase paralog family could be a factor in the host-pathogen arms race. Specifically, we present a variant paralog with a C-terminal extension that increases enzymatic activity, as well as demonstrate diverse ROS accumulation patterns *in planta*, and diverse profiles of enzymatic breakdown products amongst the members of this HGT xyloglucanase family. We therefore suggest that the host immune system must perceive and respond to fluctuations in multiple protein signatures, enzymatic catalysis rates and, potentially, multiple DAMP signatures. If the host fails to recognize the function or effects of one paralog, a selective sweep would quickly favour a variant strain, utilizing a paralog that produced a profile that was not detected, allowing immune evasion and therefore selection for that variant. Consistent with this hypothesis, this gene family shows multiple paralogs that have reduced detectable xyloglucanase function, and paralog forms which do not trigger ROS accumulation, and others which show near-absence of some classes of oligosaccharide output – a possible product of pathogen protein evolution with the selected outcome of escaping host immune detection. If, and when, the plant immune system catches up with a pathogens amended protein signature, a variant set of paralogs with an altered signature could then be selected for prominence, allowing improved evasion of the plant immune system and resulting in fixation of additional gene duplication forms with neofunctionalized functions. Collectively, we argue the data presented here and in Ma et al. (2015 and 2017) demonstrates that the *P. sojae* GH12 xyloglucanase paralog families show such an evolutionary dynamic. Taken together, these results contribute to a better understanding of the functional differences between gene duplications post-HGT, giving us improved insight into the dynamics of phytopathogenicity, as well as a better understanding of the consequences of gene flow in phytopathogenic oomycetes.

## Materials and methods

### Phylogenetic analyses

Candidate homologs (paralogs) of *P. sojae* GH12 (Pfam: PF01670, InterPro: IPR002594) were collated by performing Hidden Markov Model (HMM) searches against Ensembl genomes (restricted by taxonomy to *P. sojae* NCBI taxids 67593 and 1094619) with default parameters (48), using the raw profile HMM training set for the putative protein family (Pfam: (49)). A phylogenetic tree was constructed to investigate the evolutionary relationships between putative *P. sojae* GH12 paralogs and other oomycetes. GH12 protein homologs were identified from a selection of eukaryotic (with sampling subsequently focusing on oomycetes and fungi) and prokaryotic genomes using BLASTp (70); from these hits a multiple sequence protein alignment was constructed and aligned using automated methods in Seaview (71) using MUSCLE (72), which was then edited and masked manually. GenBank was checked for additional gene sampling of oomycetes not included in the genome-by-genome survey, and additional prokaryote and eukaryote - including fungal – genes. A preliminary phylogenetic tree was calculated using PhyML (73) to confirm all *P. sojae* GH12 paralogs sampled were part of the same oomycete monophyletic clade and predicted to be a HGT from fungi (31, 33); this was then checked manually (enabling the removal of distantly-related fungal and prokaryotic outgroups and partial in group sequences). The alignment was subject to an additional round of manual editing and masking, resulting in an alignment with 161 sequences and 211 amino acid sites. A second PhyML tree was calculated and checked, and the final ML tree was constructed with IQ-Tree v2.0.3 with ModelFinder (74) identifying WAG+R5 model. The tree was subject to non-parametric bootstrap with 200 pseudoreplicates (75), and rooted with a fungal outgroup (31, 33). The final tree was visualized with iTOL (76).

### Comparisons of putative oomycete GH12 protein structures

*P. sojae* GH12 protein sequences were aligned using Clustal Omega (77, 78), and visualized in Jalview2 (79). Three-dimensional structures were obtained with Phyre2 (Protein Homology/analogY Recognition Engine v2.0 (56) or AlphaFold (57, 58), using protein sequences without their predicted N-terminal signal peptide (determined using SignalP-6.0 (80)). Predictions of carbohydrate-binding sites for putative paralogous proteins were generated by 3DLigandSite (date searched: June/2021); the server was used to identify high-scoring homologous protein structures with ligands bound (by comparative MAMMOTH score) (59).

### Identification of transcription responses of the GH12 gene family in *P. sojae*

FungiDB contains comparative RNAseq data for *P. sojae* ensemble transcriptome samples from lifecycle stages: mycelial, cyst and 3 days post-infection (soybean hypocotyls infected with *P. sojae* strain P6497) (51, 52). Relative transcription of the GH12 paralogs were identified as transcript levels of fragments per kilobase of exon model per million mapped reads (FPKM - sequencing depth and gene depth normalised from paired-end RNA-seq data; FungiDB; (51, 52)). RNAseq data for *P. sojae* was additionally used to manually check GH12 gene models prior to the synthesis of genes for experimental study.

### Gene synthesis of GH12 genes from *P. sojae* and relatives

We identified 11 *P. sojae*-specific GH12 paralogs (see Figure 1). All 11 *P. sojae* paralogs were codon optimized for translation in *S. cerevisiae* (http://www.genscript.com/tools/rare-codon-analysis). The gene sequences were synthesized by Synbio Technologies into the vector p426-GPD (ATCC 87361) between *BamHI* (GGATCC) and *HindIII* (AAGCTT) restriction sites. *P. sojae* N-terminal signal peptide sequences were replaced with *S. cerevisiae* mating factor α (MFA) (53), in order to maximize heterologous protein secretion in yeast. MFA sequence: ATGAGATTTCCTTCAATTTTTACTGCTGTTTTATTCGCAGCATCCTCCGCATTAGCTGCTCCAGTCAACACTACAACAGAAGATGAAACGGCACAAATTCCGGCTGAAGCTGTCATCGGTTACTCAGATTTAGAAGGGGATTTCGATGTTGCTGTTTTGCCATTTTCCAACAGCACAAATAACGGGTTATTGTTTATAAATACTACTATTGCCAGCATTGCTGCTAAAGAAGAAGGGGTATCTCTCGAGAAAAGAGAGGCTGAAGCT.

### Assays for GH12 xyloglucanase function

To compare xyloglucanase function we conducted activity assays against xyloglucan (Tamarind; backbone of β-1,4-glucan; most substituted (Megazyme)). Yeast transformants were screened for released reducing sugars using 3,5-Dinitrosalicylic acid (DNS) reagent as follows: recombinant *S. cerevisiae* strains (in biological triplicate and including a p426-GPD vector-only control strain) were cultured in 20 mL synthetic complete medium (1% glucose) lacking uracil (SCM-URA) for 7 days at 30°C with 180 rpm shaking. Culture supernatants were removed by centrifugation at 4°C, concentrated 10x (Corning Spin-X UF concentrator; 10,000 kDa MWCO) and kept on ice. Total protein concentrations were measured using a Qubit Fluorometer and Thermo Scientific reagents. Concentrated supernatants were matched to 75 µg/mL, and incubated with 1% (w/v) xyloglucan in 50 mM citrate buffer (pH 7) at 30°C with 180 rpm shaking. At each time point sampled, released reducing sugars were measured by removing 60 µL of each sample into a sterile 1.5 mL tube, and adding 60 µL of DNS reagent (1% (w/v) DNS, 1M potassium sodium tartrate, 400 mM sodium hydroxide), followed by incubation at 95°C for 5 minutes to allow colour development. Samples were transferred to a sterile 96-well plate, and the absorbance was measured at 544 nm using an absorbance microplate reader (CLARIOstar; BMG LABTECH).

For the Congo red assays (Supplemental Figure 5A), 10 μL of the concentrated supernatants at 75 μg/mL total protein, were spotted onto SCM-URA agar containing 0.2% (w/v) xyloglucan (with 2% (w/v) glucose as an additional carbon source). Agar plates were incubated at 30°C for 24 hours (static), and the remaining intact polysaccharide on the plates was stained with 0.2% (w/v) Congo red (Sigma Aldrich) for 30 minutes at room temperature, and de-stained with 1 M sodium chloride for 30 minutes (with shaking) at room temperature (81). Extracellular enzyme activity was indicated by a clearing or ‘halo’ around the spots containing secreted functional enzyme.

For the cell culture-based Congo red assays (Supplemental Figure 5B), we followed an identical protocol as above, except 10 μL of *S. cerevisiae* cell culture (i.e. prior to centrifugation and collecting the supernatant) were spotted onto the SCM-URA (0.2% (w/v) xyloglucan) agar plates.

All data analysis was performed using R version 4.0.3. DNS assay data (Figure 2) were corrected using the initial (0 h) OD reading for each sample, to account for residual sugars present in concentrated supernatant samples; these data were then analysed using a Dunnett Test against the vector-only control, using package DescTools (82). DNS assay data (Figure 3, Supplemental Figure 4B) are displayed as OD readings at each timepoint; 6 h data were then analysed using a Student’s t-test against the vector-only control (two-tailed, two sample equal variance) (83).

### Detection of reactive oxygen species (ROS)

GH12 paralogs were codon-optimised for translation in *N. benthamiana* plants, and the gene sequences were synthesised by Genewiz and Twist Bioscience. Synthesised DNA fragments were cloned into vector PGWB514 (Addgene 74856) (84) by In-Fusion cloning (Takara Bio). Verified plasmids were then introduced into *Agrobacterium* strain AGL1 by a freeze-thaw method (85). Four-week-old *N. benthamiana* leaves were infiltrated with *Agrobacterium* strain AGL1 (OD_600_ = 0.4) containing the corresponding GH12 genes. Eight leaf discs from each infiltrated vector were collected at 48 h post infiltration. Leaf discs were incubated in water in white 96-well plates (Greiner Bio-One 655075) overnight. Water was then replaced with ROS assay reagent containing 100 µM Luminol (Merck 123072) and 20 µg/mL horseradish peroxidase (Merck P6782). ROS generation was measured at 0-60 minutes using a High-Resolution Photon Counting System (Photek HRPCS218). For the PAMP treatment assay, 1 μM flg22 or 1 mg/mL chitin was added to the ROS assay reagent.

### Mass Spectrometry analysis of xyloglucan digestion products

Mass Spectrometry was utilised to investigate the differences in degradation products released from xyloglucan breakdown by the xyloglucanase variants. Yeast transformants were prepared for MS analysis as follows: recombinant *S. cerevisiae* strains (in biological triplicate and including a p426-GPD vector-only control strain) were cultured in 20 mL SCM-URA (1% glucose) for 7 days at 30°C with 180 rpm shaking; the supernatants were removed by centrifugation and concentrated to 10x, as described above. Concentrated supernatants (matched to 75 µg/mL) were incubated with 1% (w/v) xyloglucan in 50 mM citrate buffer, pH 7 for 72 h at 30°C with 180 rpm shaking. At 0 and 72 h, 60 µL of each sample was removed into a sterile 1.5 mL tube, and the reactions terminated by heating at 95°C for 5 minutes. The samples were air-dried using a centrifugal vacuum concentrator. MALDI data was acquired in positive-reflectron mode (Bruker ultrafleXtreme spectrometer); all sample preparation, optimisation, and sample analysis were carried out by Wyatt Analytical Ltd.

Mass spectrometry data for paralog oligosaccharide diversity were analysed using a Dirichlet likelihood ratio test using R code published previously (86). A randomisation procedure was used to determine p-values (5,000 random samples), and goodness-of-fit testing was used to test whether the Dirichlet distribution provided an adequate model for this dataset, using 10,000 simulations (86).

### Deletion of the gene encoding *P. sojae*_482953 by CRISPR/Cas9 methods

*P. sojae* P6497 was used in this study and was routinely maintained on V8 agar plates at 25°C in the dark. CRISPR/Cas9 methods were used to disrupt the gene encoding *P. sojae*_482953 and replace it with the coding sequence of enhanced Green Fluorescent Protein (eGFP), as per Fang et al. (2017) (64). sgRNAs were designed and cloned into the backbone plasmid pYF515 by an annealing method according to Fang et al. (2017) (64). The Homology-Directed Repair (HDR) template used for gene replacement was assembled into plasmid pBluscript KS-using HiFi assembly (NEB, E2621S). Two independent mutants were obtained based on two different protoplast pools and different sgRNAs (Supplemental Table 1). Diagnostic PCR was performed to screen mutants based on the primers listed in Supplemental Table 1. Gene deletion mutants were also maintained on V8 agar plates at 25°C in the dark.

*P. sojae* P6497 wild-type (WT) and three independent mutants with verified deletion of the gene encoding *P. sojae*_482953 were tested for their ability to utilise xyloglucan as a sole carbon source. Hyphal plugs were incubated on minimal agar containing 1% (w/v) xyloglucan at 25°C (for 7 days, in the dark), after which colony morphology pictures were taken.

Mass Spectrometry was utilised to investigate the differences in degradation products released from xyloglucan breakdown by *P. sojae* WT and the gene deletion strains. Hyphal plugs of *P. sojae* WT and three independent gene deletion mutants were incubated in liquid minimal media with 1% (w/v) xyloglucan as the sole carbon source (for 7 days, in the dark). At 0 and 7 days, 60 µL of each sample was removed into a sterile 1.5 mL tube, and processed as described above. MALDI data was acquired and analysed as described above.

## Accession numbers

Sequence data can be found in the NCBI protein database under the following accession numbers: *P. sojae*_482953 (PHYSODRAFT_482953; EGZ25667.1), *P. sojae*_247788 (PHYSODRAFT_247788; EGZ25668.1), *P. sojae*_559651 (PHYSODRAFT_559651; **PsXEG1**; EGZ16757.1), *P. sojae*_355355 (PHYSODRAFT_355355; EGZ11358.1), *P. sojae*_260883 (PHYSODRAFT_260883; EGZ11355.1), *P. sojae*_520924 (PHYSODRAFT_520924; EGZ11360.1), *P. sojae*_520599 (PHYSODRAFT_520599; EGZ11361.1), *P. sojae*_360375 (PHYSODRAFT_360375; **PsXLP1**; EGZ16758.1), *P. sojae*_338074 (PHYSODRAFT_338074; EGZ11362.1), *P. sojae*_338064 (PHYSODRAFT_338064; EGZ11350.1), *P. sojae*_520248 (PHYSODRAFT_520248; EGZ11363.1), *P. nicotianae*_extension1 (AM587_10010125; KUF85461.1), *P. nicotianae*_extension2 (AM587_10010124; KUF85460.1), *P. nicotianae*_binding (AM587_10012326; KUF76577.1), *P. cactorum*_extension1 (PC111_g4312; KAG2838309.1), *P. cactorum*_extension2 (Pcac1_g9740; KAG2780169.1), and *P cactorum*_binding (Pcac_g2817; KAG2788088.1).

## Data Access

The sequence alignments and underlying data for all plots are available at: https://figshare.com/s/504fb93cc98bffb481de (DOI: 10.6084/m9.figshare.21524886).

## Acknowledgements

VA and this research was supported by a Philip Leverhulme Award (PLP-2014-147) to TAR. TAR is supported by a Royal Society University Research Fellowship (URF/R/191005). XY is supported by an award from The Gatsby Charitable Foundation and Biological Sciences Research Council (BBSRC) BBS/E/J/000PR9797 to N.J.T. Gene deletion studies by YF and JH were supported by NIH/NIAID R01 AI039115-26.

**The authors declare no conflict of interest**

## Notes

https://figshare.com/s/504fb93cc98bffb481de

### Competing Interest Statement

The authors have declared no competing interest.

## Bibliography

1. G. Van der Auwera, et al., The phylogeny of the Hyphochytriomycota as deduced from ribosomal RNA sequences of Hyphochytrium catenoides. Mol. Biol. Evol. 12, 671–678 (1995).

2. S. L. Baldauf, The Deep Roots of Eukaryotes. Science 300, 1703–1706 (2003).

3. T. Cavalier-Smith, E. E.-Y. Chao, Phylogeny and Megasystematics of Phagotrophic Heterokonts (Kingdom Chromista). J. Mol. Evol. 62, 388–420 (2006).

4. I. Riisberg, et al., Seven Gene Phylogeny of Heterokonts. Protist 160, 191–204 (2009).

5. G. W. Beakes, M. Thines, “Hyphochytriomycota and Oomycota” in Handbook of the Protists, J. M. Archibald, A. G. B. Simpson, C. H. Slamovits, Eds. (Springer International Publishing, 2017), pp. 435–505.

6. T. A. Richards, N. J. Talbot, Horizontal gene transfer in osmotrophs: playing with public goods. Nat. Rev. Microbiol. 11, 720–727 (2013).

7. Ainsworth, Bisby, Dictionary of the Fungi. Nature 191, 109–109 (1961).

8. H. Förster, M. O. Coffey, H. Elwood, M. L. Sogin, Sequence Analysis of the Small Subunit Ribosomal Rnas of Three Zoosporic Fungi and Implications for Fungal Evolution. Mycologia 82, 306–312 (1990).

9. M. C. Leclerc, J. Guillot, M. Deville, Taxonomic and phylogenetic analysis of Saprolegniaceae (Oomycetes) inferred from LSU rDNA and ITS sequence comparisons. Antonie Van Leeuwenhoek 77, 369–377 (2000).

10. D. S. S. Hudspeth, S. A. Nadler, M. E. S. Hudspeth, A COX2 molecular phylogeny of the Peronosporomycetes. Mycologia 92, 674–684 (2000).

11. D. S. S. Hudspeth, D. Stenger, M. E. S. Hudspeth, A cox2 phylogenetic hypothesis for the downy mildews and white rusts. Fungal Divers., 11 (2003).

12. M. Thines, Characterisation and phylogeny of repeated elements giving rise to exceptional length of ITS2 in several downy mildew genera (Peronosporaceae). Fungal Genet. Biol. 44, 199– 207 (2007).

13. C. G. P. McCarthy, D. A. Fitzpatrick, Phylogenomic Reconstruction of the Oomycete Phylogeny Derived from 37 Genomes. mSphere 2, e00095–17 (2017).

14. M. Latijnhouwers, P. J. G. M. de Wit, F. Govers, Oomycetes and fungi: similar weaponry to attack plants. Trends Microbiol. 11, 462–469 (2003).

15. N. P. Money, C. M. Davis, J. P. Ravishankar, Biomechanical evidence for convergent evolution of the invasive growth process among fungi and oomycete water molds. Fungal Genet. Biol. 41, 872–876 (2004).

16. L. P. N. M. Kroon, H. Brouwer, A. W. A. M. de Cock, F. Govers, The Genus Phytophthora Anno 2012. Phytopathology® 102, 348–364 (2012).

17. M. McNeil, A. Darvill, S. Fry, P. Albersheim, Structure and Function of the Primary Cell Walls of Plants. Annu. Rev. Biochem. 53, 625–63 (1984).

18. T. M. Schindler, The new view of the primary cell wall. Z. Für Pflanzenernähr. Bodenkd. 161, 499–508 (1998).

19. W. S. York, H. van Halbeek, A. G. Darvill, P. Albersheim, Structural analysis of xyloglucan oligosaccharides by 1H-n.m.r. spectroscopy and fast-atom-bombardment mass spectrometry. Carbohydr. Res. 200, 9–31 (1990).

20. W. S. York, V. S. Kumar Kolli, R. Orlando, P. Albersheim, A. G. Darvill, The structures of arabinoxyloglucans produced by solanaceous plants. Carbohydr. Res. 285, 99–128 (1996).

21. J. P. Vincken, W. S. York, G. Beldman, A. G. Voragen, Two general branching patterns of xyloglucan, XXXG and XXGG. Plant Physiol. 114, 9–13 (1997).

22. M. Hoffman, et al., Structural analysis of xyloglucans in the primary cell walls of plants in the subclass Asteridae. Carbohydr. Res. 340, 1826–1840 (2005).

23. M. J. Peña, A. G. Darvill, S. Eberhard, W. S. York, M. A. O’Neill, Moss and liverwort xyloglucans contain galacturonic acid and are structurally distinct from the xyloglucans synthesized by hornworts and vascular plants*. Glycobiology 18, 891–904 (2008).

24. S. C. Fry, et al., An unambiguous nomenclature for xyloglucan-derived oligosaccharides. Physiol. Plant. 89, 1–3 (1993).

25. J. Claverie, et al., The Cell Wall-Derived Xyloglucan Is a New DAMP Triggering Plant Immunity in Vitis vinifera and Arabidopsis thaliana. Front. Plant Sci. 9 (2018).

26. C. A. Lévesque, et al., Genome sequence of the necrotrophic plant pathogen Pythium ultimum reveals original pathogenicity mechanisms and effector repertoire. Genome Biol. 11, R73 (2010).

27. M. M. Zerillo, et al., Carbohydrate-Active Enzymes in Pythium and Their Role in Plant Cell Wall and Storage Polysaccharide Degradation. PLOS ONE 8, e72572 (2013).

28. T. A. Torto, L. Rauser, S. Kamoun, The pipg1 gene of the oomycete Phytophthora infestans encodes a fungal-like endopolygalacturonase. Curr. Genet. 40, 385–390 (2002).

29. T. A. Richards, J. B. Dacks, J. M. Jenkinson, C. R. Thornton, N. J. Talbot, Evolution of Filamentous Plant Pathogens: Gene Exchange across Eukaryotic Kingdoms. Curr. Biol. 16, 1857–1864 (2006).

30. L. Belbahri, G. Calmin, F. Mauch, J. O. Andersson, Evolution of the cutinase gene family: Evidence for lateral gene transfer of a candidate Phytophthora virulence factor. Gene 408, 1–8 (2008).

31. T. A. Richards, et al., Horizontal gene transfer facilitated the evolution of plant parasitic mechanisms in the oomycetes. Proc. Natl. Acad. Sci. 108, 15258–15263 (2011).

32. I. Misner, N. Blouin, G. Leonard, T. A. Richards, C. E. Lane, The Secreted Proteins of Achlya hypogyna and Thraustotheca clavata Identify the Ancestral Oomycete Secretome and Reveal Gene Acquisitions by Horizontal Gene Transfer. Genome Biol. Evol. 7, 120–135 (2015).

33. F. Savory, G. Leonard, T. A. Richards, The Role of Horizontal Gene Transfer in the Evolution of the Oomycetes. PLOS Pathog. 11, e1004805 (2015).

34. B. A. Kronmiller, et al., Comparative Genomic Analysis of 31 Phytophthora Genomes Reveals Genome Plasticity and Horizontal Gene Transfer. Mol. Plant-Microbe Interact. MPMI 36, 26–46 (2023).

35. D. Soanes, T. Richards, Horizontal Gene Transfer in Eukaryotic Plant Pathogens. Annu. Rev. Phytopathol. 52, 583–614 (2014).

36. R. P. Hirt, C. Alsmark, T. M. Embley, Lateral gene transfers and the origins of the eukaryote proteome: a view from microbial parasites. Curr. Opin. Microbiol. 23, 155–162 (2015).

37. F. R. Savory, D. S. Milner, D. C. Miles, T. A. Richards, Ancestral Function and Diversification of a Horizontally Acquired Oomycete Carboxylic Acid Transporter. Mol. Biol. Evol. 35, 1887–1900 (2018).

38. B. N. Adhikari, et al., Comparative Genomics Reveals Insight into Virulence Strategies of Plant Pathogenic Oomycetes. PLOS ONE 8, e75072 (2013).

39. J. McGowan, D. A. Fitzpatrick, Genomic, Network, and Phylogenetic Analysis of the Oomycete Effector Arsenal. mSphere 2, e00408–17 (2017).

40. S. Ohno, Evolution by Gene Duplication (Springer Science & Business Media, 2013).

41. A. Stoltzfus, On the Possibility of Constructive Neutral Evolution. J. Mol. Evol. 49, 169–181 (1999).

42. A. Force, et al., Preservation of Duplicate Genes by Complementary, Degenerative Mutations. Genetics 151, 1531–1545 (1999).

43. Z. Ma, et al., A paralogous decoy protects Phytophthora sojae apoplastic effector PsXEG1 from a host inhibitor. Science 355, 710–714 (2017).

44. Z. Ma, et al., A Phytophthora sojae Glycoside Hydrolase 12 Protein Is a Major Virulence Factor during Soybean Infection and Is Recognized as a PAMP[OPEN]. Plant Cell 27, 2057–2072 (2015).

45. L. V. Valen, “19. A New Evolutionary Law (1973)” in 19. A New Evolutionary Law (1973), (University of Chicago Press, 2014), pp. 284–314.

46. G. Bell, The Masterpiece of Nature: The Evolution and Genetics of Sexuality (Routledge, 2019).

47. B. T. Grenfell, A. P. Dobson, H. K. Moffatt, Ecology of Infectious Diseases in Natural Populations (Cambridge University Press, 1995).

48. R. D. Finn, et al., HMMER web server: 2015 update. Nucleic Acids Res. 43, W30–W38 (2015).

49. R. D. Finn, et al., The Pfam protein families database: towards a more sustainable future. Nucleic Acids Res. 44, D279–D285 (2016).

50. M. Blum, et al., The InterPro protein families and domains database: 20 years on. Nucleic Acids Res. 49, D344–D354 (2021).

51. J. E. Stajich, et al., FungiDB: an integrated functional genomics database for fungi. Nucleic Acids Res. 40, D675–D681 (2012).

52. E. Y. Basenko, et al., FungiDB: An Integrated Bioinformatic Resource for Fungi and Oomycetes. J. Fungi 4, 39 (2018).

53. C. J. Van Den Bergh, A. C. A. P. A. Bekkers, P. De Geus, H. M. Verheij, G. H. De Haas, Secretion of biologically active porcine prophospholipase A2 by Saccharomyces cerevisiae. Eur. J. Biochem. 170, 241–246 (1987).

54. G. L. Miller, Use of Dinitrosalicylic Acid Reagent for Determination of Reducing Sugar. ACS Publ. (2002) 10.1021/ac60147a030 (October 10, 2022).

55. E. R. Master, Y. Zheng, R. Storms, A. Tsang, J. Powlowski, A xyloglucan-specific family 12 glycosyl hydrolase from Aspergillus niger: recombinant expression, purification and characterization. Biochem. J. 411, 161–170 (2008).

56. L. A. Kelley, M. J. E. Sternberg, Protein structure prediction on the Web: a case study using the Phyre server. Nat. Protoc. 4, 363–371 (2009).

57. J. Jumper, et al., Highly accurate protein structure prediction with AlphaFold. Nature 596, 583– 589 (2021).

58. M. Varadi, et al., AlphaFold Protein Structure Database: massively expanding the structural coverage of protein-sequence space with high-accuracy models. Nucleic Acids Res. 50, D439– D444 (2022).

59. M. N. Wass, L. A. Kelley, M. J. E. Sternberg, 3DLigandSite: predicting ligand-binding sites using similar structures. Nucleic Acids Res. 38, W469–W473 (2010).

60. C. Lamb, R. A. Dixon, THE OXIDATIVE BURST IN PLANT DISEASE RESISTANCE. Annu. Rev. Plant Physiol. Plant Mol. Biol. 48, 251–275 (1997).

61. G. Felix, J. D. Duran, S. Volko, T. Boller, Plants have a sensitive perception system for the most conserved domain of bacterial flagellin. Plant J. 18, 265–276 (1999).

62. L. Gómez-Gómez, G. Felix, T. Boller, A single locus determines sensitivity to bacterial flagellin in Arabidopsis thaliana. Plant J. 18, 277–284 (1999).

63. K. Kuchitsu, H. Kosaka, T. Shiga, N. Shibuya, EPR evidence for generation of hydroxyl radical triggered byN-acetylchitooligosaccharide elicitor and a protein phosphatase inhibitor in suspension-cultured rice cells. Protoplasma 188, 138–142 (1995).

64. Y. Fang, L. Cui, B. Gu, F. Arredondo, B. M. Tyler, Efficient Genome Editing in the Oomycete Phytophthora sojae Using CRISPR/Cas9. Curr. Protoc. Microbiol. 44, 21A.1.1–21A.1.26 (2017).

65. B. A. Kronmiller, et al., Comparative genomic analysis of 31 Phytophthora genomes reveal genome plasticity and horizontal gene transfer. Mol. Plant-Microbe Interactions® (2022) 10.1094/MPMI-06-22-0133-R (November 3, 2022).

66. Z. Li, et al., A C-Terminal Proline-Rich Sequence Simultaneously Broadens the Optimal Temperature and pH Ranges and Improves the Catalytic Efficiency of Glycosyl Hydrolase Family 10 Ruminal Xylanases. Appl. Environ. Microbiol. 80, 3426–3432 (2014).

67. T.-N. Wen, J.-L. Chen, S.-H. Lee, N.-S. Yang, L.-F. Shyur, A Truncated Fibrobacter succinogenes 1,3−1,4-β-d-Glucanase with Improved Enzymatic Activity and Thermotolerance. Biochemistry 44, 9197–9205 (2005).

68. Y. Xia, et al., N-glycosylation shields Phytophthora sojae apoplastic effector PsXEG1 from a specific host aspartic protease. Proc. Natl. Acad. Sci. 117, 27685–27693 (2020).

69. Y. Sun, et al., Plant receptor-like protein activation by a microbial glycoside hydrolase. Nature 610, 335–342 (2022).

70. S. F. Altschul, W. Gish, W. Miller, E. W. Myers, D. J. Lipman, Basic local alignment search tool. J. Mol. Biol. 215, 403–410 (1990).

71. N. Galtier, M. Gouy, C. Gautier, SEAVIEW and PHYLO_WIN: two graphic tools for sequence alignment and molecular phylogeny. Bioinformatics 12, 543–548 (1996).

72. R. C. Edgar, MUSCLE: multiple sequence alignment with high accuracy and high throughput. Nucleic Acids Res. 32, 1792–1797 (2004).

73. S. Guindon, O. Gascuel, A Simple, Fast, and Accurate Algorithm to Estimate Large Phylogenies by Maximum Likelihood. Syst. Biol. 52, 696–704 (2003).

74. S. Kalyaanamoorthy, B. Q. Minh, T. K. F. Wong, A. von Haeseler, L. S. Jermiin, ModelFinder: fast model selection for accurate phylogenetic estimates. Nat. Methods 14, 587–589 (2017).

75. B. Q. Minh, et al., IQ-TREE 2: New Models and Efficient Methods for Phylogenetic Inference in the Genomic Era. Mol. Biol. Evol. 37, 1530–1534 (2020).

76. I. Letunic, P. Bork, Interactive Tree Of Life (iTOL) v5: an online tool for phylogenetic tree display and annotation. Nucleic Acids Res. 49, W293–W296 (2021).

77. M. A. Larkin, et al., Clustal W and Clustal X version 2.0. Bioinformatics 23, 2947–2948 (2007).

78. F. Madeira, et al., The EMBL-EBI search and sequence analysis tools APIs in 2019. Nucleic Acids Res. 47, W636–W641 (2019).

79. A. M. Waterhouse, J. B. Procter, D. M. A. Martin, M. Clamp, G. J. Barton, Jalview Version 2—a multiple sequence alignment editor and analysis workbench. Bioinformatics 25, 1189–1191 (2009).

80. F. Teufel, et al., SignalP 6.0 predicts all five types of signal peptides using protein language models. Nat. Biotechnol. 40, 1023–1025 (2022).

81. Wood, P.J. and Weisz, J, Detection and Assay of (1-4)-(8-D-Glucanase,(1-3)-(8-D-Glucanase,(1-3)(1-4)-(3-D-Glucanase, and Xylanase Based on Complex Formation of Substrate with Congo Red’. Cereal Chem 641 *Pp*8-15 (1987).

82. Signorell, A., Aho, K., Alfons, A., Anderegg, N., Aragon, T. and Arachchige, C, DescTools: Tools for descriptive statistics. R Package Version 099 45 (2022).

83. Student, The Probable Error of a Mean. Biometrika 6, 1–25 (1908).

84. T. Nakagawa, et al., Improved Gateway binary vectors: high-performance vectors for creation of fusion constructs in transgenic analysis of plants. Biosci. Biotechnol. Biochem. 71, 2095–2100 (2007).

85. R. Höfgen, L. Willmitzer, Storage of competent cells for Agrobacterium transformation. Nucleic Acids Res. 16, 9877 (1988).

86. L. M. Shaw, et al., DirtyGenes: testing for significant changes in gene or bacterial population compositions from a small number of samples. Sci. Rep. 9, 2373 (2019).

87. S. Whelan, N. Goldman, A General Empirical Model of Protein Evolution Derived from Multiple Protein Families Using a Maximum-Likelihood Approach. Mol. Biol. Evol. 18, 691–699 (2001).

