## Supplemental Data for "Duplication and neofunctionalization of a horizontally-transferred xyloglucanase as a facet of the red queen co-evolutionary dynamic"

Supplemental Figure 1. GH12 protein sequence alignment

Supplemental Figure 2. GH12 ROS accumulation in *N. benthamiana*

Supplemental Figure 3. Flg22 and chitin-triggered ROS accumulation in *N. benthamiana*

Supplemental Figure 4. *P. sojae*_559651 (structurally-inferred additional binding site)

Supplemental Figure 5. GH12 agar plate-based enzymatic assay

Supplemental Figure 6. *P. sojae*_247788 full-length and truncated proteins (72 h)

Supplemental Figure 7. *P. sojae*_338064 gene model correction notes

Supplemental Figure 8. *P. sojae*_482953 deletion mutant (genotyping)

Supplemental Figure 9. *P. sojae*_482953 deletion mutant (xyloglucan phenotype)

Supplemental Table 1. Primers and sgRNA used to generate *P. sojae*_482953 deletion mutant

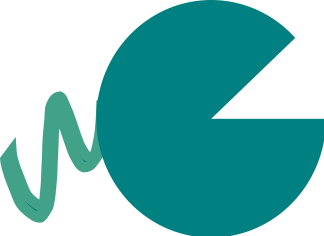

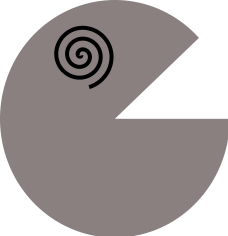

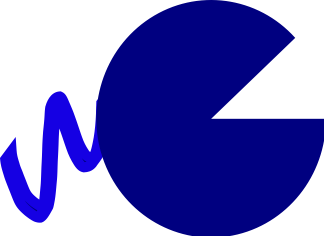

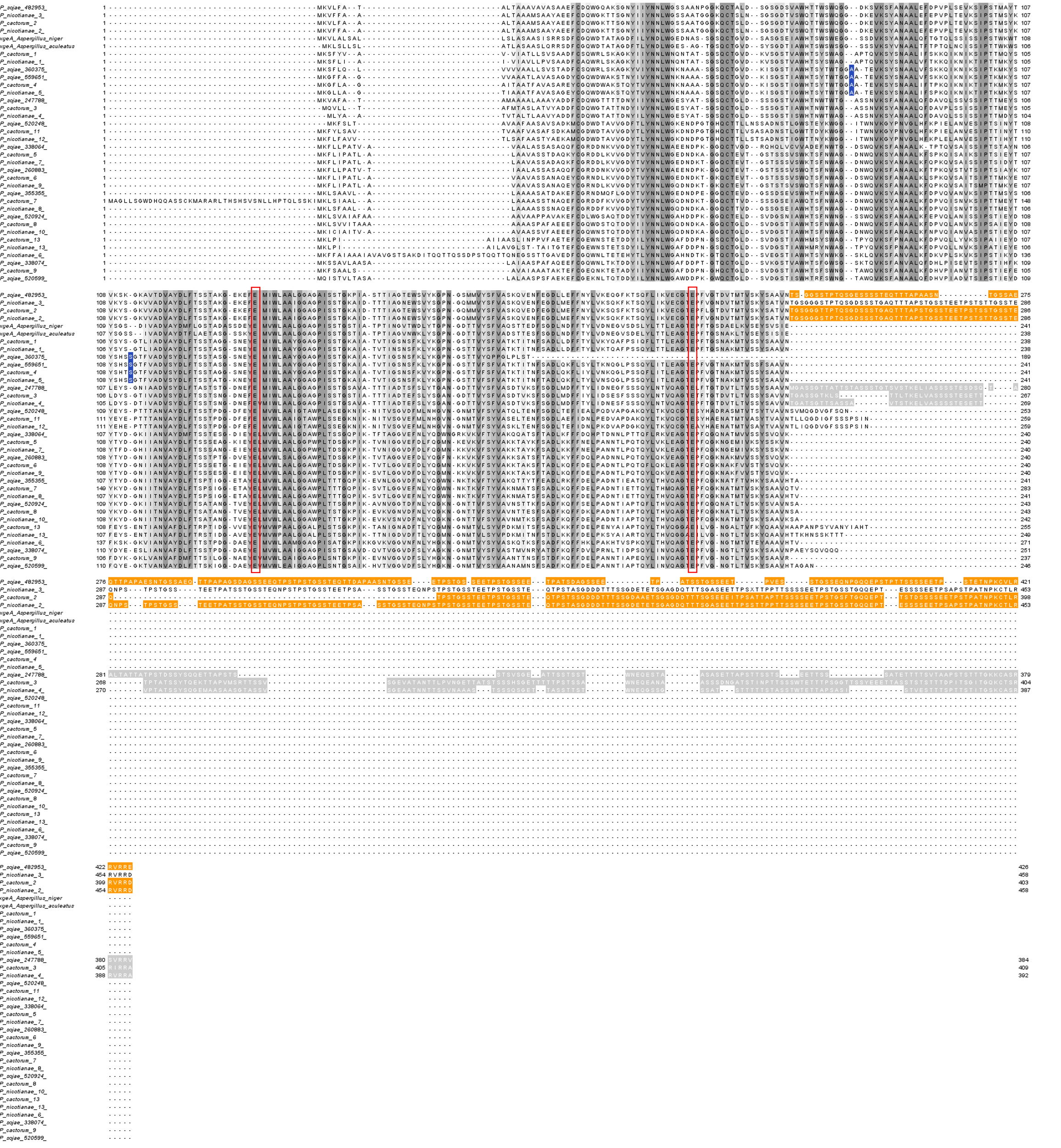

**Supplemental Figure 1.** Protein sequence alignment of GH12 proteins from *P. sojae*, *P. cactorum*, *P. nicotianae,* and fungal GH12 (xgeA) from *Aspergillus* spp. generated by Clustal Omega (70, 71) and visualized in Jalview2 (72). C-terminal extension (1) from *P. sojae*, *P. cactorum*, and *P. nicotianae* are highlighted in orange, and C-terminal extension (2) from *P. sojae*, *P. cactorum*, and *P. nicotianae* are highlighted in grey. Blue boxes highlight the positions of two indels (Alanine; A, and Serine, S), present in *P. sojae*_559651 (PsXEG1) and orthologs in *P. nicotianae* and *P. cactorum*, and *P. sojae*_360375 (PsXLP1). Red boxes highlight the positions of the two glutamic acid residues required for catalytic activity. Cartoon symbols indicate the *P. sojae* paralog with a structurally-inferred binding site (grey), and paralogs with C-terminal tails 1 and 2 (blue and green, respectively).

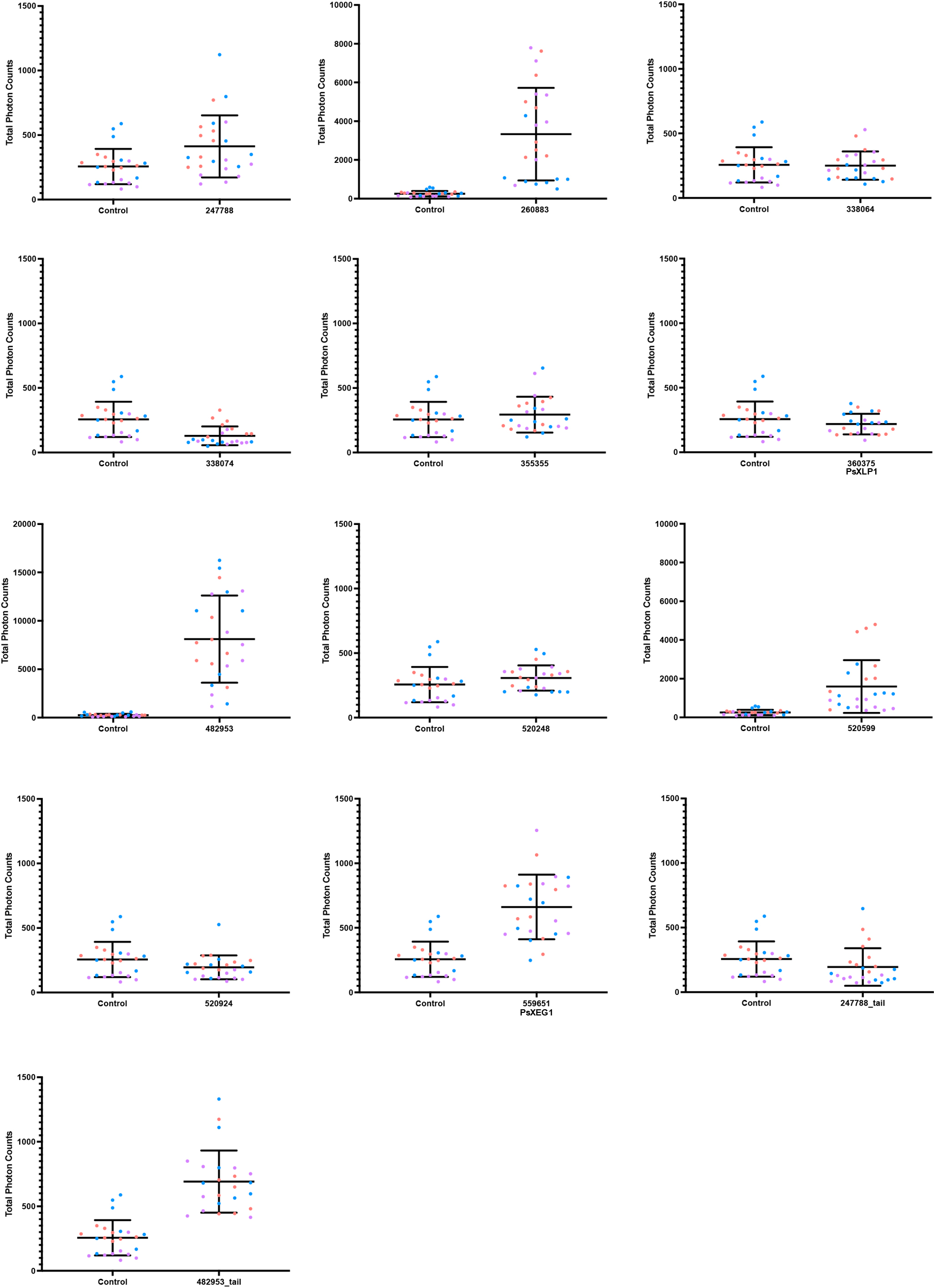

**Supplemental Figure 2**. *P. sojae* xyloglucanase paralogs were codon-optimized for expression in plants and infiltrated into the leaves of *N. benthamiana*. The dynamics of ROS production in *N. benthamiana* plants was measured from 0-60 minutes. Significant ROS accumulation was detected after infiltration with *P. sojae*_482953, 260883 and 520599. Interestingly, we find that the *P. sojae*_482953 C-terminal extension alone triggers ROS generation, but significantly reduced compared to the full-length protein, and not statistically significant from the vector-only control. Agroinfiltration of *P. sojae*_247788 and 559651 (PsXEG1) also triggered ROS generation in *N. bethamiana*, but significantly reduced compared to *P. sojae*_482953, 260883 and 520599 proteins, and total photon counts at 60 minutes were not statistically significant from the vector-only control. We did not find evidence of ROS accumulation as a result of infiltration with *P. sojae*_338064, 338074, 355355, 360375 (PsXLP1), 520248, 520924, and the *P. sojae*_247788 C-terminal extension only. Horizontal bars represent the mean and SD. The experiments were performed 3 times (orange: replicate 1, blue: replicate 2, purple: replicate 3).

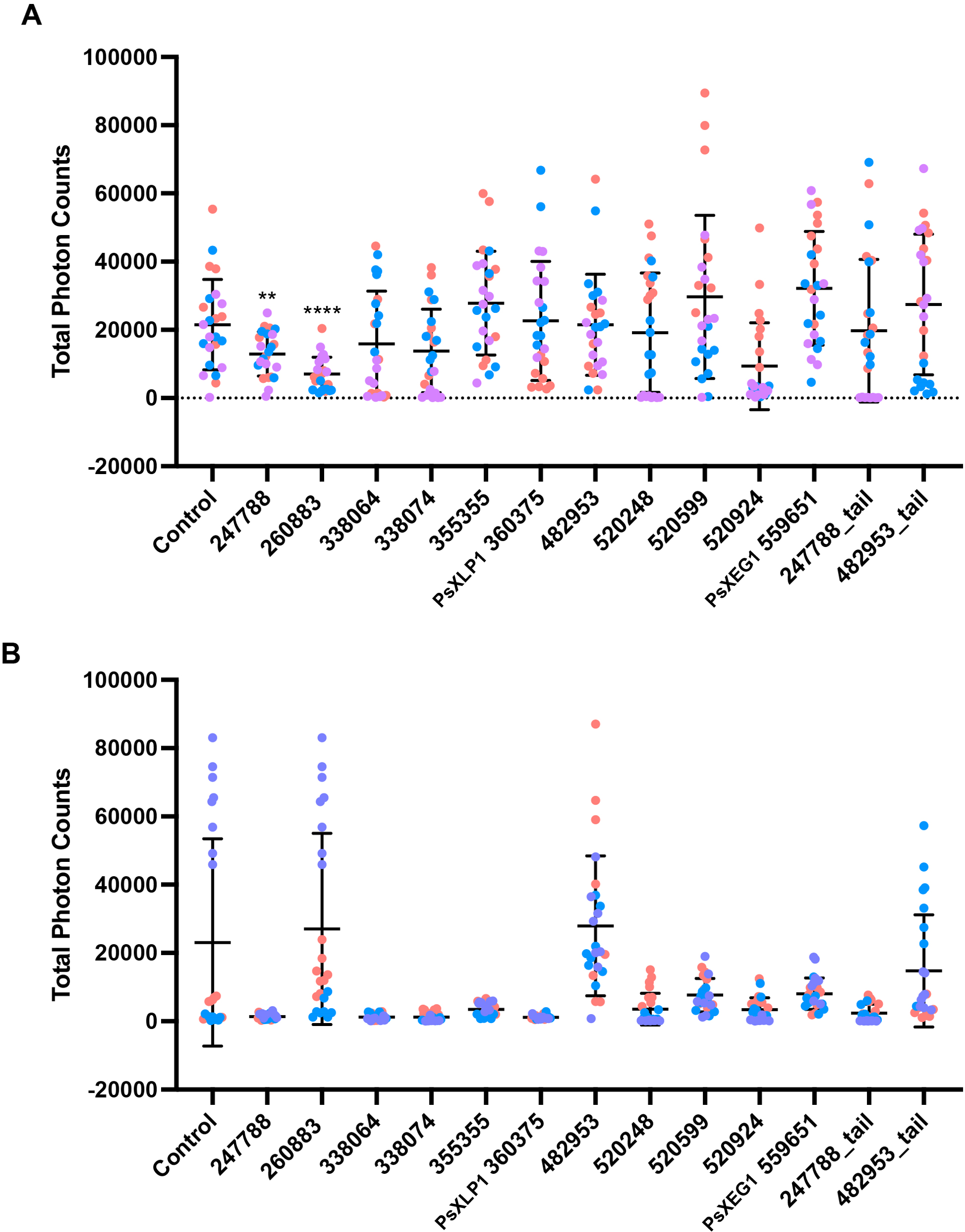

**Supplemental Figure 3.** *P. sojae* xyloglucanase paralogs were codon-optimized for expression in plants and infiltrated into the leaves of *N. benthamiana* alongside treatment with Flg22 (A) or chitin (B), representing well-known pathogen-associated molecular patterns (PAMPs) (60–62). **A.** We find that *P. sojae*_247788 and 260883 can significantly supress flg22-triggered ROS generation in *N. benthamiana.* **B.** No significant effects of the xyloglucanase variants on chitin-triggered ROS generation were observed. Horizontal bars represent the mean and SD. The experiments were performed 3 times (orange: replicate 1, blue: replicate 2, purple: replicate 3).

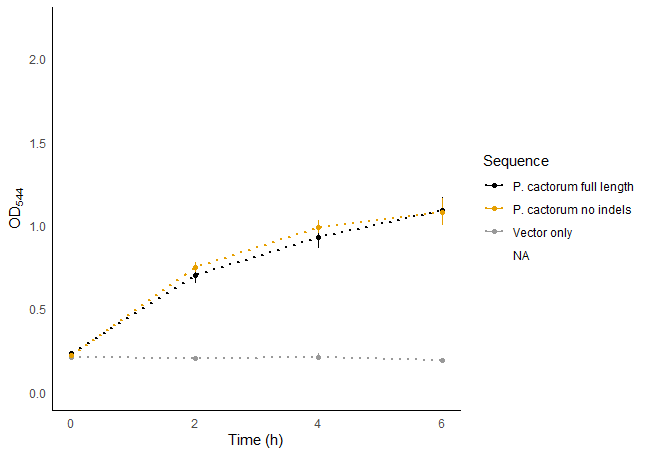

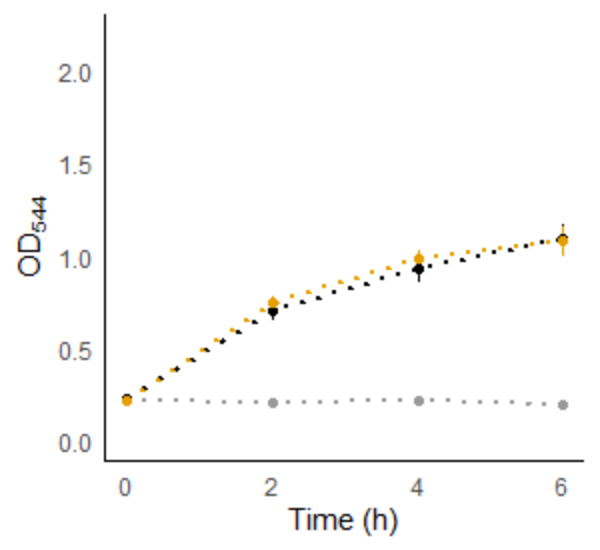

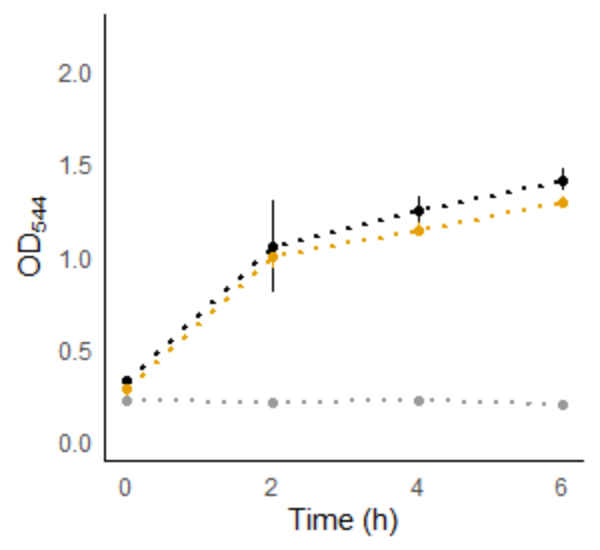

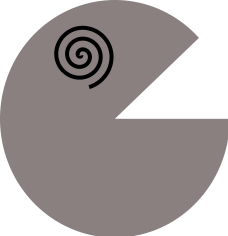

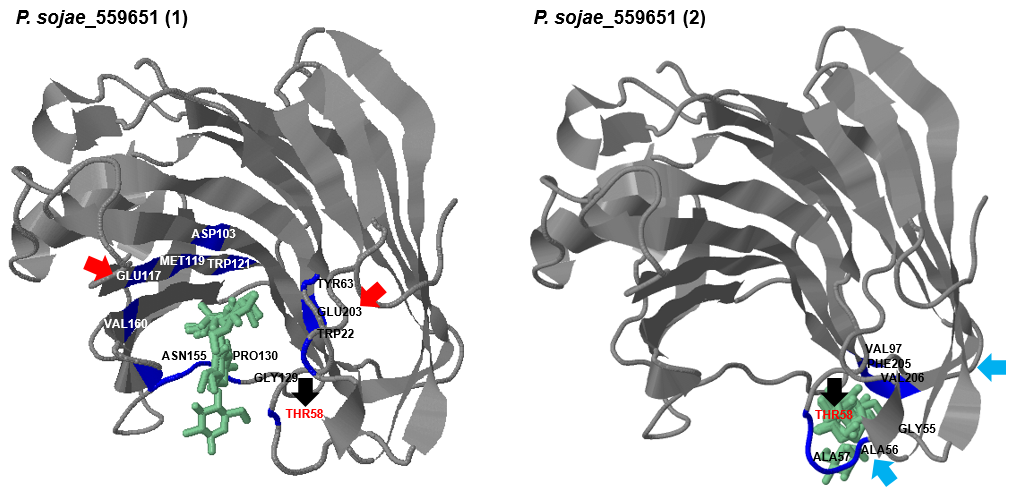

**B.**

***P. cactorum***

***P. sojae***

**A.**

Structurally-inferred binding site

**[2]**

**[1]**

**Full length**

**No indels**

**Vector only**

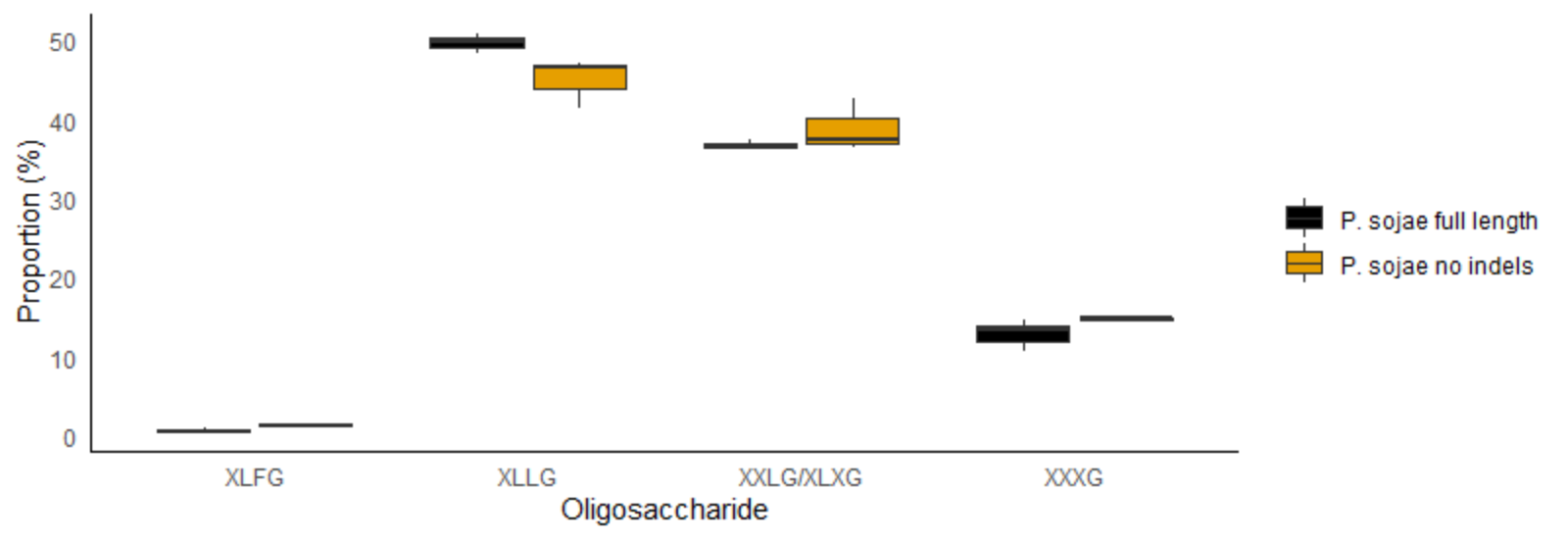

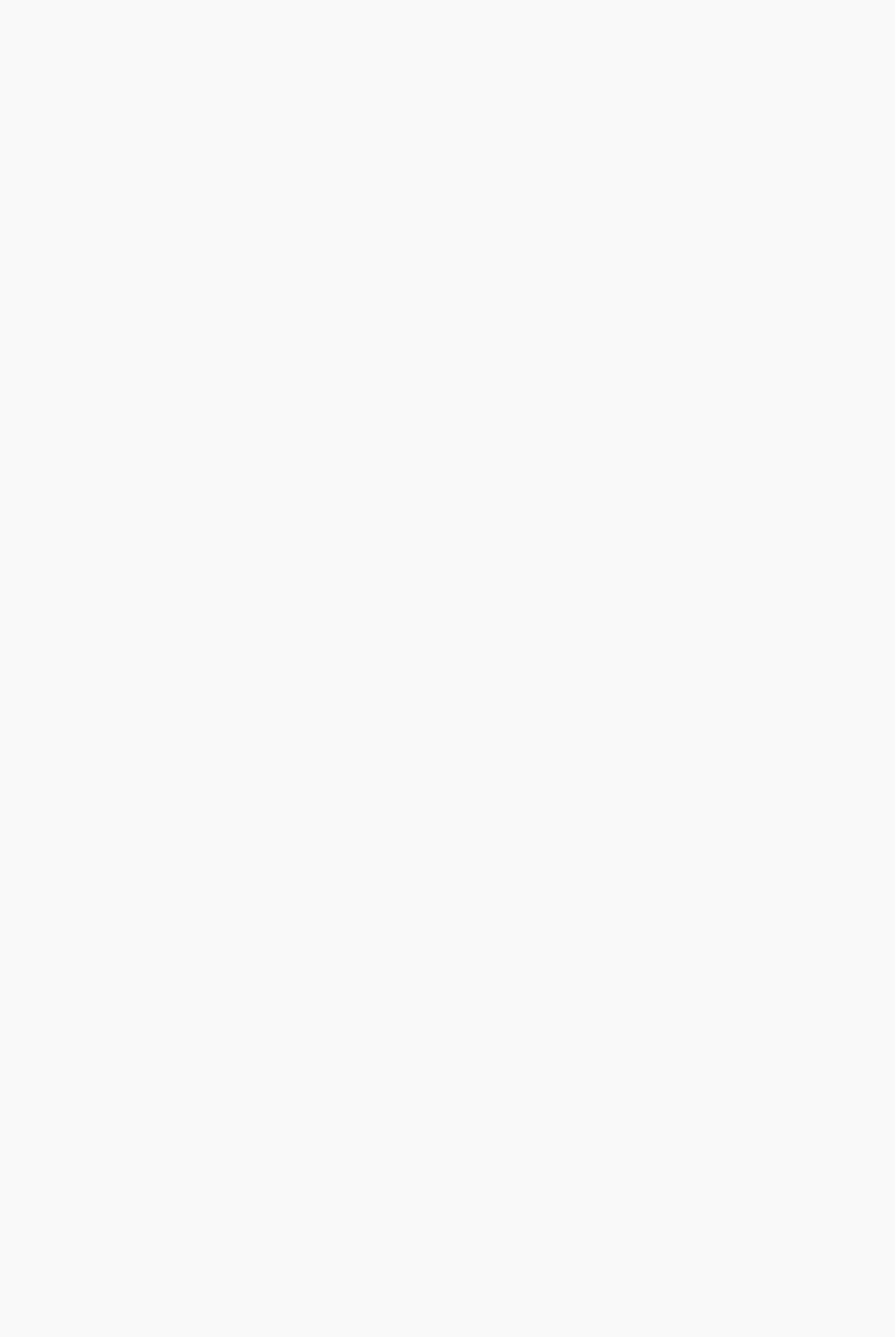

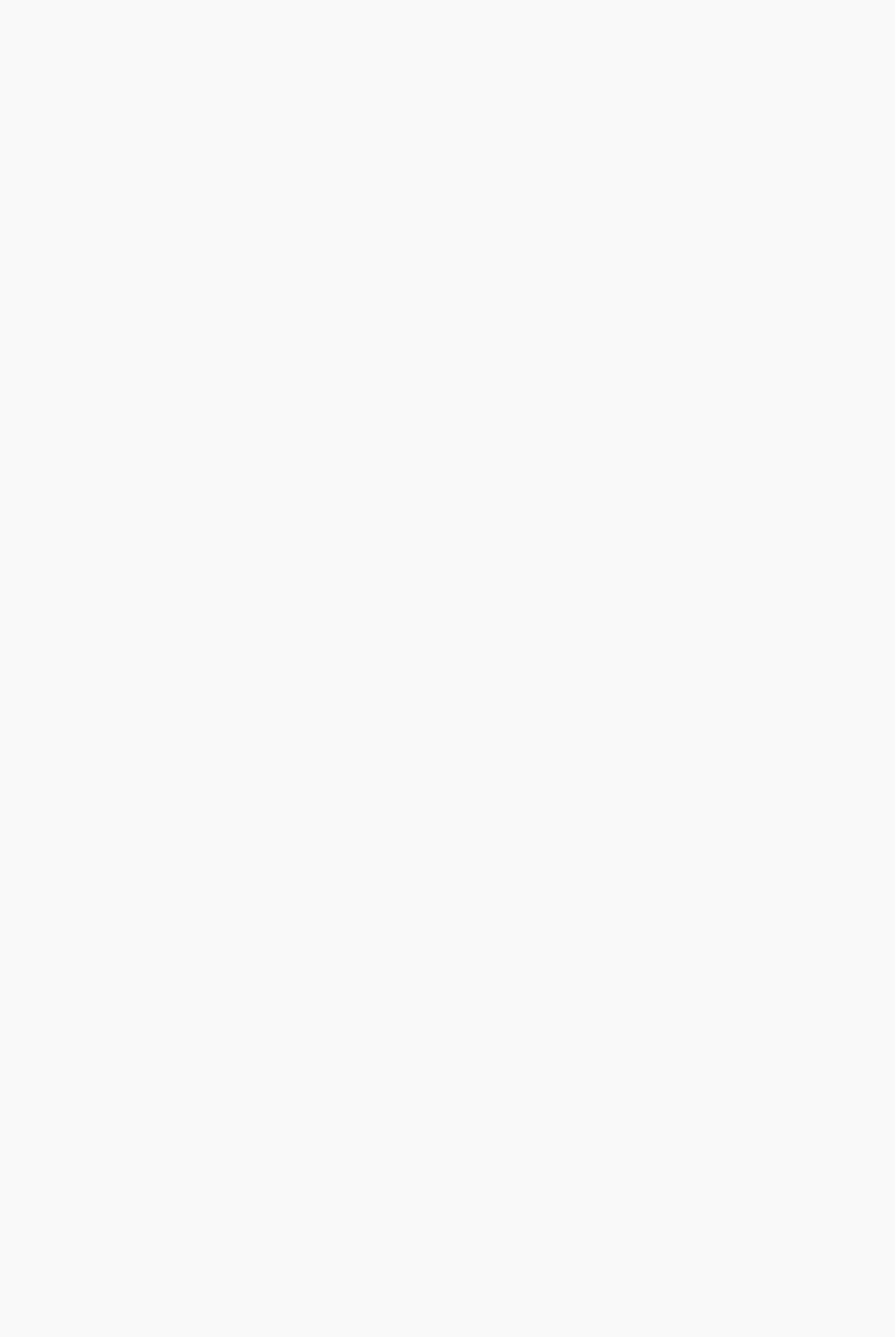

**C.**

**Supplemental Figure 4. A.** The three-dimensional structure of *P. sojae*_559651 (PsXEG1) was obtained with Phyre2 (Protein Homology/analogY Recognition Engine v2.0 (54)); predictions of substrate-binding sites (active site residues) were generated by 3DLigandSite (57), and are labelled in the figure. Known glutamic acid (catalytic) residues are shown by the red arrows. For *P. sojae*_ 559651 (PsXEG1), two substrate-binding sites were predicted by 3DLigandSite - amino acids predicted for each binding site are labelled [1] and [2]. The black arrow indicates the threonine (Thr58) residue common in both predicted sites; two putative indels (alanine and serine) that are important for the *in silico* binding site prediction for this paralog are indicated by the blue arrows. **B.** *P. sojae*_559651 (PsXEG1; full-length) and *P. sojae*_559651 (PsXEG1; no indels), and ortholog in *P. cactorum* secreted into *S. cerevisiae* culture supernatants were incubated with 1% (w/v) xyloglucan at 30°C, pH7. No significant difference in enzymatic activity was detected upon xyloglucan over 6 h. No significant reducing sugars were detected in the vector-only sample (N=3, +/- SD). **C.** *P. sojae*_559651 (PsXEG1; full-length and no indels) secreted into *S. cerevisiae* culture supernatants were incubated with 1% (w/v) xyloglucan at 30°C, pH 7, for 72 h. MALDI-MS spectra confirmed the release of xyloglucan oligosaccharides. Four peaks of interest were observed; ions with *m/z* of ~1085, 1247, 1409 and 1571 – putatively corresponding to the oligosaccharides XXXG, XXLG (or XLXG), XLLG, and XLFG respectively (24). The relative intensity of species identified at these peaks were compared by calculating the ratios between the areas under the peaks, to probe putative differences in preferential binding of the xyloglucan backbone. We find no evidence for differences in oligosaccharides released between the two protein variants.

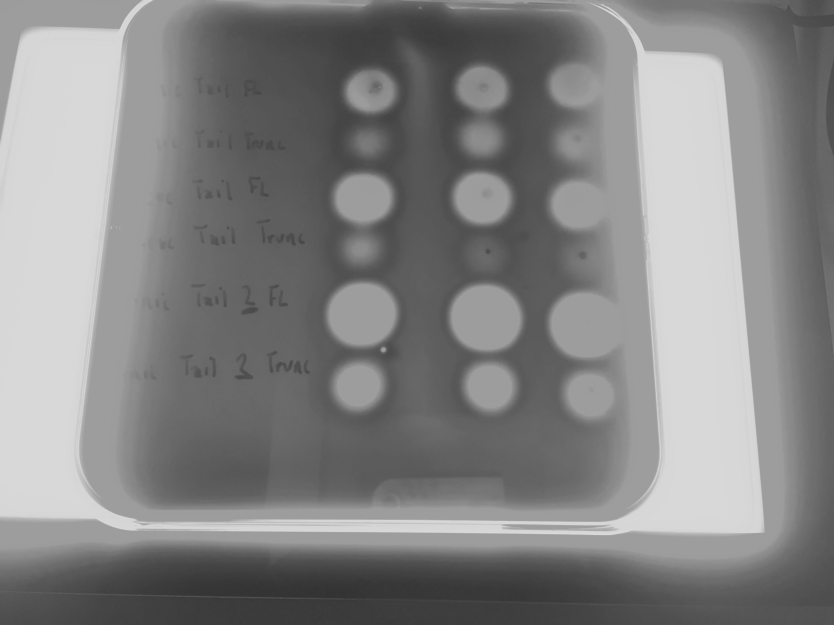

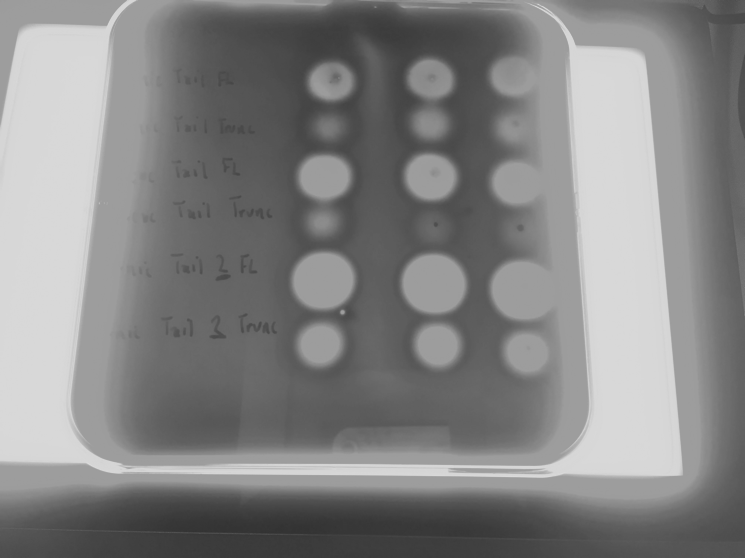

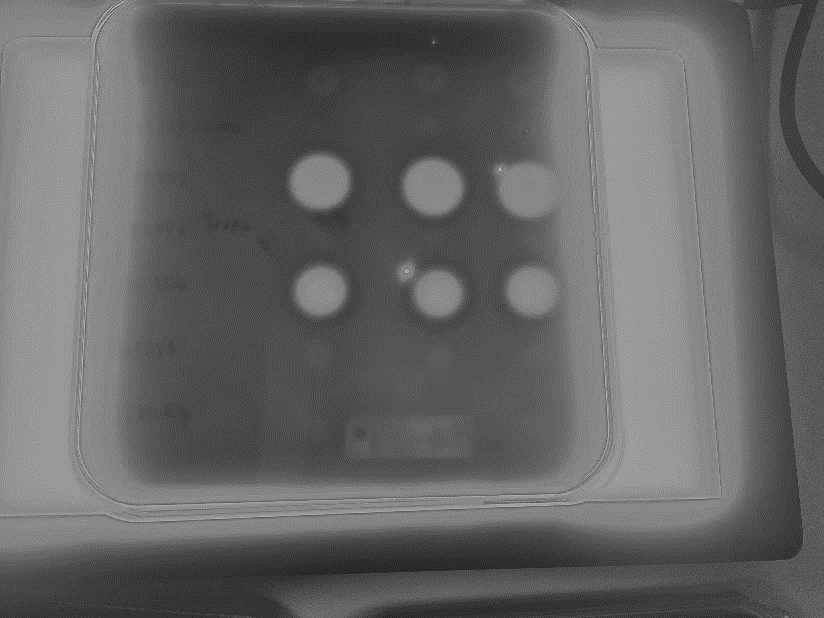

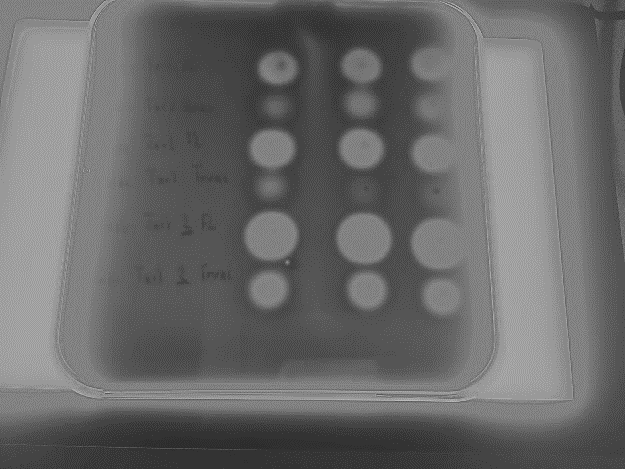

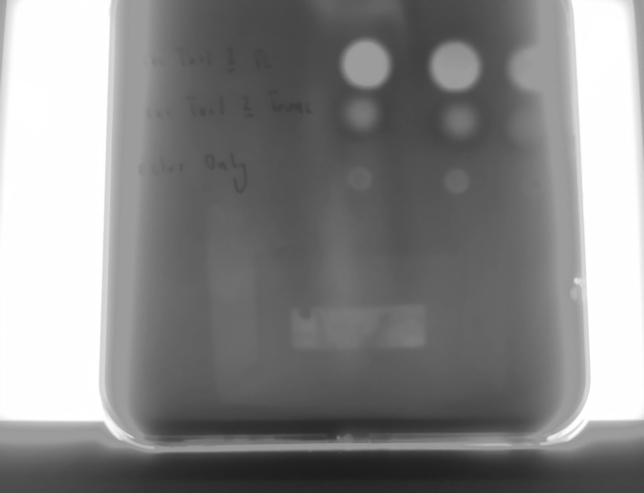

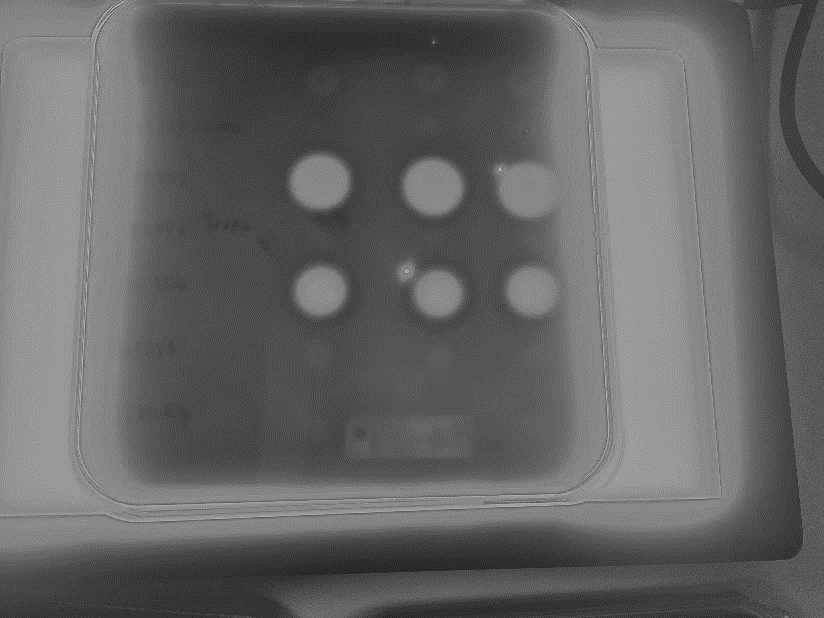

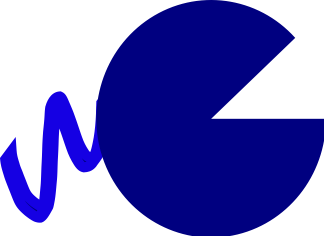

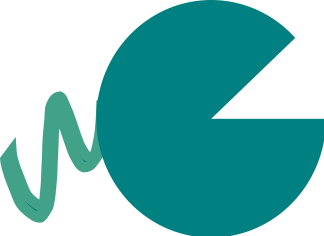

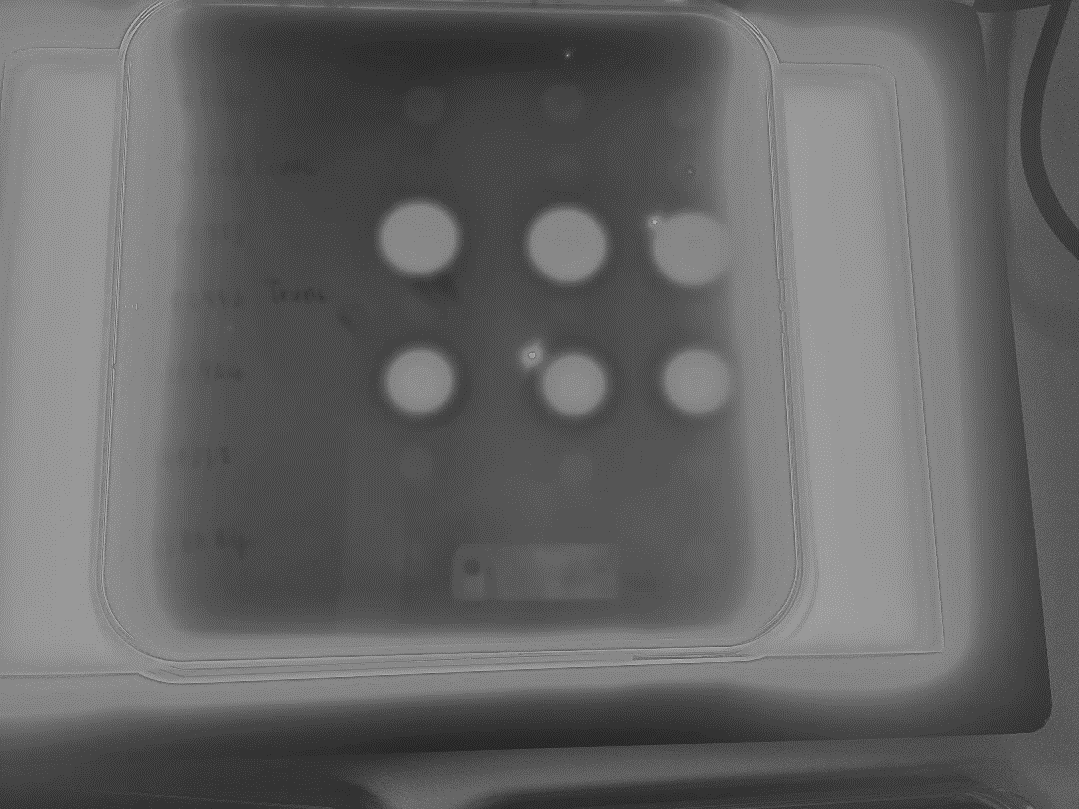

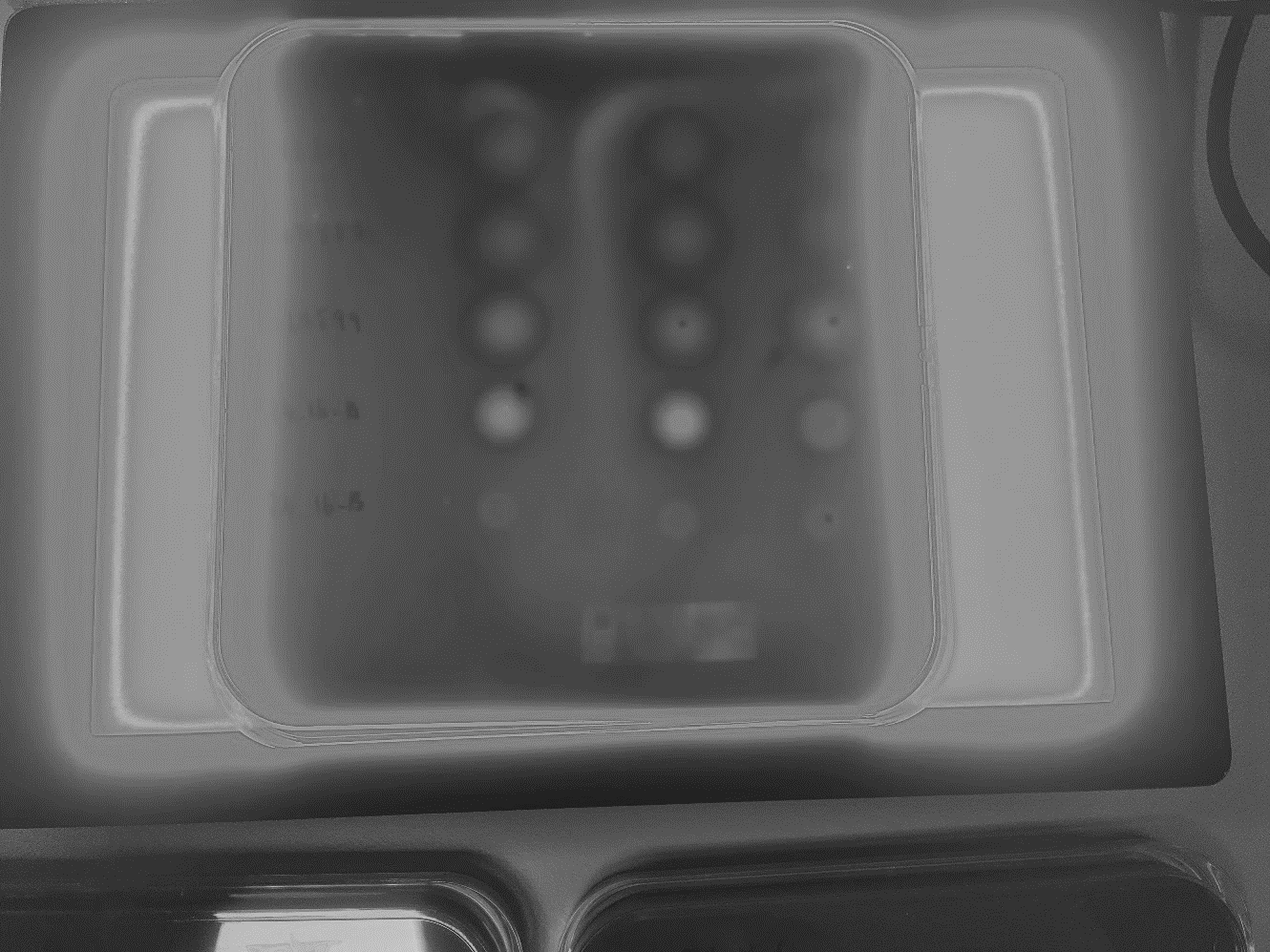

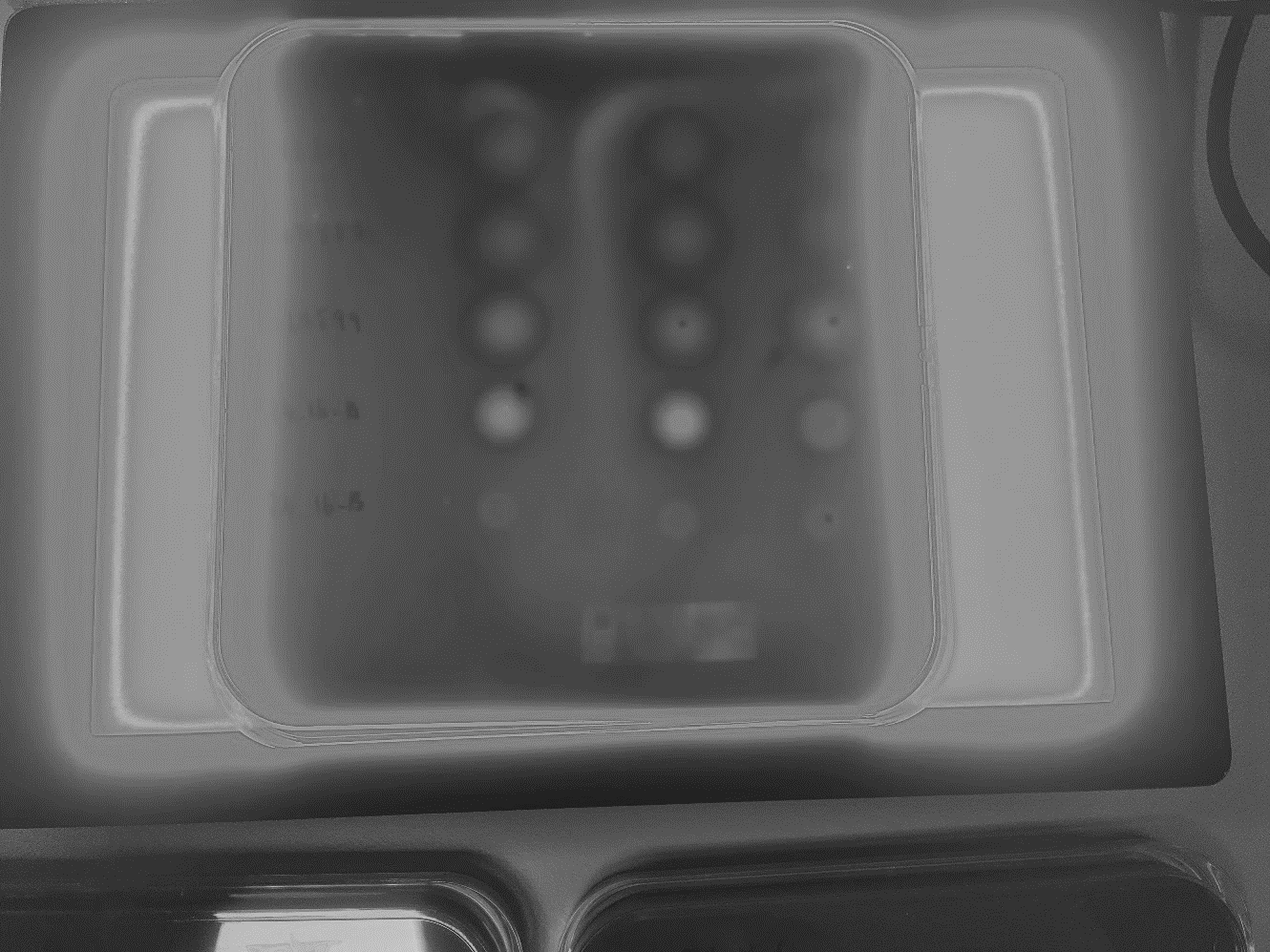

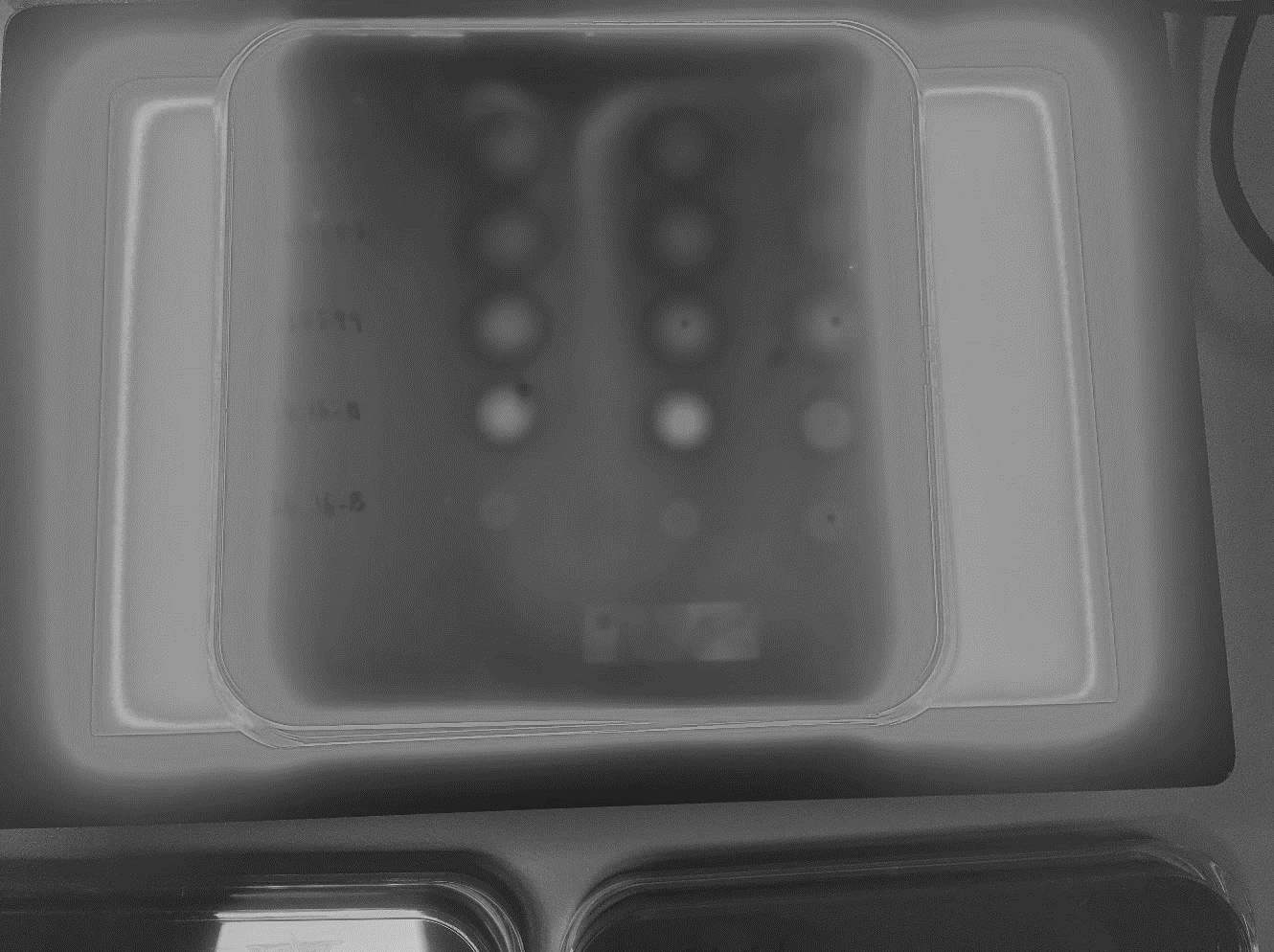

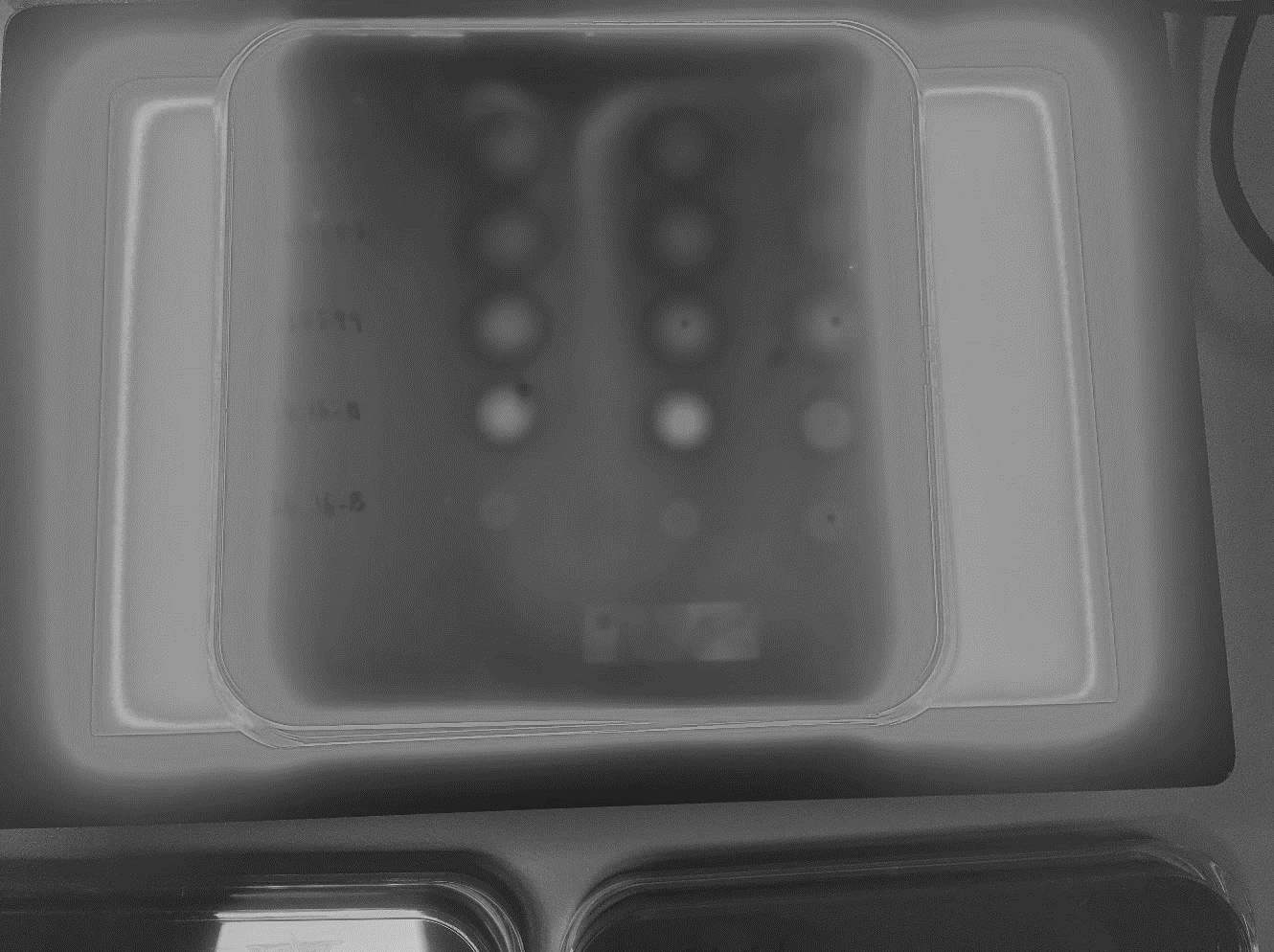

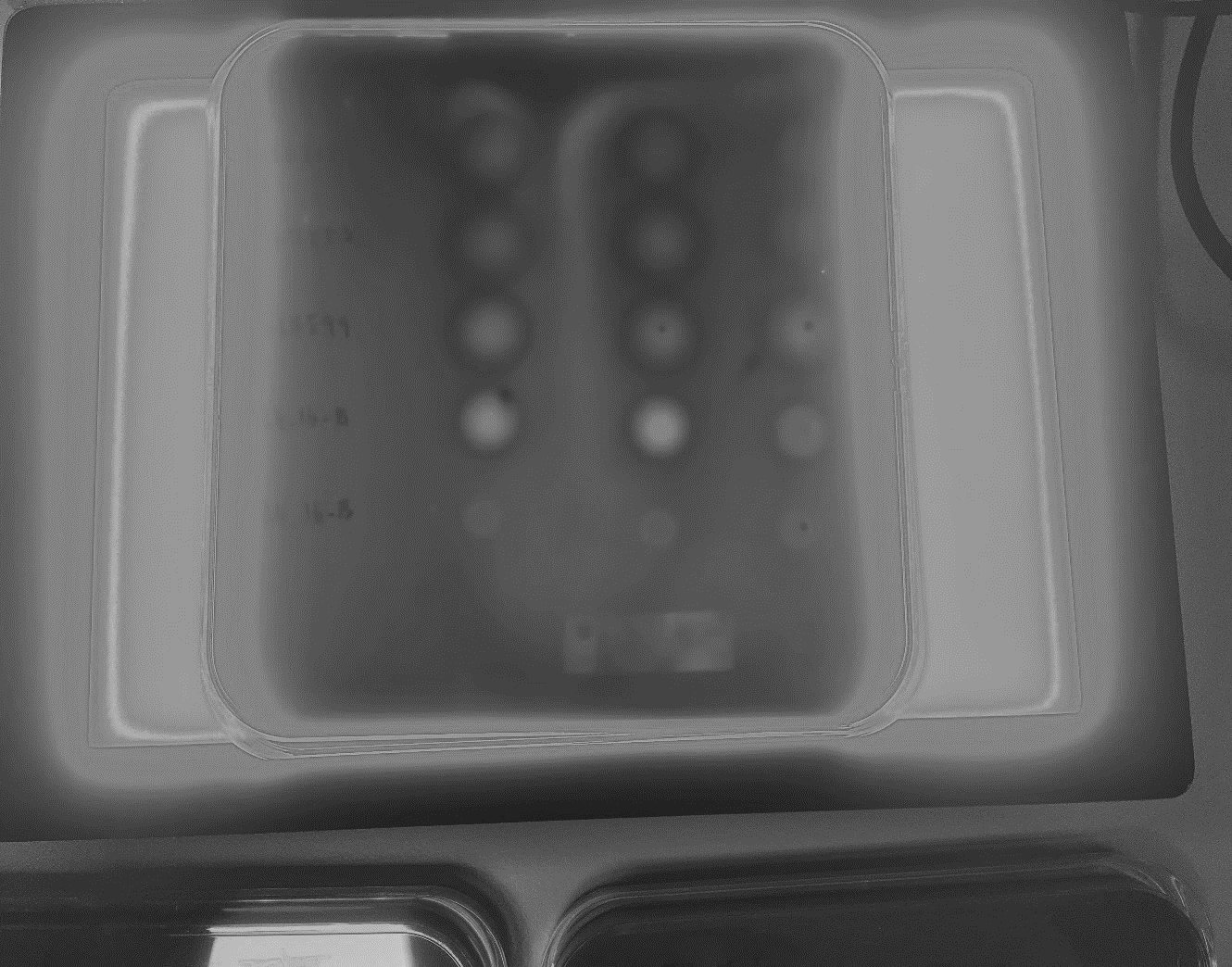

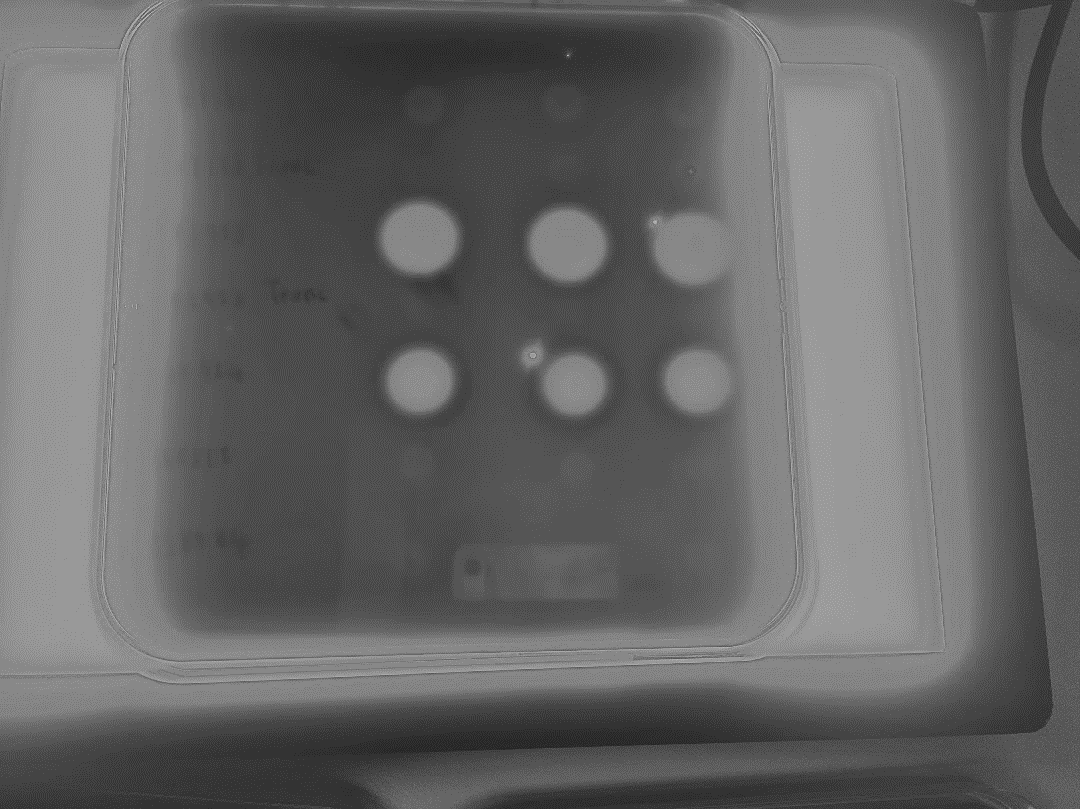

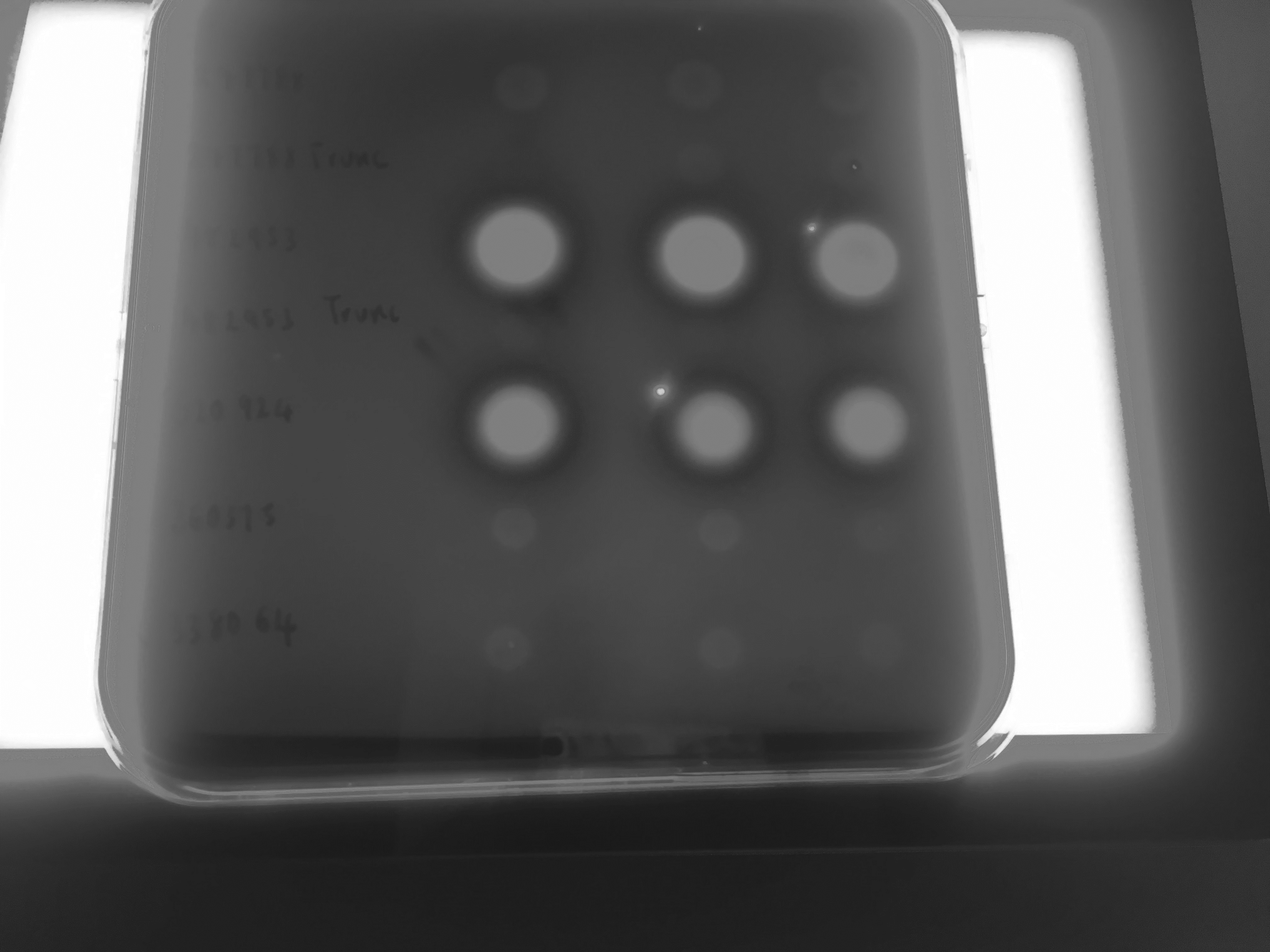

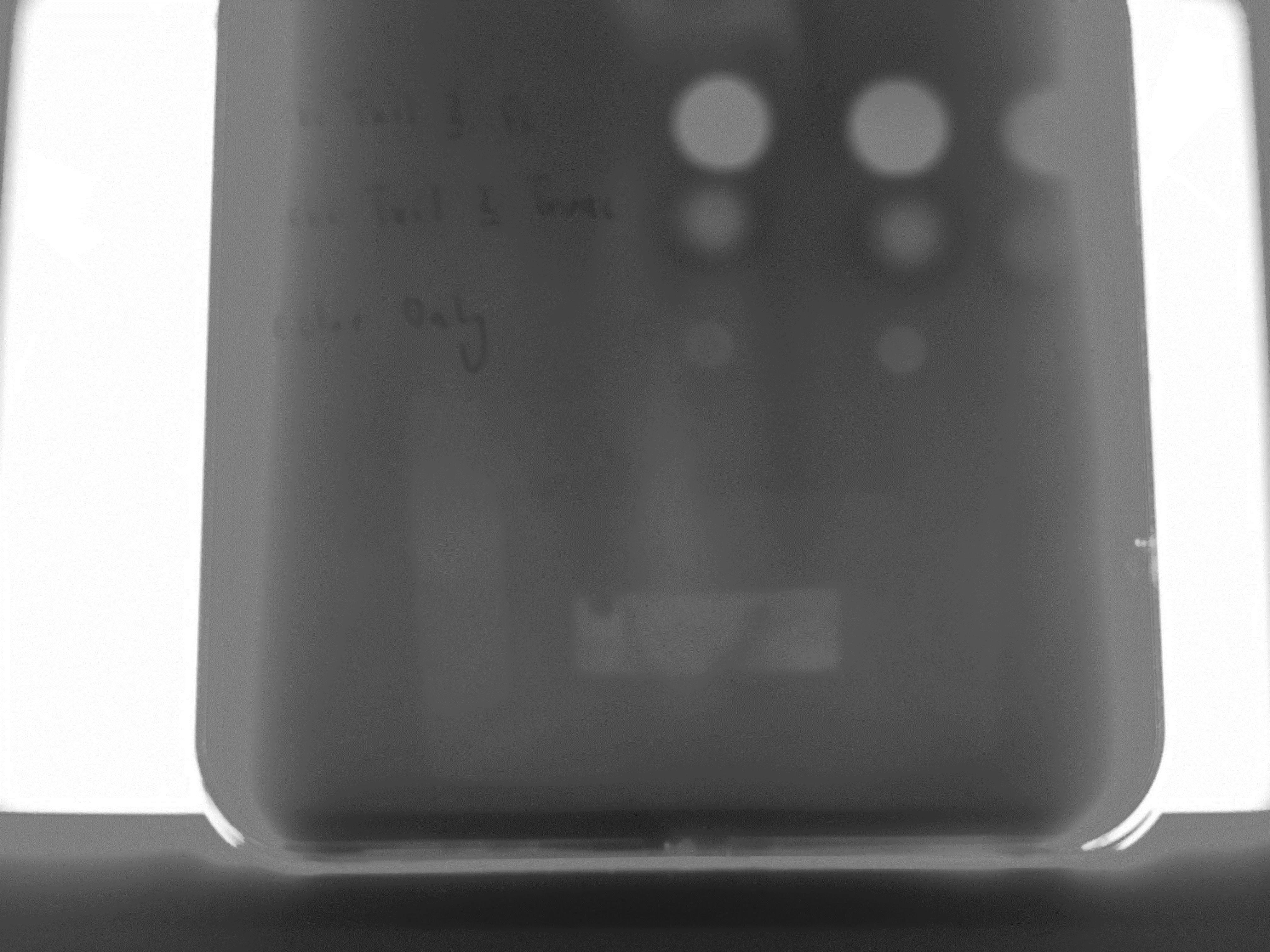

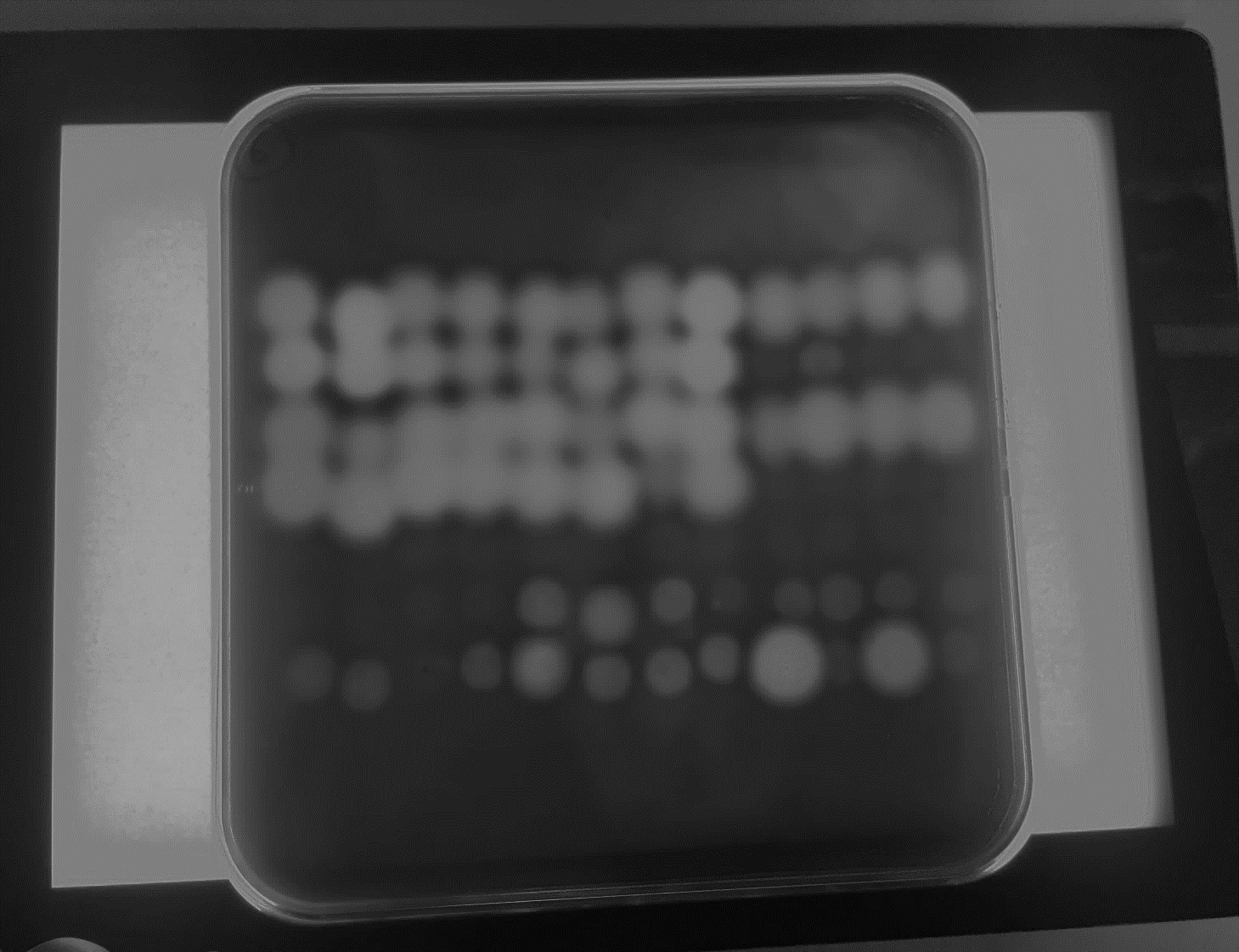

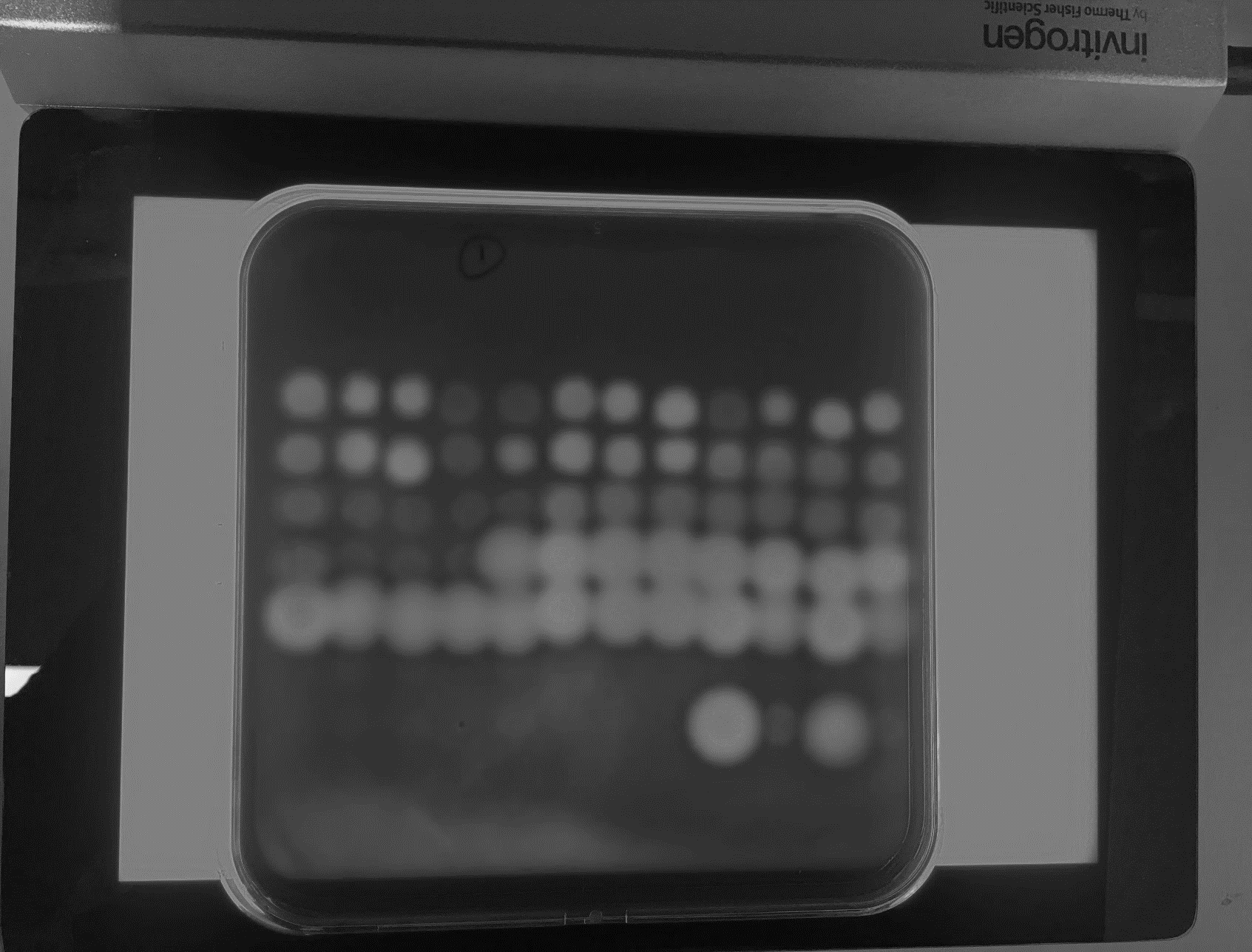

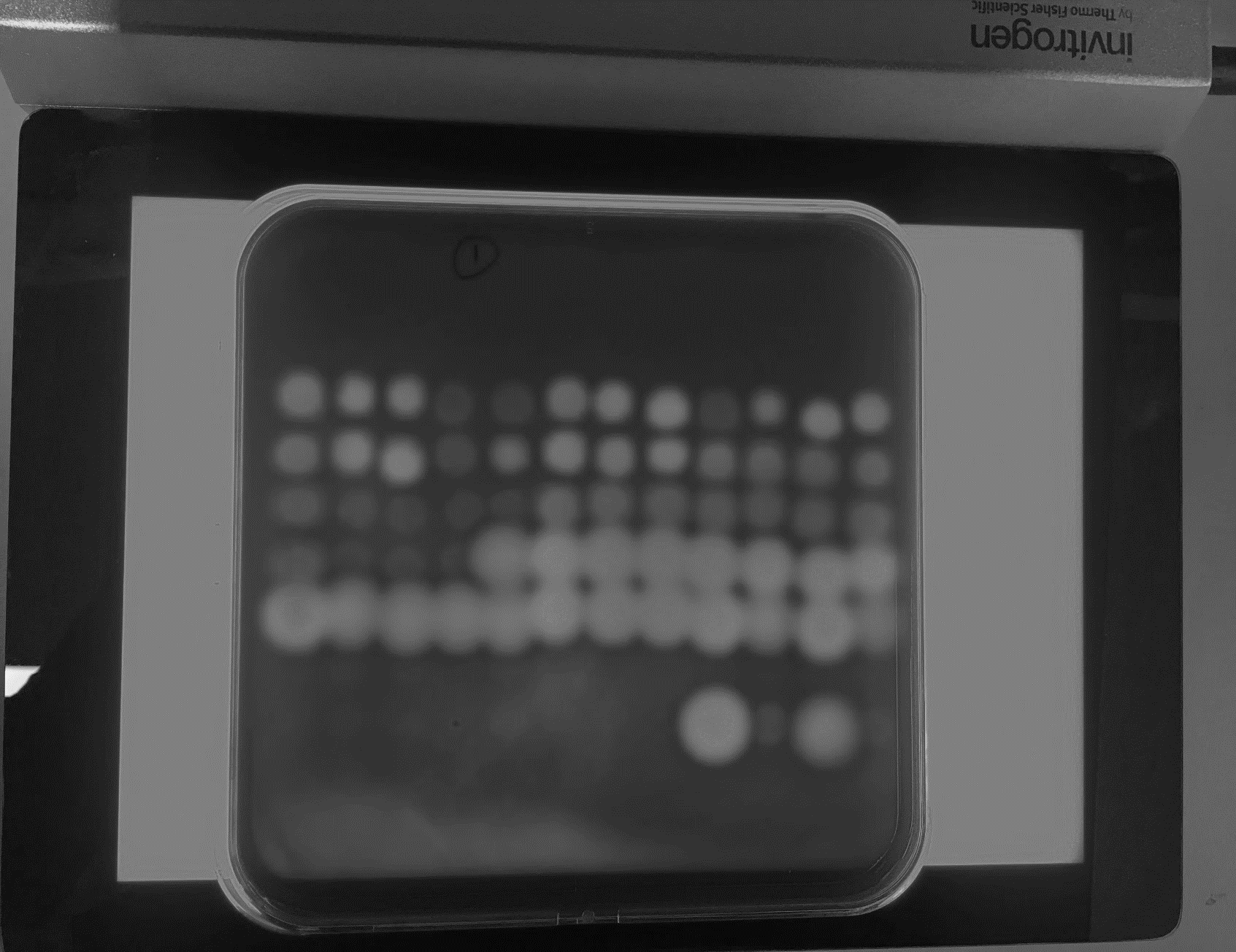

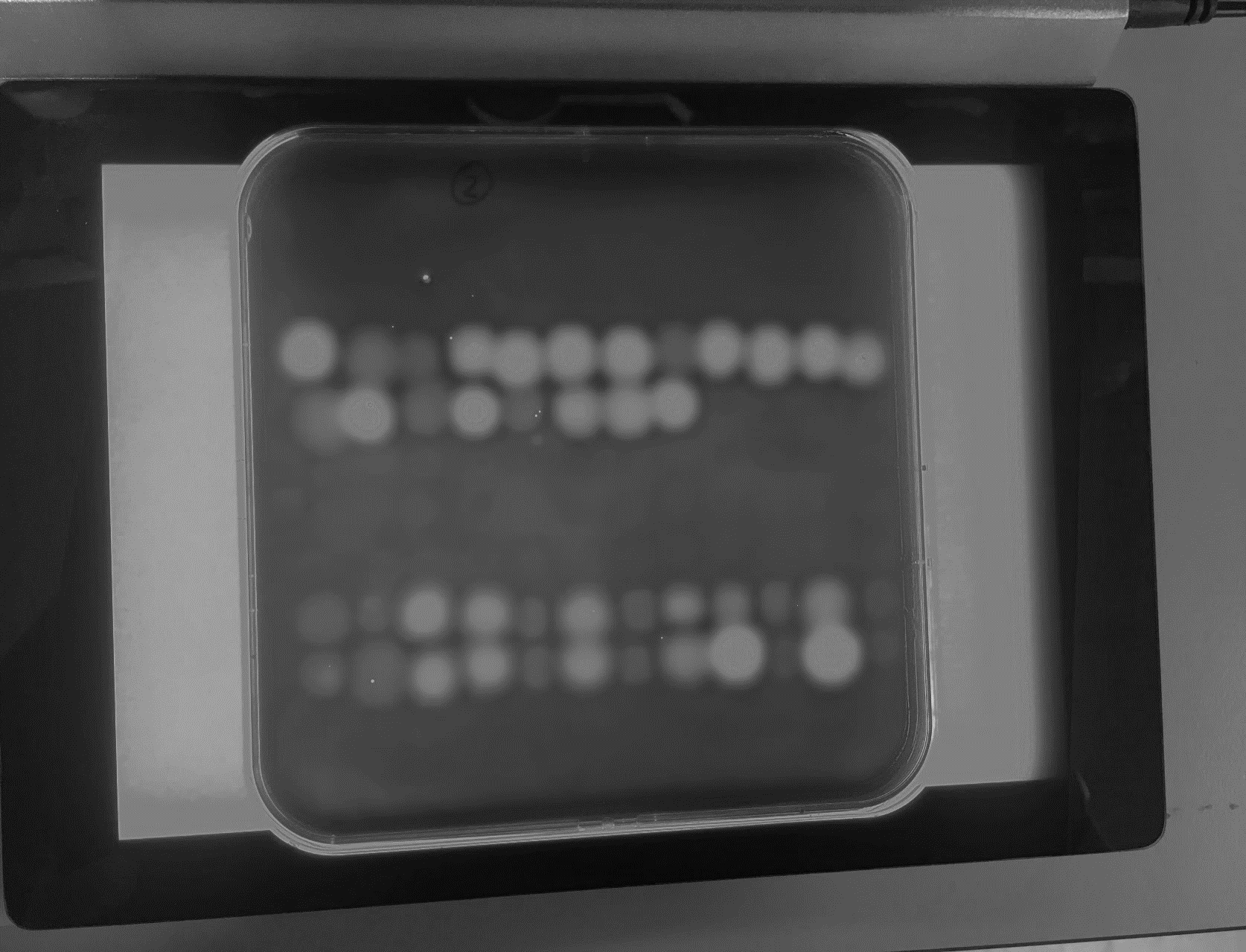

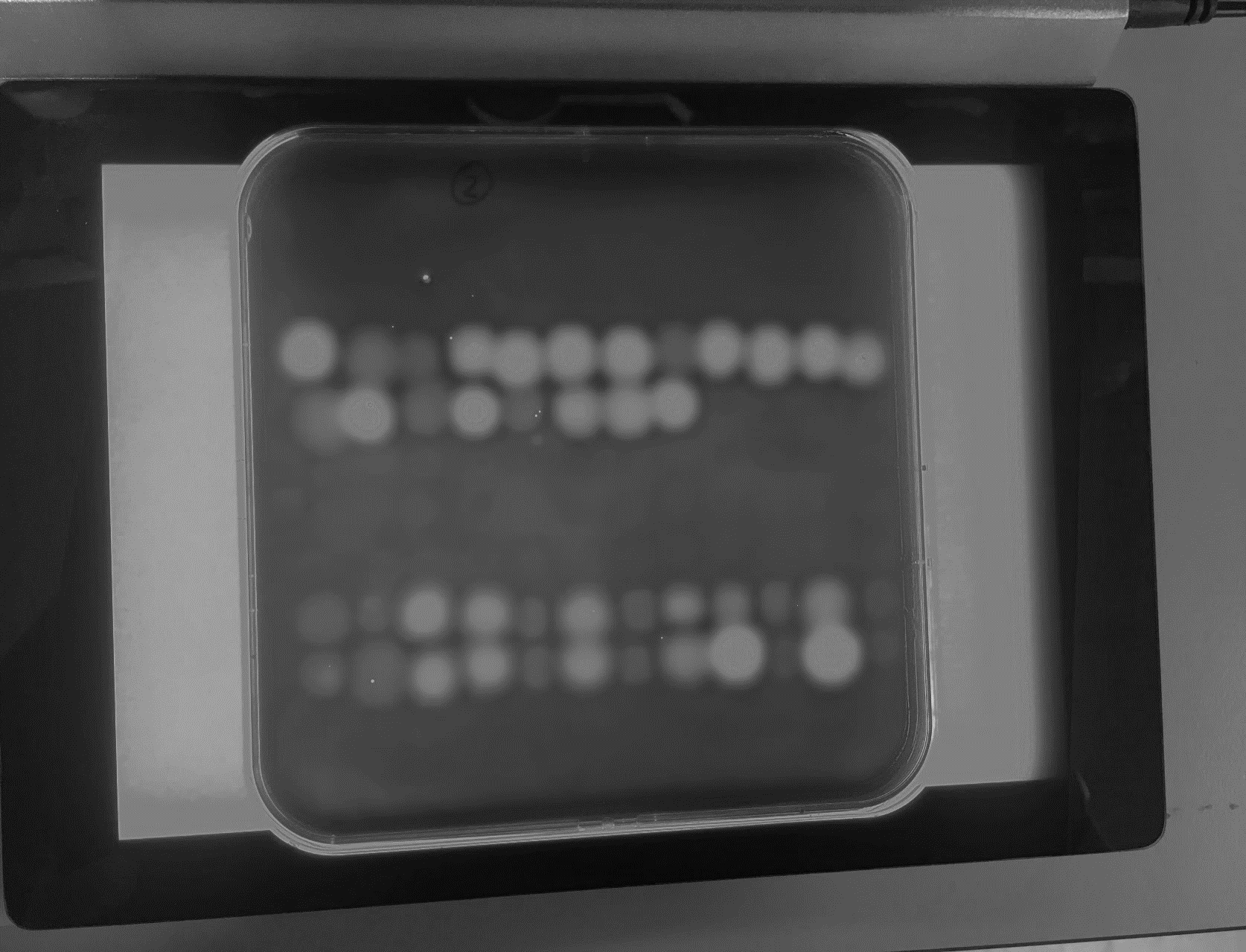

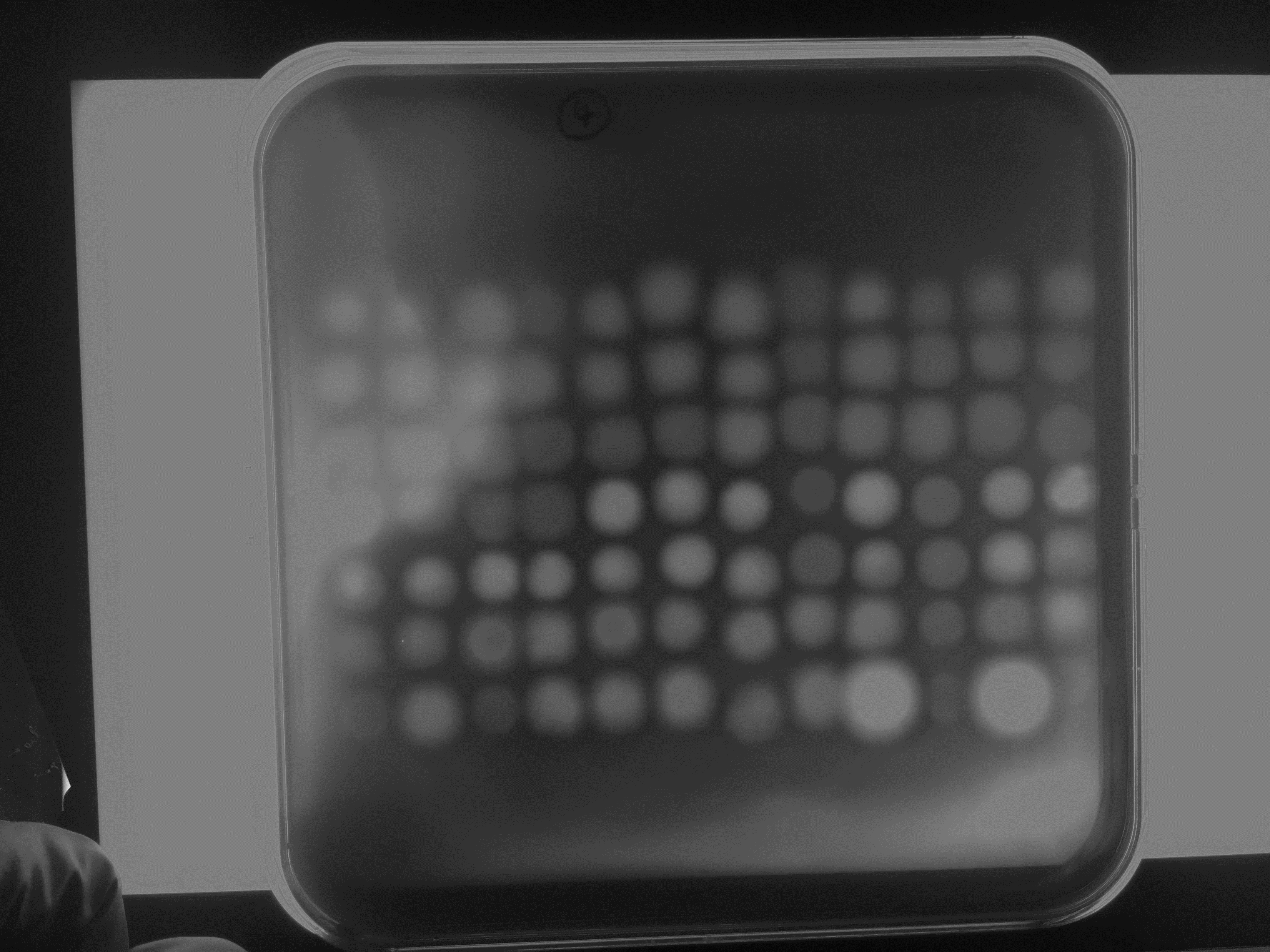

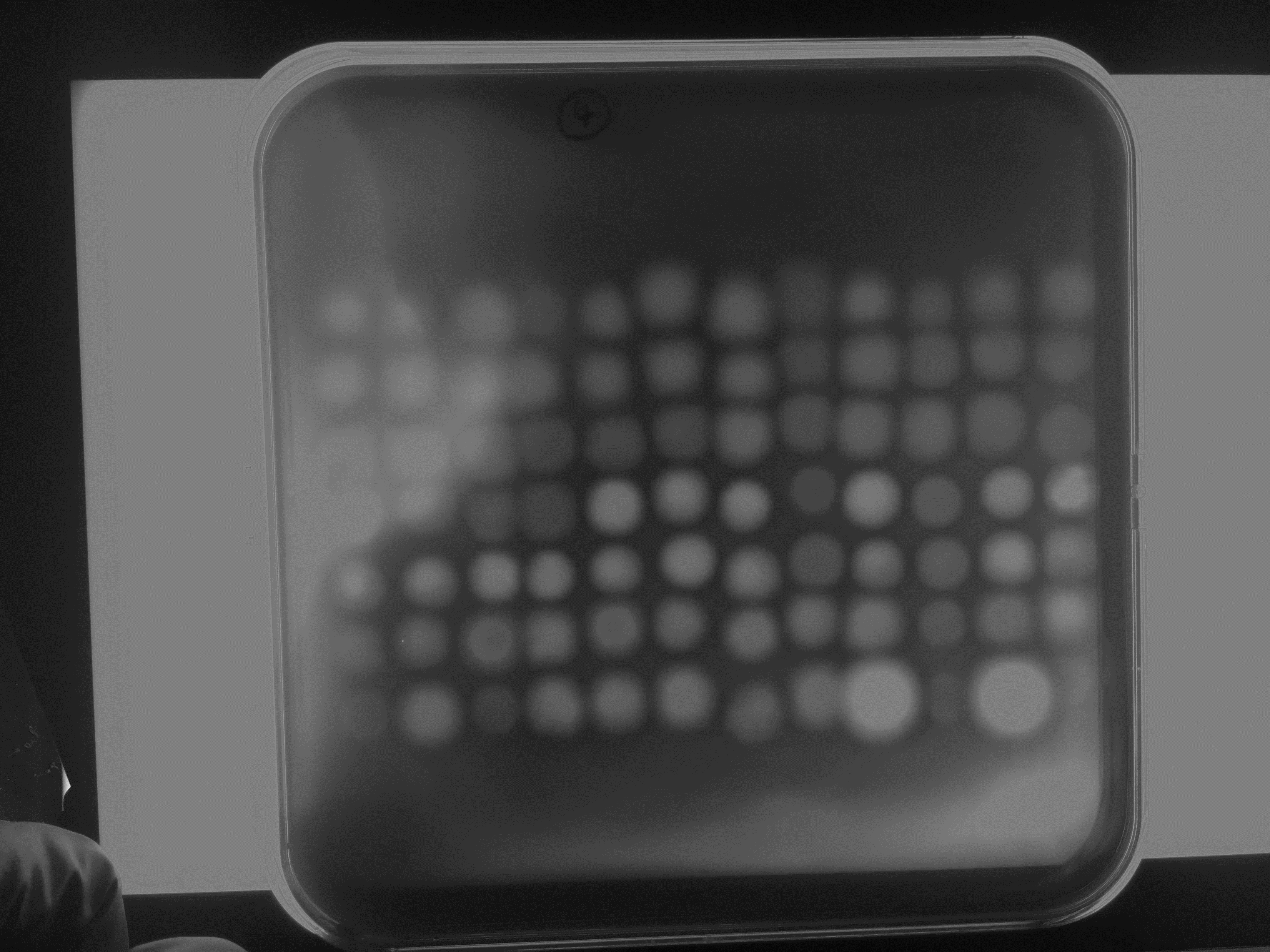

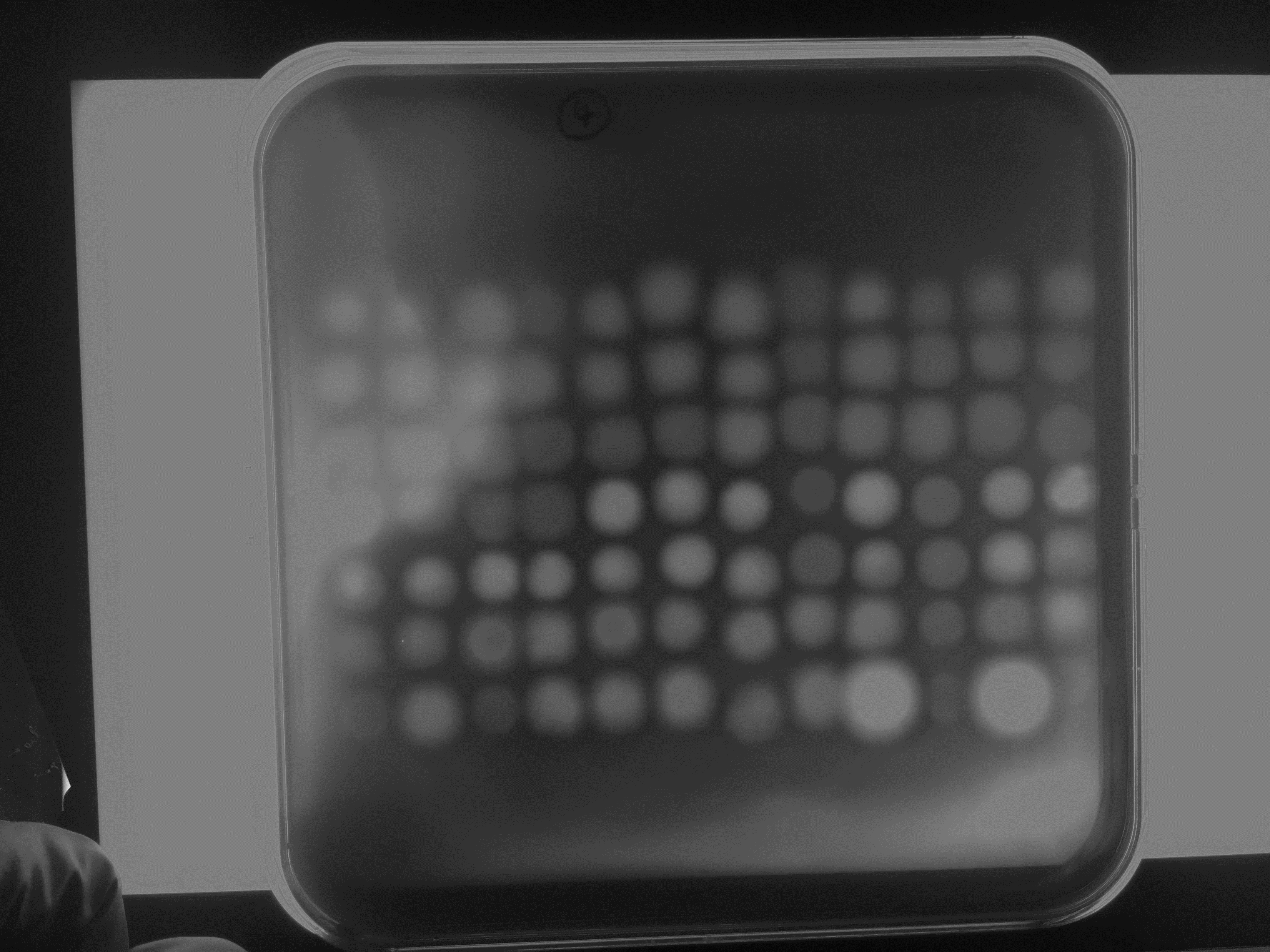

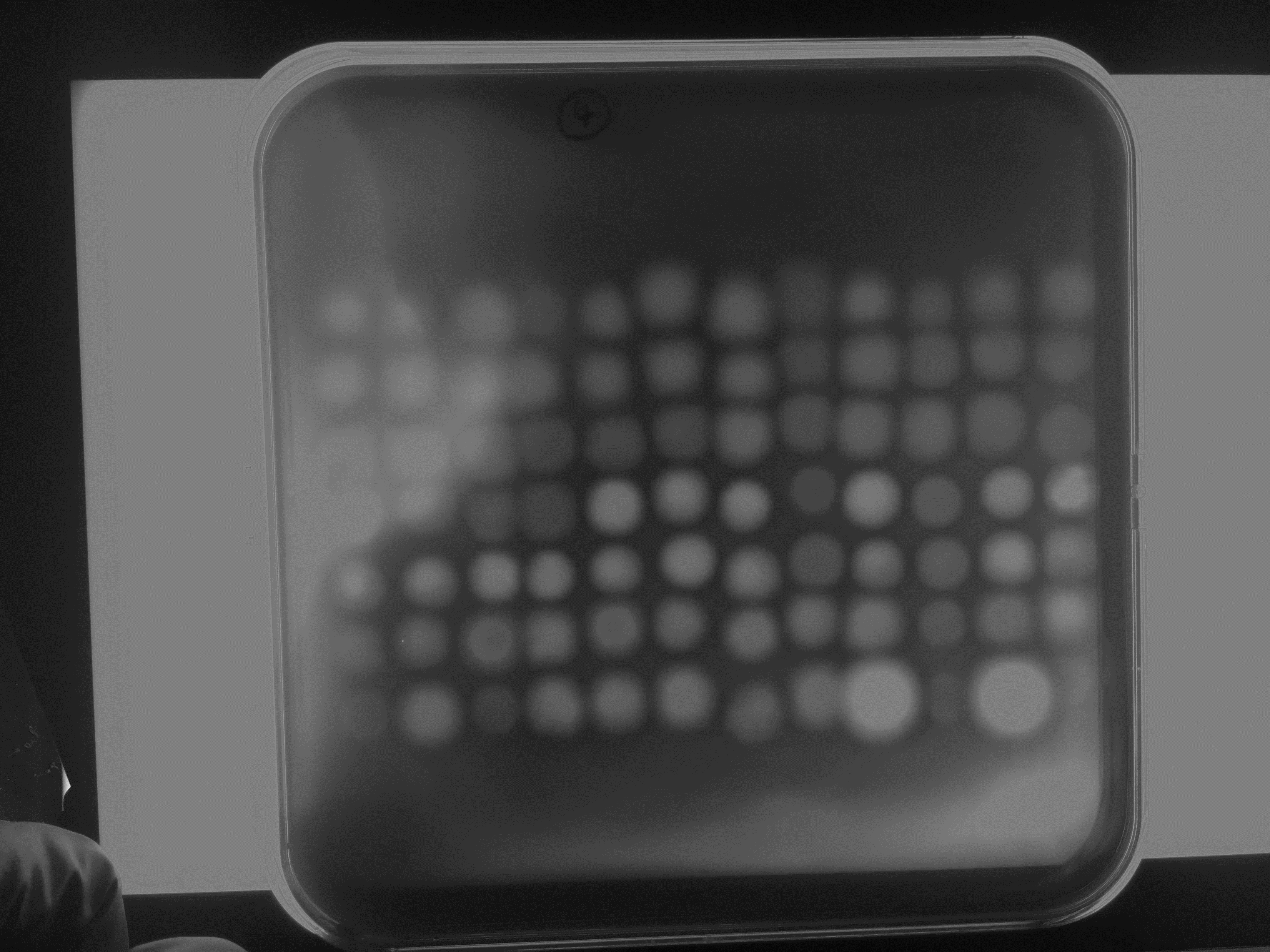

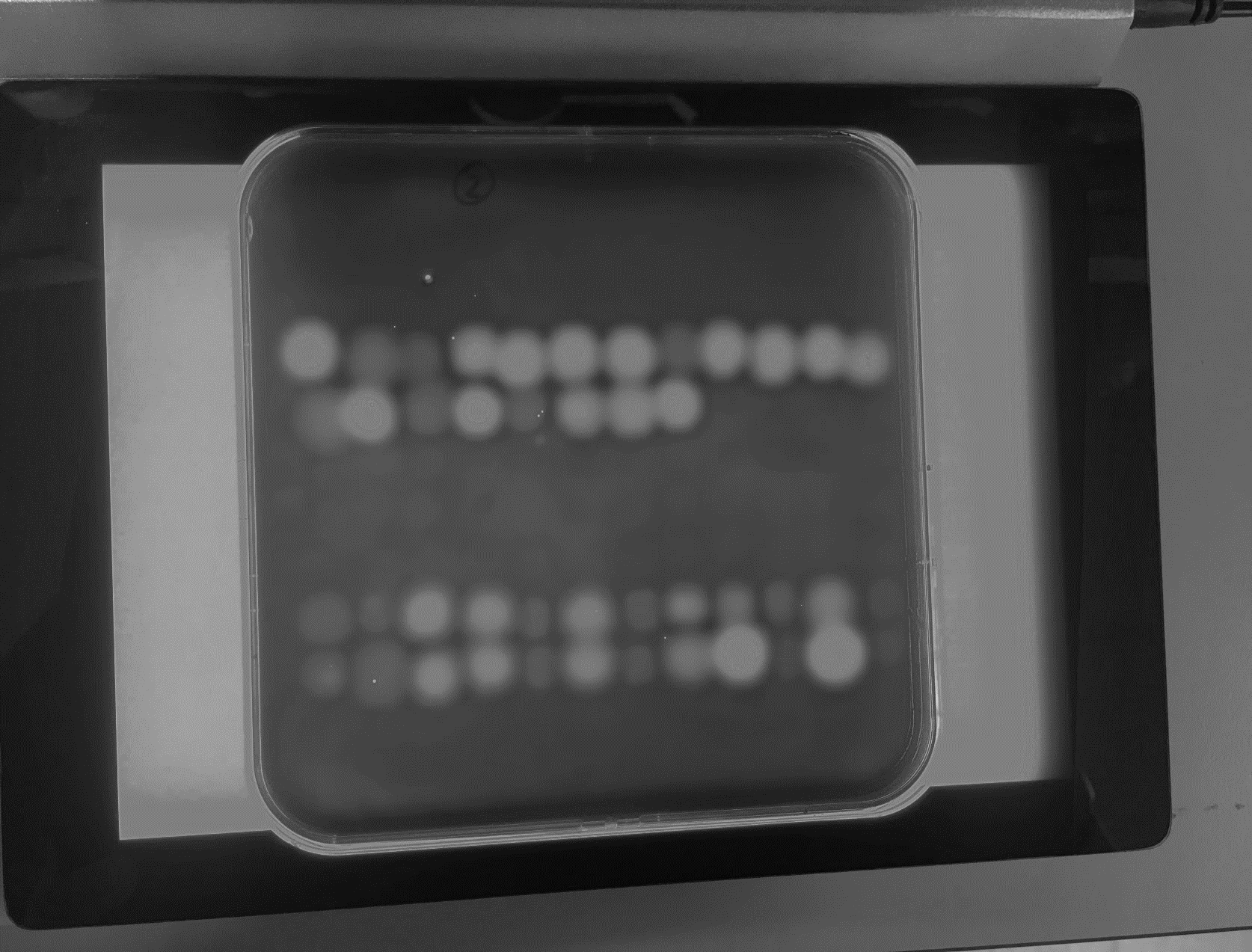

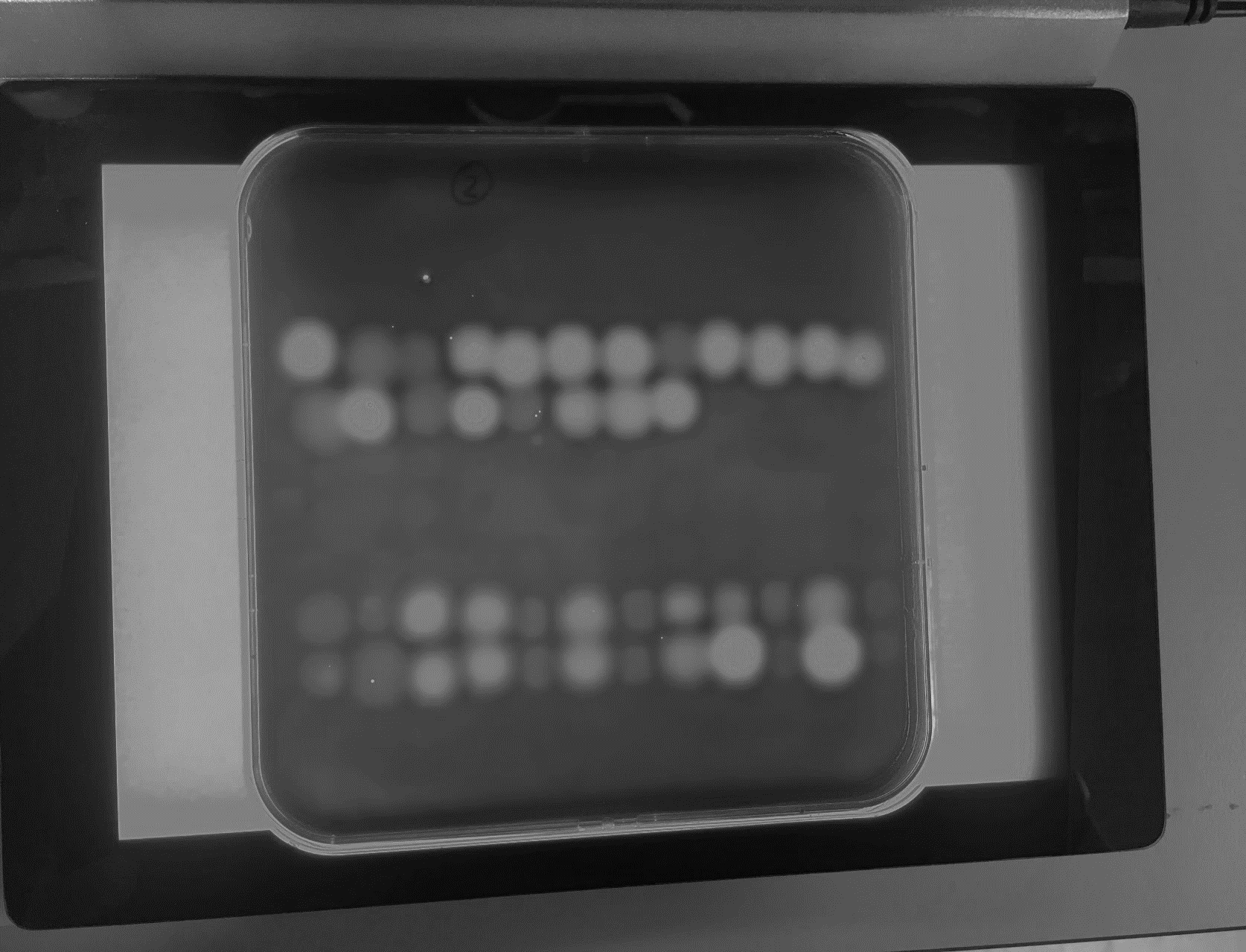

C-terminal extension (1)

C-terminal extension (2)

***P. nicotianae***

***P. cactorum***

***P. nicotianae***

***P. cactorum***

***P. sojae***

***P. sojae***

**Full length**

**Truncated**

**Full length**

**Truncated**

**Full length**

**Truncated**

**Full length**

**Truncated**

**Full length**

**Truncated**

**Full length**

**Truncated**

***P. sojae*_338074**

***P. sojae _*520924**

***P. sojae _*355355**

***P. sojae_*520599**

***P. sojae _*338064**

***P. sojae _*260883**

***P. sojae_*520248**

***P. sojae _*360375**

**Vector only**

***P. sojae_*482953**

***P. sojae_*247788**

***P. sojae_*520924**

***P. sojae_*355355**

***P. sojae_*260883**

***P. sojae_*520599**

***P. sojae_*338074**

***P. sojae_*520248**

***P. sojae_*338064**

**A.**

**B.**

***P. sojae*_559651 (Ps XEG1)**

***P. sojae_*360375 (PsXLP1)**

**Supplemental Figure 5. A.** *P. sojae* xyloglucanase paralogs and orthologs in *P. nicotianae* and *P. cactorum* secreted into *S. cerevisiae* culture supernatants were spotted onto synthetic complete media (minus uracil, for vector maintenance) containing 0.2% (w/v) xyloglucan, and incubated for 24 h at 30°C. Congo red staining of intact xyloglucan allows for detection of halos, where xyloglucan has been digested. Halos were visible after staining in all cases except *P. sojae_*520248, *P. sojae*_338064, *P. sojae*_360375 (PsXLP1), and the vector-only sample (N=3). **B.** *S. cerevisiae* strains expressing GH12 paralogs (i.e. in culture) were spotted onto synthetic complete media (minus uracil, for vector maintenance) containing 0.2% (w/v) xyloglucan, and incubated for 24 h at 30°C. Congo red staining revealed halos for *P. sojae*_520248, suggesting this paralog is active towards xyloglucan using a cell culture-based assay (N=3-4).

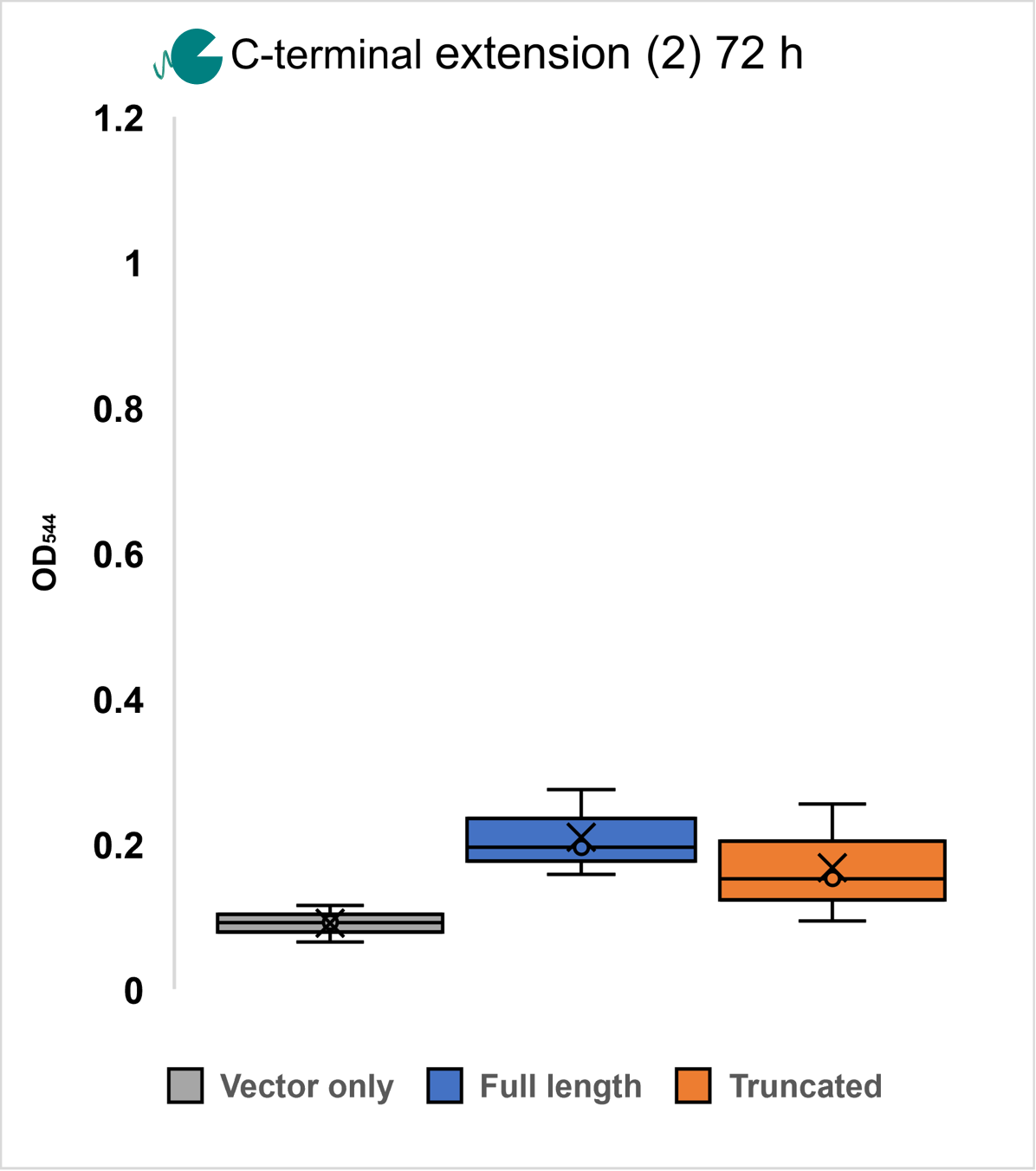

**Supplemental Figure 6 (left).** *P. sojae*_247788 (full-length and truncated) proteins secreted into *S. cerevisiae* culture supernatants were incubated with 1% (w/v) xyloglucan at 30°C, pH7; an increase in absorbance (OD_544_) of DNS reagent added to the samples is suggestive of an increase in the reducing sugars released (i.e. from the breakdown of the substrate). After 72 h of incubation, we did not detect significant differences in released reducing sugars between the *P. sojae*_247788 full-length and truncated proteins (N=3, +/- SD).

***P. sojae*_338064 gene model correction**

Predicted protein sequence pre-correction

>

MKFLLPATVAVAALASSASAQQFCGRDDNKVVGDYTVYNNLWAEENDPKGGQCTFNWTGDSWQVKSFANAALNTAYNYTYDGKIIANVAYDMFTSSTESGDIEYELMVWLAALGDAWPLTSSGQPIKTFTAGRKVKVFTYVAKQQATSFTADLKFFFDQHPTDNNLPTTQFLRKVEAGTEPFQGQNATMVVSSYSVQVK

Corrected DNA sequence (introns highlighted in grey)

>

ATGAAGTTCCTGCTCCCCGCCACCGTCGCCGTCGCCGCTTTGGCCTCGTCCGCCAGTGCTCAGCAGTTCTGCGGTCGCGACGACAACAAGGTGGTCGGAGACTACACCGTGTACAACAACCTGTGGGCCGAGGAAAACGACCCCAAGGGCGGCCAGTGCACCgtaggtgaccggcagcaccagctcgtctgtgtcgtagcagacgagTTCAACTGGACCGGAGACTCGTGGCAGGTCAAGTCGTTCGCCAACGCCGCGCTCAAGacgcccacgcaggtatcggcgatctcttcaatccccacgagCACCGCCTACAATTACACGTACGACGGCAAGATCATCGCCAATGTGGCGTACGACATGTTCACGAGCTCGACCGAGAGCGGTGATATCGAATACGAGCTGATGGTGTGGTTGGCCGCTCTGGGCGACGCTTGGCCTTTGACGTCCAGTGGCCAGCCCATCAAGACCTTCACTGCCGGTggtgtcgagtttaacctgtaccaggactggaacGGAAGGAAGGTCAAGGTCTTCACGTACGTCGCCAAGCAGCAGGCCACGAGCTTCACGGCGGACCTCAAGTTCTTCTTCGACCAGCATCCCACCGACAACAACTTGCCGACGACTCAGTTTCTGAGGAAGGTGGAGGCCGGCACGGAGCCGTTCCAGGGCCAGAACGCCACGATGGTCGTGTCGTCCTACTCAGTGCAGGTCAAGTAA

Corrected protein sequence (used in this study)

>

MKFLLPATVAVAALASSASAQQFCGRDDNKVVGDYTVYNNLWAEENDPKGGQCTVGDRQHQLVCVVADEFNWTGDSWQVKSFANAALKTPTQVSAISSIPTSTAYNYTYDGKIIANVAYDMFTSSTESGDIEYELMVWLAALGDAWPLTSSGQPIKTFTAGGVEFNLYQDWNGRKVKVFTYVAKQQATSFTADLKFFFDQHPTDNNLPTTQFLRKVEAGTEPFQGQNATMVVSSYSVQVK

**Supplemental Figure 7.** The gene encoding *P. sojae*_338064 was predicted to encode three introns (highlighted in grey), which were not supported by available transcriptome data (FungiDB; (50, 51)), therefore the sequence was subject to manual correction before use in this study.

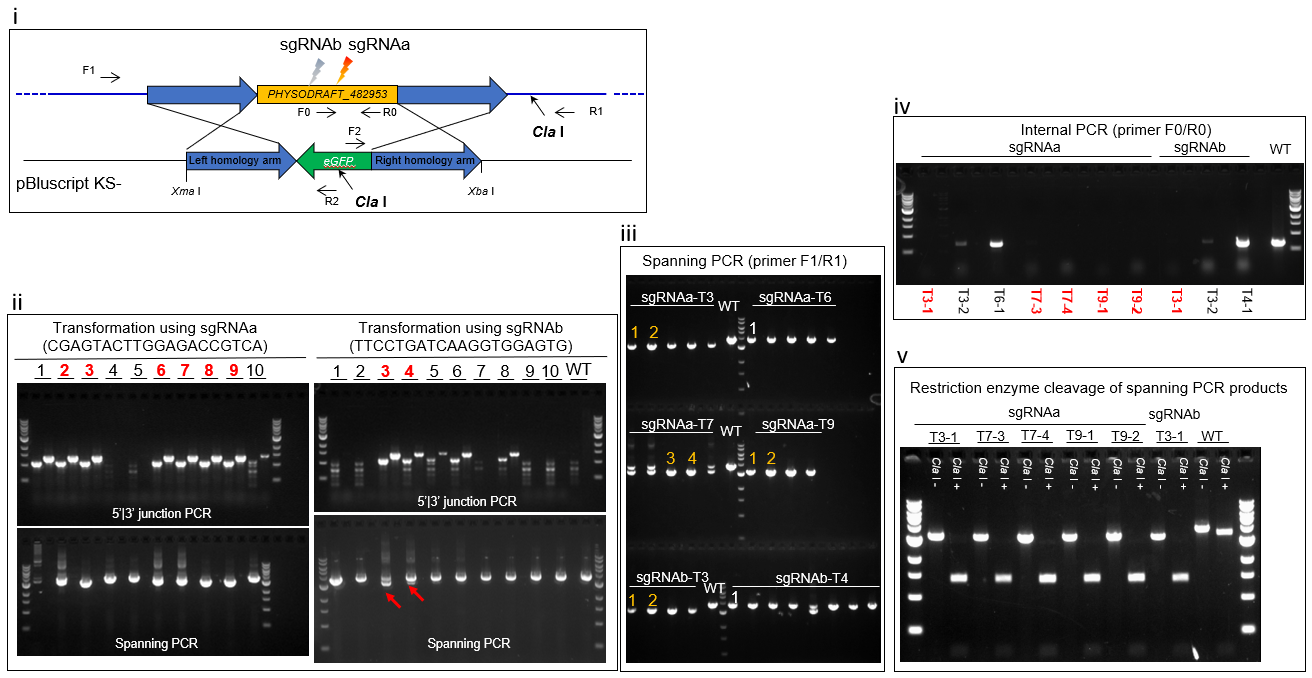

**Supplemental Figure 8.** Genotyping results of primary and zoospore-isolated *P. sojae* transformants. **(i)** Schematic of HDR-mediated modification of the target gene *P. sojae*_482953 (PHYSODRAFT_482953), stimulated by Cas9/sgRNA-induced double-strand break. **(ii)** Two sgRNAs were used for two independent *P. sojae* transformations (see Supplemental Table 1 for sequences). 10 transformants of each transformation were arbitrarily picked for gDNA miniprep and genotyping. For 5’-|3’-junction PCR results, the first well of each transformant shows the PCR result of 5’ junction, while the second is from 3’-junction. Transformants showing correct PCR amplication (5’- and 3’-junction PCR as well as spanning PCR) are labelled in red. These transformants were passaged to V8 agar before zoospore isolation as per Fang et al. (2017) (58). The spanning PCR of some transformants showed two bands (shown by the red arrows), indicating heterozygosity or mixture of wild-type and mutants – these were also picked for homokaryonization (passage to V8 agar supplemented with G418). **(iii)** 4-8 zoospore-germinated mycelial colonies were randomly picked for gDNA miniprep and genotyping. Representative mycelial colonies of each transformant that showed correct sizes (labelled in yellow), and incorrect sizes (labelled in white) of spanning PCR were selected for further verification in iv and v. **(iv)** Verification of selected mycelial colonies using internal PCR (correct colonies are labelled in red and bold). **(v)** Spanning PCR of mycelial colonies that showed correct internal PCR were further checked by restriction digests. Spanning PCR products of all selected transformants were cleaved by *ClaI* efficiently, indicating true homozygous mutants. Mutants selected for phenotyping: sgRNAa T3-1, sgRNAa T9-1, and sgRNAa T9-2.

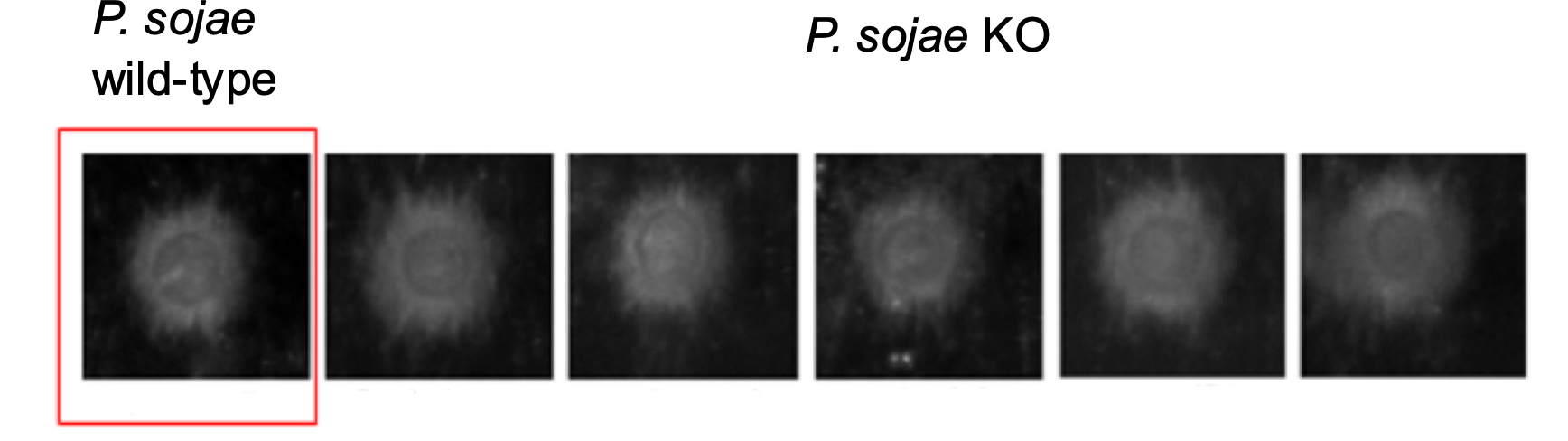

**A.**

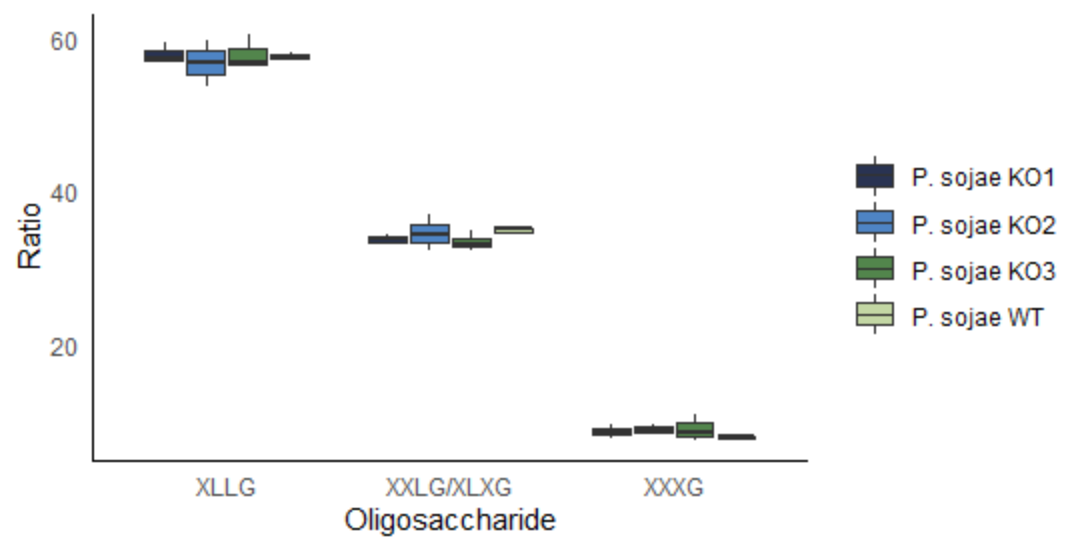

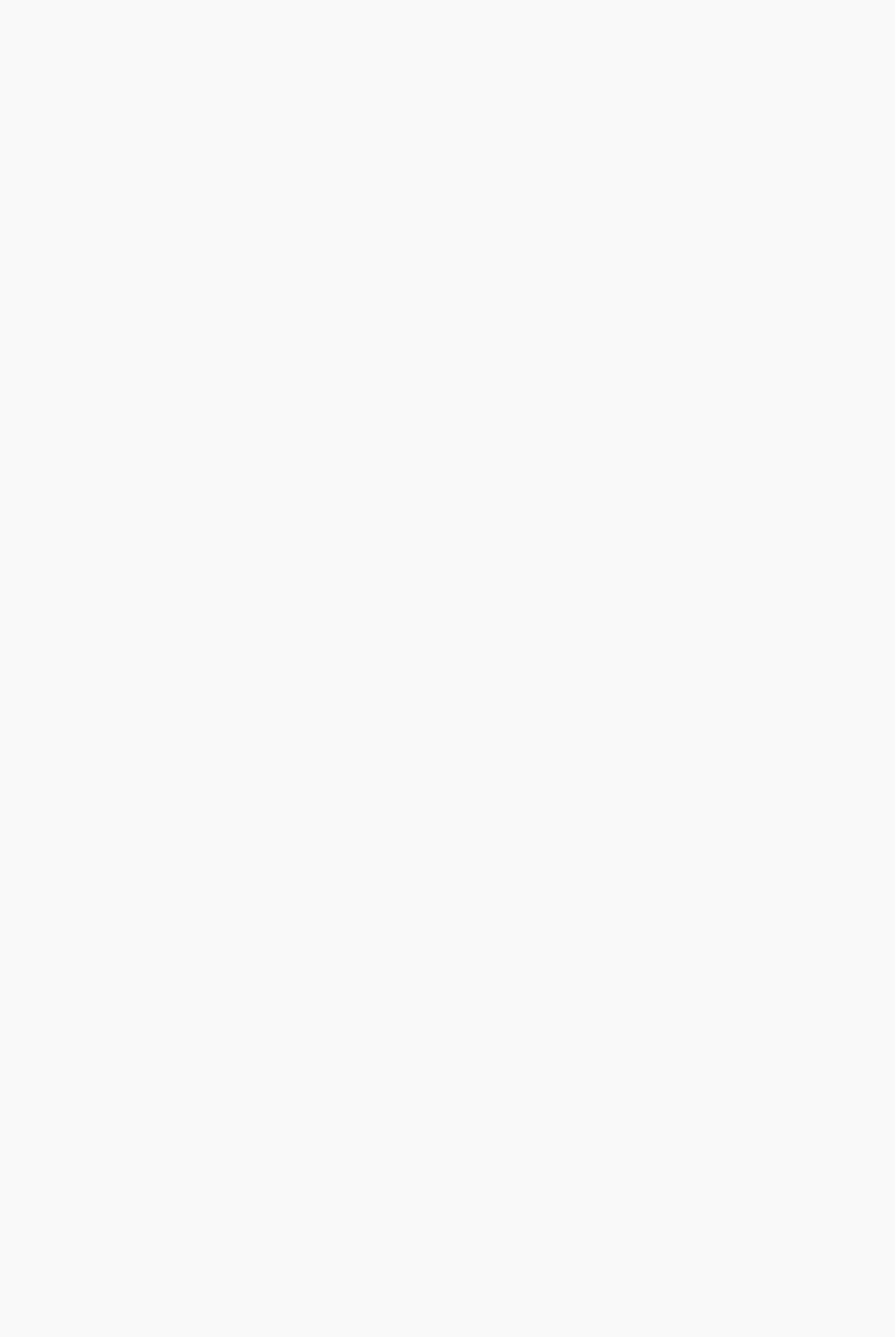

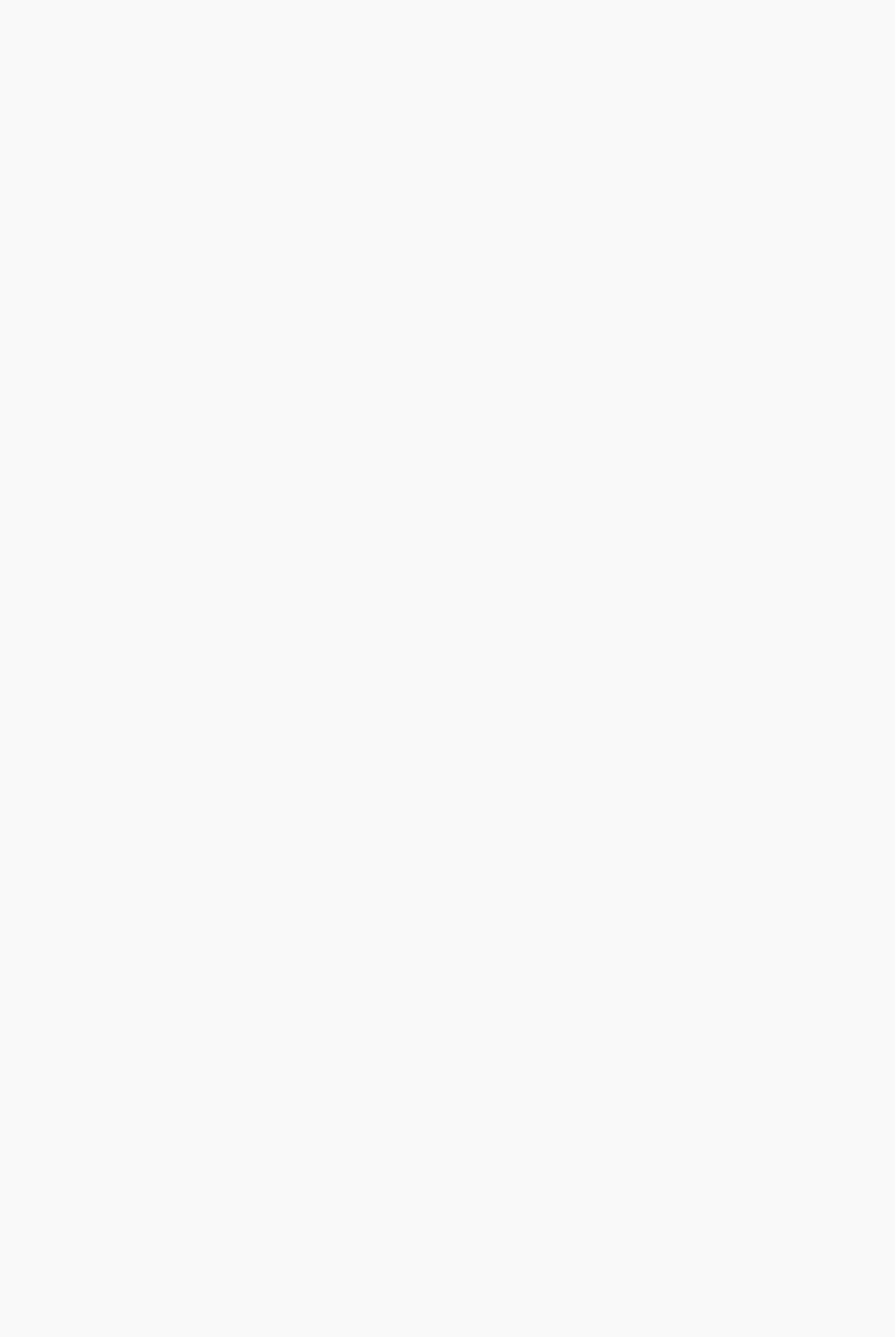

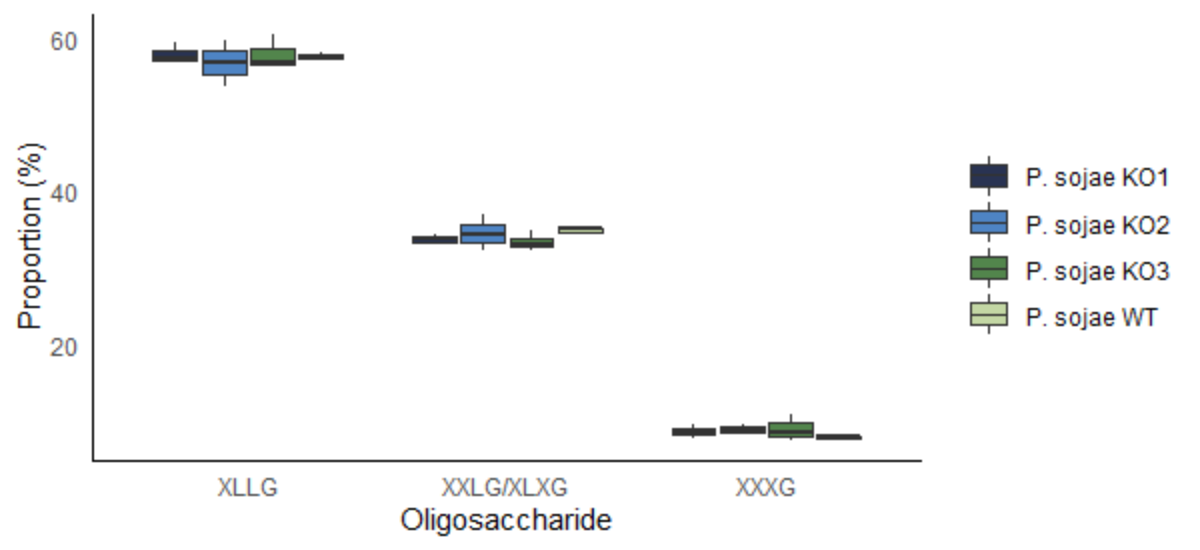

**B.**

**Supplemental Figure 9. A.** Three independent *P. sojae* mutants with verified deletion of the gene encoding *P. sojae*_482953 were tested for their ability to utilise xyloglucan as a sole carbon source, by incubating hyphal plugs on minimal agar containing 1% (w/v) xyloglucan at 25°C (for 7 days, in the dark), after which colony morphology pictures were taken. Phenotypic analysis of the gene deletion strains (labelled ‘*P. sojae* KO’ with 1-2 replicates for each *P. sojae* mutant) indicated that there was no significant effect to growth on xyloglucan as the sole carbon source, based on comparisons with the wild-type (WT) strain (i.e. there were no differences in the diameters of mycelial growth between isolates that would suggest an altered capacity to utilise xyloglucan resulting from the gene deletion). **B.** Hyphal plugs of *P. sojae* WT and three independent gene deletion mutants were incubated in liquid minimal media with 1% (w/v) xyloglucan (for 7 days, in the dark). MALDI-MS spectra confirmed the release of xyloglucan oligosaccharides into the culture media. Three peaks of interest were observed; ions with *m/z* of ~1085, 1247, and 1409 – putatively corresponding to the oligosaccharides XXXG, XXLG (or XLXG), and XLLG, respectively (24). The relative intensity of species identified at these peaks were compared by calculating the ratios between the areas under the peaks, to probe putative differences in preferential binding of the xyloglucan backbone. We find no evidence for differences in oligosaccharides released between the WT and gene deletion strains (labelled ‘*P. sojae* KO1-KO3').

| **Name** | **Sequence** | **Use in this study** |
| --- | --- | --- |
| JOHE50046/YF/VA | CTAGCCAGGAACTGATGAGTCCGTGAGGACGAAACGAGTAAGCTCGTCTTCCTGATCAAGGTGGAGTG | sgRNA version a. The 20 nt sgRNA sequence is highlighted |
| JOHE50047/YF/VA | AAACCACTCCACCTTGATCAGGAAGACGAGCTTACTCGTTTCGTCCTCACGGACTCATCAGTTCCTGG |  |
| JOHE50048/YF/VA | CTAGCTACTCGCTGATGAGTCCGTGAGGACGAAACGAGTAAGCTCGTCCGAGTACTTGGAGACCGTCA | sgRNA version b. The 20 nt sgRNA sequence is highlighted |
| JOHE50049/YF/VA | AAACAGACGGTCTCCAACTACTCGGACGAGCTTACTCGTTTCGTCCTCACGGACTCATCAGCGAGTAG |  |
| pBSKSM_LA_Fw | gagctccaccgcggtggcggccgctctagaCTGTAAGATCCGTTTGGC | HDR template (‘left arm’) – eGFP sequence in red |
| eGFP_LA_Rv | cgcccttgcccatGCTTGGTCGAGTTGGATG |  |
| LA_eGFP_Fw | aactcgaccaagcATGGGCAAGGGCGAGGAA | HDR template (‘eGFP’) – eGFP sequence in red |
| RA_eGFP_Rv | atgaaatccacatCTACTTGTAGAGTTCATCCATGCCATG |  |
| eGFP_RA_Fw | actctacaagtagATGTGGATTTCATTTAGATGAACTG | HDR template (‘right arm’) – eGFP sequence in red |
| pBSKSM_RA_Rv | aagcttgatatcgaattcctgcagcccgggTCGAGTCAGCGTGTGTAAC |  |
| JOHE50375 | TTCTCGTCACGGCGGAAAACA | Genotyping, 5'_junction_PCR, (Supplemental Figure 5) |
| JOHE45529 | CCATGCCCGAAGGCTACG |  |
| JOHE45530 | TTGATGCCGTTCTTCTGCTTGTC | Genotyping, 3'_junction_PCR, (Supplemental Figure 5) |
| JOHE50376 | CGGCTCAGCAGTATCGCAACC |  |
| JOHE50477 | GCCGAGTACTTGGAGACCGTCATG | Genotyping, internal PCR (Supplemental Figure 5) |
| JOHE50478 | GAAGGTTCTGTTCGCCACTGCTCTG |  |

**Supplemental Table 1.** Primers and sgRNA used to generate the *P. sojae* gene deletion mutant.
